# Expansion-Assisted Iterative-FISH defines lateral hypothalamus spatio-molecular organization

**DOI:** 10.1101/2021.03.08.434304

**Authors:** Yuhan Wang, Mark Eddison, Greg Fleishman, Martin Weigert, Shengjin Xu, Fredrick E. Henry, Tim Wang, Andrew L. Lemire, Uwe Schmidt, Hui Yang, Konrad Rokicki, Cristian Goina, Karel Svoboda, Eugene W. Myers, Stephan Saalfeld, Wyatt Korff, Scott M. Sternson, Paul W. Tillberg

**Author notes:** These authors jointly supervised this work (Scott M. Sternson, Paul W. Tillberg). Corresponding authors. (S.M.S.); (P.W.T.). Innate Pharma, 2273 Research Boulevard, Suite 350, Rockville, MD 20850, USA.

## Abstract

Determining the spatial organization and morphological characteristics of molecularly defined cell types is a major bottleneck for characterizing the architecture underpinning brain function. We developed <u>E</u>xpansion-<u>As</u>sisted Iterative <u>F</u>luorescence *In <u>S</u>itu* <u>H</u>ybridization (EASI-FISH) to survey gene expression in brain tissue, as well as a turnkey computational pipeline to rapidly process large EASI-FISH image datasets. EASI-FISH was optimized for thick brain sections (300 µm) to facilitate reconstruction of spatio-molecular domains that generalize across brains. Using the EASI-FISH pipeline, we investigated the spatial distribution of dozens of molecularly defined cell types in the lateral hypothalamic area (LHA), a brain region with poorly defined anatomical organization. Mapping cell types in the LHA revealed nine novel spatially and molecularly defined subregions. EASI-FISH also facilitates iterative re-analysis of scRNA-Seq datasets to determine marker-genes that further dissociated spatial and morphological heterogeneity. The EASI-FISH pipeline democratizes mapping molecularly defined cell types, enabling discoveries about brain organization.

**Highlights:**

- EASI-FISH enables robust gene expression profiling in thick brain slices
- A turnkey analysis pipeline for facile analysis of large EASI-FISH image datasets
- EASI-FISH reveals novel subregions of the lateral hypothalamus
- Identification of rare cell types based on morphological and spatial heterogeneity

## Introduction

Neuronal diversity is a major contributor to brain functions (Zeng and Sanes, 2017). Neuronal subtypes (cell types) have been defined based on morphology, connectivity, gene expression, electrical properties, and selective functional-response-tuning. The recent emergence of low-cost, high-throughput single-cell RNA-Sequencing technology (scRNA-Seq) has enabled systematic identification of new cell types in the brain (Saunders et al., 2018; Tasic et al., 2018; Zeisel et al., 2018). Molecular definition of cell types provides a way to classify neurons and survey neuronal heterogeneity. In addition, marker-genes can be leveraged to selectively interrogate the function of specific cell types in neural circuits (Luo et al., 2018).

Experimental methods and computational analysis pipelines have been developed that make scRNA-Seq a ubiquitous and indispensable technique in neuroscience research (Butler et al., 2018). However, establishing the spatial organization of cell types predicted by scRNA-Seq analysis requires mapping the co-expression patterns of dozens of genes in the same cells in three-dimensional (3D) tissue volumes. For widespread usage, methods for spatial analysis of gene expression should address several key issues. 1) *In situ* measurement of gene expression should be sensitive, quantitative, and consistent with measurements made by scRNA-Seq. 2) Gene expression measurement in tissue samples should be stable and robust to normal storage and experimental manipulation. 3) Measurement of gene expression should use widely available laboratory equipment. 4) Computational analysis of spatial gene expression should be automated. 5) Transcripts must be accurately assigned to individual cells. 6) Gene expression analysis should be performed in thick tissues to enable 3D mapping of cell types, which will facilitate alignment and detailed comparison of samples from different brains and is important for establishing generalizable molecular and structural features.

Fluorescent *in situ* hybridization (FISH) is especially well-suited to satisfy these criteria. Although multiple methods have been reported (Chen et al., 2015b; Codeluppi et al., 2018; Moffitt et al., 2018; Nicovich et al., 2019; Qian et al., 2020; Shah et al., 2016; Wang et al., 2018), it remains a bottleneck for the vast majority of research labs due to requirements for specialized equipment, complex procedures, and challenging multi-step computational analyses. Most methods suitable for more than 10 marker-genes require complex experimental setups and barcoding/decoding schemes (Chen et al., 2015b; Shah et al., 2016). Attempts to amplify *in situ* RNA by conversion to cDNA also have limited efficiency and may result in nonlinear amplification (Ke et al., 2013; Wang et al., 2018). Additionally, most methods suffer from optical crowding, which limits accurate RNA quantification, and are restricted to single-cell-layer tissue sections (10-20 µm), obscuring three-dimensional (3D) relationships of cell types to brain structure (Hashikawa et al., 2020).

To overcome these limitations, we developed <u>E</u>xpansion-<u>As</u>sisted Iterative-FISH (EASI-FISH) with multi-round multiplexed RNA-FISH in 300 µm thick brain sections. Expansion microscopy (ExM) (Chen et al., 2015a; Tillberg et al., 2016) is advantageous for high-resolution imaging in thick tissue. Imaging thick tissues also enables dozens of cell layers to be included in a single sample, allowing 3D reconstruction of tissue volumes. This tissue thickness is also well-suited for *post hoc* analysis of tissues obtained from other experimental modalities, such as brain slice recordings and *in vivo* imaging. We also developed a turnkey analysis pipeline that allows for rapid and automated data processing, facilitating adoption of high-plex FISH in thick tissue as a routine laboratory method for tissue analysis.

We applied EASI-FISH to the mouse lateral hypothalamus (LHA), a brain region that has been studied for decades as an important motivational center regulating ingestive, social, arousal, and autonomic functions (Bernardis and Bellinger, 1993; Petrovich, 2018; Stuber and Wise, 2016). Despite extensive functional investigation, the understanding of the LHA is limited by poor anatomical definition. Some LHA parcellations, based on differences in cellular density or axonal projections, have been reported in rat (Geeraedts et al., 1990; Veening et al., 1987), but these have not been adopted for segregation of LHA cell types. Here, we performed EASI-FISH using molecularly defined cell type markers identified from LHA scRNA-Seq datasets and uncovered an unexpected parcellation of the LHA not previously predicted from cell density measurements. This includes unreported laminar structures to which different cell types segregate. EASI-FISH also reveals morphological diversity of neuronal types in this region. Iterative use of EASI-FISH and scRNA-Seq allowed us to further subdivide spatially separated members of a transcriptional cluster with distinct marker-genes. Taken together, our results demonstrate the capability of the EASI-FISH data acquisition and analysis pipeline, in combination with scRNA-Seq data, to readily access an unprecedented view of the organization of cell types in the LHA that underlies the multifaceted functions of this important brain area.

## Results

### EASI-FISH protocol development

We designed and implemented EASI-FISH (Figure 1A) in thick tissue sections from cortex, central amygdala (CEA), and LHA by building on expansion microscopy enhanced smFISH (exFISH) (Chen et al., 2016), where tissue is physically expanded by embedding in a swellable hydrogel (Chen et al., 2015a; Tillberg et al., 2016). Uniform expansion is achieved by proteolytic digestion of the embedded tissue, while preserving RNA via covalent attachment to the hydrogel mesh. Anchored mRNAs are available for detection by FISH methods, including amplification with the hybridization chain reaction (HCR) (Figure 1A and 1B). Proteolytic digestion and volumetric expansion (2× linear expansion) reduces tissue autofluorescence (95% reduction) and light scattering, yielding a composite material that is refractive index-matched for water immersion objective lenses. Expansion also increases the effective imaging resolution, increasing the number of individual RNA molecules that may be resolved per cell (Figure 1C).

**Figure 1.**
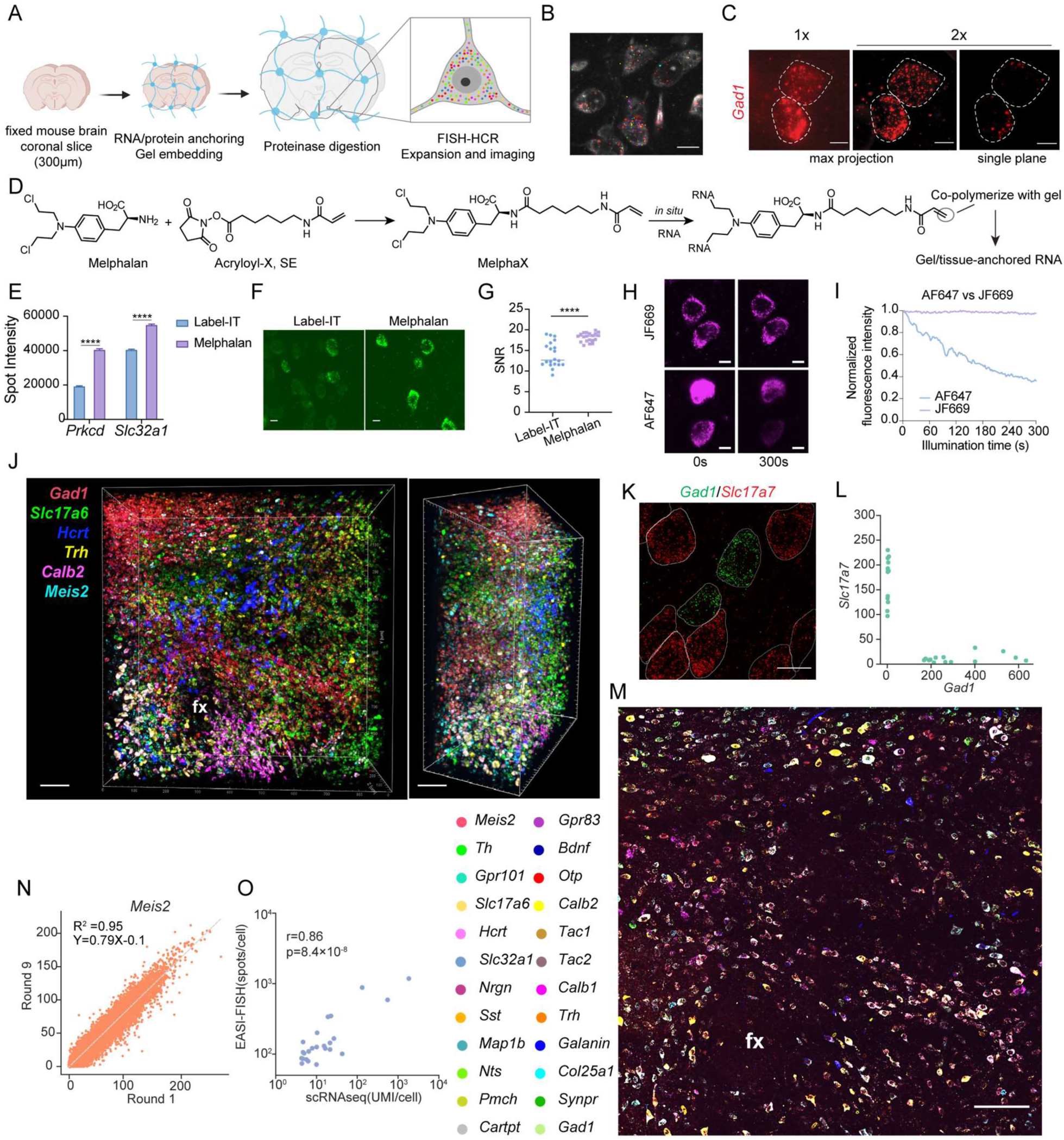
EASI-FISH method. **(A)** Schematic of the EASI-FISH platform. (**B**) Example image of gene expression detected by EASI-FISH. Scale bar: 10 µm. (**C**) Axial projection and single-plane images of *Gad1* expression in two cortical neurons before (1× expansion factor) and after expansion (2× expansion factor). Scale bar: 5 µm (pre-expansion length). (**D**) Chemical structure of Melphalan and chemical reaction with Acryloyl-X, SE to produce MelphaX, which reacts with mRNA and can be incorporated into a hydrogel matrix. (**E**) *Prkcd* and *Slc32a1* spot fluorescence intensity with RNA anchoring by Label-IT (0.1mg/ml) or Melphalan (0.1mg/ml). (**F**) Representative images and (**G**) quantification of signal-to-noise ratio (SNR) of images where *Gad1* mRNA was detected in cortex using different RNA anchoring methods. Scale bar: 5µm. The SNR was defined as the ratio of mean pixel value to the standard deviation of pixel values. (**H**) Representative images and (**I**) quantification showing photostability comparison between hairpins conjugated with Alexa-fluor 647 (AF-647) and Janelia-Fluor 669 (JF-669). Scale bar: 5 µm. (**J**) 3D rendering of a thick tissue volume generated by EASI-FISH with 2 rounds of 3-plex FISH. Scale bar:100 µm. fx: fornix. (**K**) Representative image and (**L**) quantification of *Slc17a7* and *Gad1* expression in the cortex. Scale bar: 10 µm. (**M**) Representative image from 8 rounds of 3-plex EASI-FISH (aligned) in one LHA tissue volume with 24 marker-genes (single optical plane shown). Imaged tissue dimension: 0.8 mm × 0.8 mm × 0.3 mm before expansion. Scale bar:100 µm. fx: fornix. (**N**) Spot count with EASI-FISH in round 1 and round 9 for the same gene (*Meis2*). (**O**) Correlation analysis for 24 marker-genes between EASI-FISH spot counts and scRNA-Seq UMIs from the LHA. **** p < 0.0001. Error bars: SEM. Statistics: **Table S1**.

We optimized the exFISH procedure for improved detection accuracy and robust sample processing across multiple rounds. First, to covalently anchor RNA molecules to the hydrogel, we used the bis-nitrogen mustard, Melphalan, instead of the exFISH linker, Label-IT. Like Label-IT, Melphalan is an alkylating agent with a primary amine available for conjugation to NHS esters, but it is widely available through major chemical vendors, 50 times less expensive than Label-IT, and has two alkylating moieties per molecule. After reacting Melphalan with the succinimidyl ester of 6-((Acryloyl)amino)hexanoic Acid (Acryloyl-X, SE), the product (MelphaX) is applied to tissue and reacts with nucleotides, functionalizing them for incorporation into a tissue-gel network by polymerization of the acryloyl “tails” (Figure 1D). Importantly, RNA retention with Melphalan was comparable to Label-IT, as measured by spot count (Figure S1A). Furthermore, Melphalan significantly increased the brightness of individual spots (Figure 1E) and reduced the autofluorescence background compared with Label-IT (Figure 1F). This improved the signal-to-noise ratio by 25%, which increases spot detection sensitivity (Figure 1G).

Tissue clearing and isotropic expansion depends on protein digestion. We optimized this step for thick tissue by inclusion of the ionic detergent, sodium dodecylsulfate, in the protease digestion as well as by increasing tissue expansion in the digestion step through reduced salt concentration. This led to greatly improved and rapid optical clearing and reagent penetration through 300-µm-thick tissue volumes compared to the original exFISH protocol (Chen et al., 2016) (Figure S1B).

For amplification of FISH signal, we chose to use the hybridization chain reaction (HCR) (Dirks and Pierce, 2004) as the probe and amplification oligos are short (50-100nt) and can therefore rapidly penetrate into thick tissue. In contrast, another FISH signal amplification method, RNAscope (Wang et al., 2012), did not show sufficient reagent penetration in thick tissues (Figure S1C-D). We adopted HCR v3.0 (Choi et al., 2018), which requires binding of two adjacent probes for signal amplification, thereby reducing non-specific spots. We also optimized the hybridization conditions (probe concentration, wash conditions, etc.) to improve detection specificity (see Methods and Figure S1E).

Selective plane illumination fluorescence microscopy (SPIM or ‘light sheet microscopy’) provides a decisive advantage for rapidly imaging in large tissue samples. SPIM avoids photobleaching of out-of-focus fluorophores and accelerates image acquisition ∼100-fold compared with confocal microscopy. Image acquisition in EASI-FISH samples with single transcript sensitivity following HCR amplification was readily performed with a turnkey SPIM microscope (Zeiss Z.1 microscope, also see Methods).

We found that HCR spots were susceptible to light-induced fragmentation, producing mobile spots, which reduced spot detection fidelity. Fragmentation could be reduced to acceptable levels either by reducing laser intensity or by including anti-fade compounds such as p-phenylenediamine (PPD), though the latter method reduced the initial brightness of AlexaFluor 546 to an unacceptable level (see Methods, **Supplemental Information movie 1**, **Table S2** and Figure S1H for details). We observed fast fluorophore photobleaching using commercial HCR probes conjugated with AlexaFluor-647. To improve photostability, we custom-labelled the corresponding HCR amplification hairpins with the photostable far-red dye JF-669 (Grimm et al., 2017) (Figure 1H and 1I).

For most neuroscience applications, dozens of marker-genes are sufficient to map the distribution of neuronal cell types. In our approach, three transcript species were detected in each imaging round, compatible with the spectral capabilities of most standard fluorescence microscopes. Each round of FISH can be analyzed independently and thus allows flexibility and robust experimental design. To perform multiple rounds of FISH, we adapted a stripping and re-probing strategy that uses DNase I to remove probes and HCR amplification product from each previous round (Figure S1F) (Lubeck et al., 2014). For round-to-round registration and cell segmentation, we used cytosolic RNA stained with DAPI (4′,6-diamidino-2-phenylindole), which we refer to as cytoDAPI (Xu et al., 2020). Although DAPI is primarily used as a DNA stain, after DNase treatment it provides a good near-UV cytosolic stain that is abolished by RNase treatment (Figure S1G). With these improvements, EASI-FISH allows robust, high quality and multiplexed FISH imaging of thick tissue volumes (1 mm × 1 mm × 0.3 mm in pre-expansion dimensions) (Figure 1J).

### EASI-FISH data processing

Multi-round, high resolution imaging of thick tissue specimens for EASI-FISH produces multi-terabyte images that present analysis challenges. Therefore, we built computational image processing tools to handle these large datasets in a consistent and efficient manner (Figure 2A and 2B).

**Figure 2.**
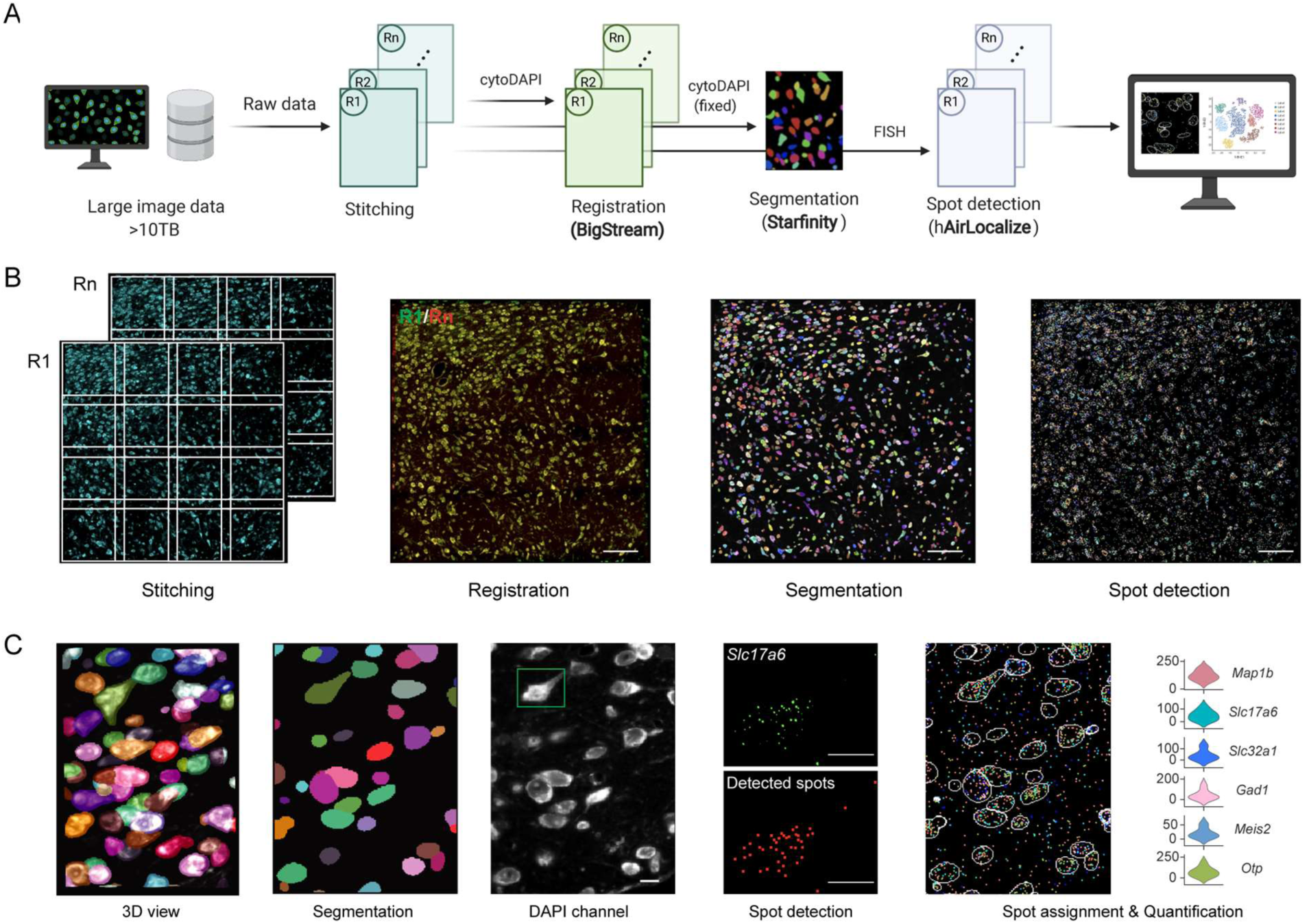
EASI-FISH analysis pipeline. **(A)** EASI-FISH data processing workflow. **(B)** Representative images showing stitching, registration, segmentation and spot detection in large image volumes. Scale bar: 100 µm. **(C)** Example of segmentation with *Starfinity* (3D view: 3D rendering of segmented cell body shape, segmentation: single optical plane) and representative Airlocalize-enabled spot detection in cell highlighted with green square. Scale bar: 10 µm.

#### Stitching

For EASI-FISH, acquiring a native sample volume of 1 mm × 1 mm × 0.3 mm requires collecting an image 8-times this volume after 2-fold expansion. For imaging these large volumes, multiple sub-volumes (tiles) were sequentially acquired, followed by computational stitching into a single large image. We used an Apache Spark-based high-performance stitching pipeline (Gao et al., 2019). The pipeline automatically performs a flat-field correction for each tile to account for intensity variations across the light sheet. It then derives the globally optimal translation for each tile that minimizes the sum of square distances to competing optimal pairwise translations estimated by phase-correlation (Preibisch et al., 2009) (Figure S2A).

#### Round-to-round registration

Next, image volumes across each round of FISH were aligned. In EASI-FISH tissue, DAPI-stained RNA provided cytosolic contours that were used for round-to-round alignment. Because sample handling could cause small deformations and 3D shifts in field-of-view (FOV) during image acquisition, we developed a robust and fully automated non-rigid registration pipeline. The analysis pipeline first performed fast global affine transformation using a feature-based random sample consensus (RANSAC) algorithm (Fischler and Bolles, 1981) (Figure S2B). The image volume was then divided into overlapping blocks and another round of feature-based affine transformation was performed before a fast 3D deformable registration (Yushkevich, 2016) was applied to each block. The pipeline is highly accurate, with 99% ± 0.8% structural similarity between fixed and nine moving image volumes (see Methods, Figure S2C), and it is more than 10-times faster compared to other deformable registration methods (e.g. ANTs) (Yushkevich, 2016).

#### Cell segmentation

DAPI-stained RNA provided a cytosolic signal that was suitable to generate cell segmentation masks, which were then used to assign transcripts to individual cells. The high accuracy of the registration pipeline allowed us to apply cell segmentation masks from a single round of imaging to all other rounds, which simplified analysis and reduced computation time.

Accurate segmentation of *in situ*-stained volumetric (3D) fluorescence image data has been a long-standing challenge that can considerably degrade the accuracy of multiplexed FISH analysis pipelines. Most approaches use thin tissue sections and an assumption of a single cell layer, which is often invalid. To overcome this challenge, we developed a deep learning-based automatic 3D segmentation pipeline, called *Starfinity* (Figure 2C). *Starfinity* is an extension of *StarDist*, an earlier cell detection approach (Schmidt et al., 2018; Weigert et al., 2020) and is based on the dense prediction of cell border distances and their subsequent aggregation into pixel affinities (see Methods). After generating appropriate training data, a *Starfinity* model was trained to predict cell body shapes from DAPI-stained RNA images, outperforming several other segmentation and machine-learning based methods (Figure S2D and **Table S3**). We manually inspected ∼5% of automatically segmented cells from 4 samples (a total of ∼4,000 out of 80,000 cells) and found that 93% of cells were properly segmented, 4% of cells over-segmented, 1% of cells under-segmented, and 2% of cells were contaminated by neighboring cells (Figure S2E). Because most over-segmented cells (62%) can be identified and rapidly corrected *post hoc* by quantitative and semi-automated criteria (see Methods), the final estimated segmentation accuracy was 95.5%.

#### Spot detection

Imaging EASI-FISH processed tissue using the Zeiss Z.1 microscope produces higher background intensity than confocal microscopy due to light-sheet excitation beam thickness. Therefore, we adapted Airlocalize (Lionnet et al., 2011) for spot detection because it implements a local background correction step that subtracts background fluorescence surrounding each spot that is associated with out-of-focus spots (Figure 2C). To rapidly process >10 terabytes of image data, we developed hAirlocalize (<u>h</u>igh throughput spot detection based on <u>Airlocalize</u>) to accelerate the spot detection process by breaking the image volume into overlapping blocks and processing each in parallel.

For cells with very high gene expression, where single spots cannot be resolved even with ∼2× linear expansion, we measured the total intensity per cell and converted the integrated intensity to spot counts based on measured well-isolated single spot intensities for these genes (see Methods). We validated this approach with low and medium expressed genes when comparing the estimated spot count with hAirlocalize measurements (Figure S2F). Based on scRNA-Seq, we found that gene expression variability between cells goes up with mean expression level (Figure S2G), such that spots for a given gene may be resolvable in some cells but not others. Therefore, we chose a spot density threshold to determine whether to use hAirlocalize spot counts or total-intensity-estimated spot counts for a given gene in each cell (see Methods).

To enable portability and reusability of the EASI-FISH analysis described above, we also built a self-contained, highly flexible, and platform agnostic computational pipeline which supports end-to-end EASI-FISH data analysis. The image analysis pipeline is freely available, open source, and modular. It can rapidly process large datasets greater than 10 TB in size with minimal manual intervention (see Methods).

### EASI-FISH is sensitive and stable

Using this analysis pipeline, we evaluated the performance of EASI-FISH. RNA detection efficiency with EASI-FISH was 81% ± 14% as measured by targeting single transcripts with two colors of interleaved 10-probe sets (Figure S1I), comparable to other methods (Chen et al., 2016; Chen et al., 2015b). To determine the sensitivity of EASI-FISH, we analyzed genes from the lateral hypothalamus (LHA) with low expression levels according to scRNA-Seq. Low-expressed genes, *Klhl13* (RNA-Seq UMI_mean_=48) and *Igf1* (UMI_mean_=15), are co-expressed in all melanin-concentrating hormone (*Pmch)*-expressing neurons in the LHA, so we used the fraction of *Pmch^+^* neurons in which we cannot detect these genes as an estimate of false negative rate at the cell level. Among the *Pmch^+^* neurons that were analyzed, 34/34 (100%) expressed *Klhl13* and 38/41 (93%) expressed *Igf1* with an average background-subtracted spot count per cell of 195 for *Klhl13* and 41 for *Igf1* (Figure S1J), indicating a low dropout (false negative) rate with EASI-FISH.

False positive spots could contribute to marker-gene spot counts by incorrect HCR amplification, spot fragmentation, or true RNA puncta detection in *en passant* neuronal processes that are adjacent to the cell body. Because we used HCR3.0, incorrect HCR amplification was low (1 per 3000 µm^3^). For analysis of false positive background spots, we selected genes that were known to be mutually exclusive, for example, in the LHA, cells expressing *Tacr3* have undetectable levels of *Pdyn* and *vice versa* based on scRNA-Seq (see Methods and below). Detection of these two genes with EASI-FISH showed complementary expression patterns in the LHA, with a very low spot count of *Pdyn* and *Tacr3* in *Tacr3^+^* and *Pdyn^+^* cells, respectively (1 per 50 µm^3^, ∼30 spots/cell) (Figure S1K). We observed a similarly low false positive background detection rate for mutually exclusive genes for *Slc17a7* (*Vglut1*) and *Gad1* in cortical neurons (Figure 1K and L).

Spot count measurements were highly reproducible across multiple rounds (Figure S1L), between replicates of EASI-FISH experiments from different animals (Figure S1M) and were also highly correlated with scRNA-Seq data (r=0.96, p=0.0081, based on measurement of *Klhl13*, *Igf1*, *Pdyn* and *Tacr3*). High RNA retention was observed when re-probing for the same targets after at least 7 rounds, even with an elapsed time of more than 40 days (93.5%, n = 4 genes averaged based on measurements from 6 animals, 2 brain regions: LHA and CEA) (Figure S1N), demonstrating excellent RNA stability. Taken together, EASI-FISH showed high reproducibility between samples, minimal RNA loss across hybridization rounds, good correlation with scRNA-Seq data, high sensitivity, and a low drop-out rate.

### Application of EASI-FISH for profiling LHA molecular markers

We applied the EASI-FISH sample processing and data analysis pipeline for in depth examination of the spatial distribution and morphological properties of molecularly defined cell types in the LHA. We performed scRNA-Seq on manually picked LHA neurons and combined this data with published LHA scRNA-Seq data collected with droplet-based methods (Mickelsen et al., 2019; Rossi et al., 2019) to determine consensus cell clusters across three datasets (Figure S3A). Clustering analysis of the combined data identified 17 glutamatergic *Slc17a6* (encoding *Vglut2*)-expressing clusters (labeled as e1-e17) and 17 GABAergic *Slc32a1* (encoding *Vgat*)-expressing clusters (labeled as i1-i17) (Figure S3B-E). Each cluster included cells from at least 2 of the 3 datasets. A combinatorial set of 24 marker-genes was selected for the subsequent EASI-FISH experiment based on specificity to map major cell types (**Table S5** and Supplemental Information fig.1).

We used EASI-FISH to profile these marker-genes in tissue volumes (1 mm × 1 mm × 0.3 mm) taken from the tuberal LHA in three 8-week-old C57BL6/J male mice (Figure 3A). The tuberal LHA is associated with eating, drinking, and arousal, and it corresponds to the area used for the scRNA-Seq tissue samples. Ten rounds of 3-plex FISH (24 total unique genes) were performed on these samples, including repeat-rounds (for validation) at the end of the data collection sequence to assess sample stability (Figure 1M). Consistent with initial EASI-FISH characterization, measurements in these samples showed excellent RNA retention (90%, n = 2 genes, 3 mice) between round 1 and round 9, even with low-expression transcripts (Figure 1N and Figure S1O). Consistent with our initial optimization analysis (above), there was high correlation with scRNA-Seq UMI measurements across 24 marker-genes (r=0.86, p=8.4 × 10^-8^), and the EASI-FISH pipeline detected an average of 13 (±1.6)-fold more molecules per cell compared to scRNA-Seq UMI counts (Figure 1O). In these large LHA samples, the false positive detection rate was low, as measured by spot counts from marker-genes that showed orthogonal expression patterns (Figure S1P).

**Figure 3.**
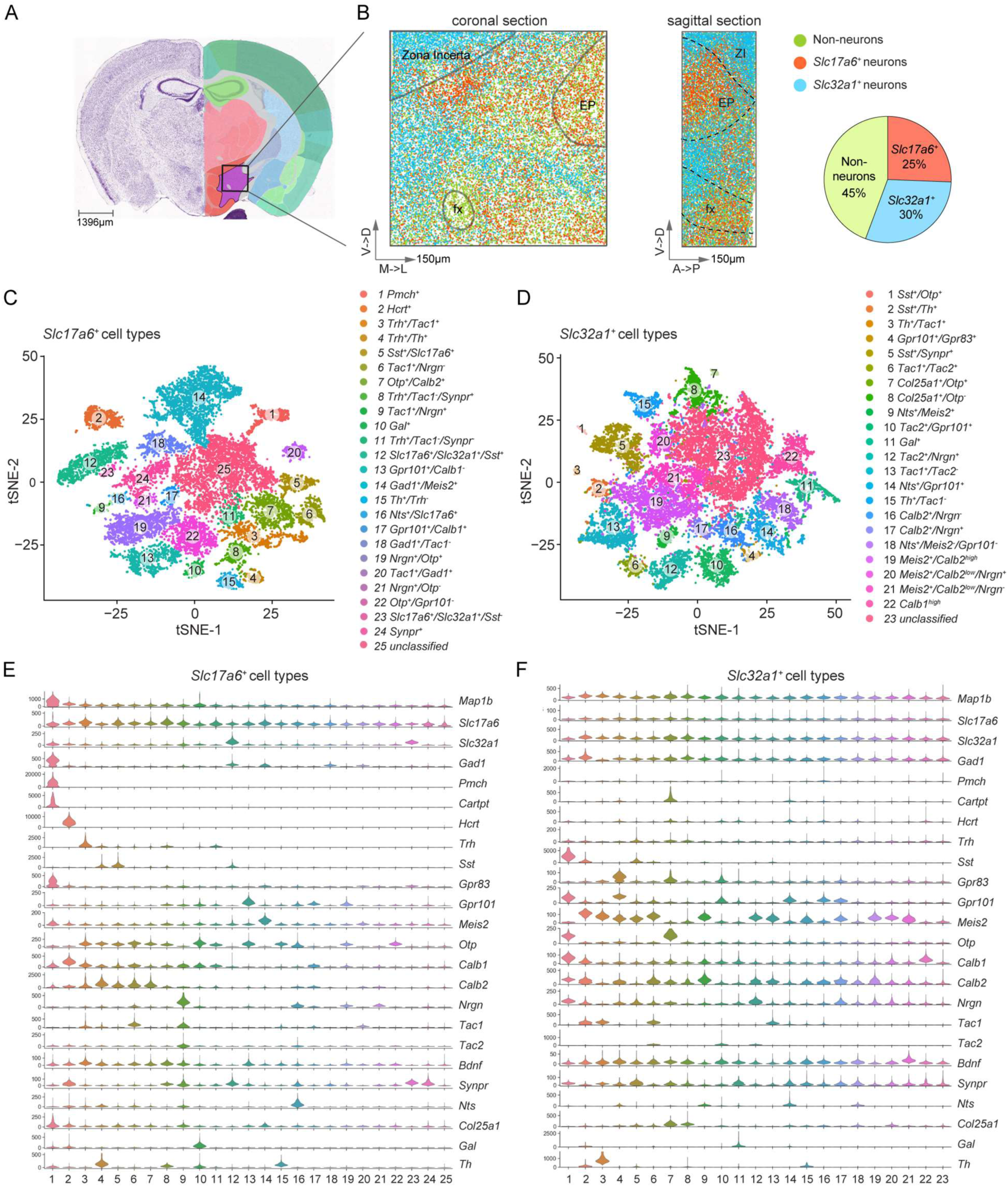
EASI-FISH for profiling LHA molecular markers. **(A)** Imaged region from LHA mapped onto the coronal mouse brain atlas. Image credit: Allen Institute. **(B)** Spatial organization and proportion of excitatory and inhibitory neurons and non-neurons in the imaged LHA region (total: 66,488 cells, 3 mice). Each dot indicates the centroid position of a cell. Image dimensions are pre-expansion. Scale arrows: 150 µm. EP: entopeduncular nucleus, fx: fornix, ZI: zona incerta **(C-D)** t-distributed stochastic neighbor embedding (tSNE) plot for **(C)** excitatory and **(D)** inhibitory neurons in the LHA, with cell types color-coded by cluster. **(E-F)** Expression (spot counts) of 24 FISH marker-genes in the **(E)** excitatory and **(F)** inhibitory clusters, shown by violin plots.

Using the EASI-FISH analysis pipeline, we identified a total of ∼86,000 cells from three specimens. Incomplete cells on the tissue surface were removed from downstream analysis, leaving 66,488 (77%) cells (details in **Table S6**). Among these cells, 55% (36,423 cells) were neurons, based on expression of the neuronal marker *Map1b*, and the remaining *Map1b^−^* cells were classified as non-neurons (Figure 3B).

We used a *de novo* approach to identify cell types in the LHA based on 24-plex marker-gene expression. Consistent with scRNA-Seq (Figure S3A), there was a dichotomy among LHA neurons based on the expression of *Slc17a6* (*Vglut2*) and *Slc32a1* (*Vgat*). Therefore, we grouped neurons into *Slc17a6*-expressing (45%, 16,394 cells) and *Slc32a1*-expressing (55%, 20,029 cells) for further analysis (Figure 3C, D). Among these neurons, 79% could be classified based on differential expression of marker-genes, while 7% of *Slc17a6*^+^ neurons (n=2787) and 14% of *Slc32a1*^+^ neurons (n=5034) showed low or no expression of marker-genes other than *Map1b*, *Slc17a6*, *Slc32a1*, *Gad1* and were grouped as unclassified clusters from each type (Ex-25 and Inh-23). Clustering *Slc17a6*^+^ neurons by marker-genes separated them into 24 molecularly defined clusters, (Figure 3E, Figure S4A, S4C and Supplemental Information fig.2) and the *Slc32a1*-expressing population was clustered into 22 molecularly defined neuronal subtypes (Figure 3F, Figure S4B, S4D and Supplemental Information fig.3) marker-gene. These molecularly defined clusters were detected in all three animals (Figure S4A, B), except Inh-1, which was located medially and was only captured in two out of the three samples because of slight medial-lateral spatial differences at the edge of the field of view. Gene expression within identified molecularly defined clusters was also highly correlated across biological replicates (Figure S4E), indicating no batch effects for EASI-FISH measurements and analysis. Among the molecularly defined clusters, Inh-3, Inh-15, and Inh-21 clusters were dominated by cells from the zona incerta (ZI), Ex-12 and Ex-23 were enriched for cells from the entopeduncular nucleus (EP) (Supplemental Information fig.4 and 5, also see Methods). FISH clusters from these large LHA datasets were correlated with scRNA-Seq data (Figure S4F, G), and all scRNA-Seq clusters were represented in the FISH dataset (Figure S4H, I, p <0.05).

**Figure 4.**
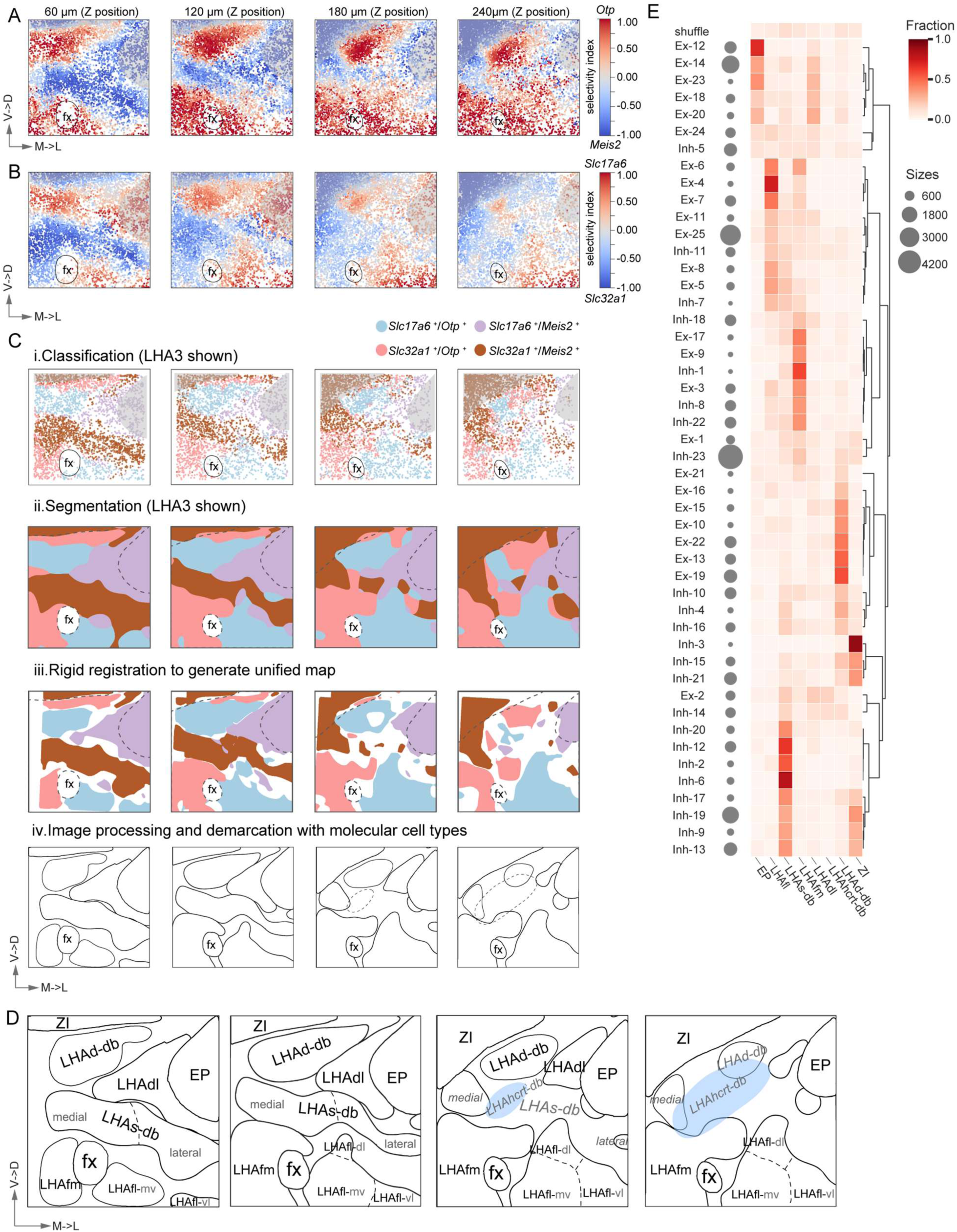
Spatial reconstruction of LHA reveals spatio-molecularly defined subregions. **(A-B)** Selective neighborhood enrichment of **(A)** *Otp*/*Meis2* and **(B)** *Slc17a6*/*Slc32a1*. Each dot indicates the centroid position of a neuron. Panels are 60 µm axial projections of sub-volumes. From left to right: anterior to posterior (rostral→caudal). **(C)** Automated workflow for 3D molecular parcellation. C-i: Classify regions based on their relative enrichment for *Otp*, *Meis2*, *Slc17a6* and *Slc32a1*. C-ii: 3D segmentation with Gaussian Mixture Models. C-iii: Consensus parcellation map after rigid registration across animals. C-iv: Boundaries from unified map are smoothed. Neighboring brain regions (ZI and EP) are shaded in gray in Figure 4A, B and C-i and highlighted with dotted line in C-ii and C-iii. Data from one animal (LHA3) was shown in Figure 4A, B, C-i and C-ii. **(D)** Spatio-molecular parcellation of the LHA, including additional subdivisions based on molecularly defined cell types. From left to right: anterior to posterior (rostral→caudal). ZI: zona incerta; EP: entopeduncular nucleus; fx: fornix; LHAd-db: LHA dorsal diagonal band; LHAdl: LHA dorsal lateral region; LHAs-db: LHA suprafornical diagonal band; LHAfm: LHA medial fornical region; LHAfl: LHA lateral fornical region. Solid lines indicate parcellations based on *Otp*, *Meis2*, *Slc17a6*, and *Slc32a1* expression. Annotations in gray and boundaries in dotted lines indicate subdomains within LHAs-db and LHAfl based on spatial segregation of molecularly defined cell types. Annotations in gray/italic indicate transition zones. *Hcrt* subdivision is shaded in light blue. Image scales are in physical units before expansion. Scale arrows in Figure 4A-D: 150 µm. **(E)** Excitatory and inhibitory cluster enrichment in the parcellated subregions, as compared to shuffling cell type identity (top, see Methods).

### Spatial reconstruction of LHA reveals molecularly defined subregions

The LHA is one of the largest and most intensively studied regions in the hypothalamus. However, previous studies do not demarcate subregions of the lateral hypothalamus in mouse (Franklin, 1997; Lein et al., 2007), and, in the rat, only limited parcellation has been proposed by combining cytoarchitectural and connectivity information (Geeraedts et al., 1990; Hahn et al., 2019; Hahn and Swanson, 2010, 2015; Veening et al., 1987).We found that many molecularly defined neuronal cell types were intermingled in the mouse tuberal LHA region. An average of 16 molecularly defined cell types were present within a neighborhood of 50 µm radius (Figure S5A), with the predominant cell type accounting for only 27% of the cells, on average (Figure S5B).

#### Computational LHA parcellation

Typically, fine neuroanatomical parcellation has been performed based on examining differences in local cell density and drawing boundaries “by eye”. Because the EASI-FISH pipeline provides detailed molecular information along with high resolution spatial information, we pursued a fully automated machine learning approach to identify LHA subregion boundaries by combining the molecular, spatial, and cell density information in these datasets. We also prioritized reproducibility of the computational parcellation by only accepting consensus regions that can be automatically aligned across samples from multiple mice.

To examine the structural organization of the LHA, we leveraged the hierarchy of the cell type gene expression profiles using the neurotransmitter transporters *Slc17a6* and *Slc32a1* as well as the transcription factors *Otp* and *Meis2* (Figure S5C and D) that have important developmental and cell specification functions in the LHA (Romanov et al., 2020). We observed a central *Meis2*-expressing wedge within the LHA that bisected *Otp*-enriched areas near the fornix and the ZI (Figure 4A), which were further subdivided by *Slc17a6* and *Slc32a1* (Figure 4B).

To execute an unbiased parcellation of the LHA, we developed a computational approach with the following steps: 1) automated volumetric segmentation based on spatial distribution of cells co-expressing combinations of *Otp*, *Meis2*, *Slc17a6* and *Slc32a1*; 2) rigid registration of the segmented volumes to align samples from biological replicates; 3) generation of a consensus parcellation across multiple samples; 4) further parcellation based on the distribution of molecularly defined cell types (Figure 4C). All steps were performed using the entire image volume, which we present for display purposes using axial projections of 4 sub-volumes (Figure 4A-C).

First, for tissue volumes from each animal, we plotted the spatial distribution of cells selectively expressing *Otp/Meis2* and *Slc17a6/Slc32a1* and applied a spatial density based smoothing (see Methods, example images shown in Figure 4A and 4B). We then classified each neuron into broad cell classes (*Otp/Slc17a6, Otp/Slc32a1, Meis2/Slc17a6, Meis2/Slc32a1*) based on the relative spatial enrichment of the two pairs of genes (example images shown in Figure 4C-i) and used Gaussian mixture models for 3D segmentation of the imaged tissue volumes (example images shown in Figure 4C-ii).

Before assessing the generalization of the LHA parcellation across animals, we performed rigid alignment on segmented tissue volumes from three animals to account for geometric differences during sample collection and imaging. The degree of alignment was validated on fiducial landmarks (ZI, EP and the fornix) (averaged intersection over union (IoU) between LHA1 and LHA3: 0.71; between LHA2 and LHA3: 0.76) as well as spatial distribution of marker-genes (Figure S5E). To estimate the consensus anatomical parcellation, we performed Simultaneous Truth and Performance Level Estimation (STAPLE) (Warfield et al., 2004), which eliminated discordance at boundaries of the segmented subregions (Figure 4C-iii).

Computational parcellation based on *Otp*, *Meis2*, *Slc17a6* and *Slc32a1*, defined 5 zones in the LHA (Figure 4C-iv and Figure 4D), most of which have not been described. Two prominent bands run diagonally at an approximately 60-degree angle from each other. The dorsal diagonal band (LHAd-db) runs directly below the ZI and is enriched for excitatory neurons (Figure 5A). A suprafornical diagonal band (LHAs-db) runs dorsal and lateral to the fornix and is enriched in inhibitory neurons (Figure 5B). The diagonal bands surround the wedge-like *Meis2*-enriched excitatory subregion in the dorsal lateral region (LHAdl), which is flanked laterally by the EP. The medial fornical area (LHAfm) is intermixed with excitatory and inhibitory neurons, with a higher fraction of inhibitory neurons (Figure 5D). The subregion lateral to the fornix (LHAfl) is enriched for excitatory neurons (Figure 5E). In the posterior portion of the tissue volume, the LHAd-db and LHAdl transition away, leaving this portion of the volume not well demarcated by the above 4 genes. Instead, this region is enriched for hypocretin (*Hcrt^+^*) neurons (Supplemental Information fig.4), an important neuropeptide secreting population, which is enriched in a diagonal band running in the same direction as the LHAd-db. We defined this sixth subregion at the caudal aspect of the LHA volume the hypocretin neuron enriched-diagonal band (LHAhcrt-db).

**Figure 5.**
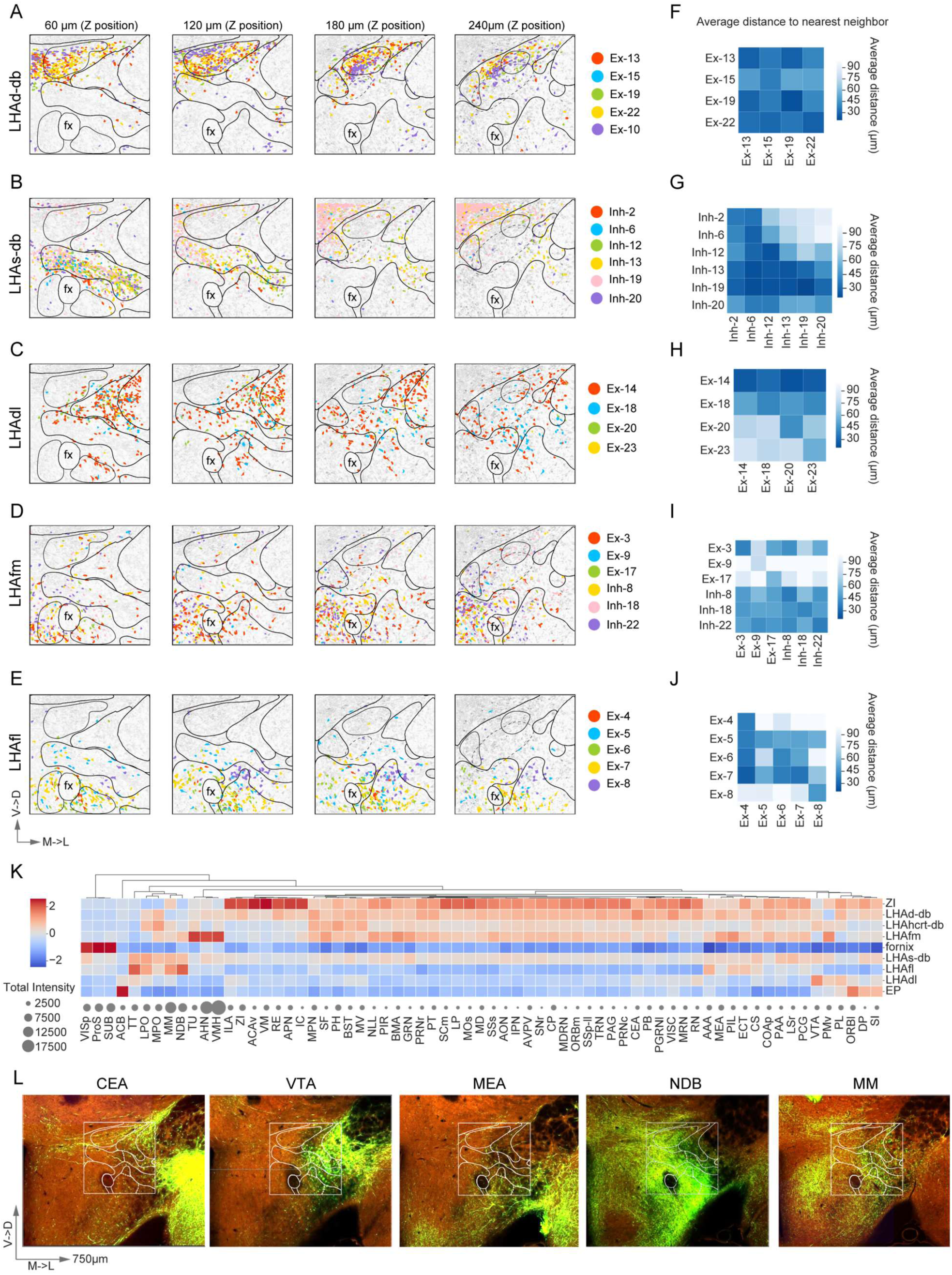
Molecularly defined cell types and axonal inputs are enriched in LHA subregions. **(A-E)** Top molecularly defined clusters enriched in **(A)** LHAd-db, **(B)** LHAs-db, **(C)** LHAdl, **(D)** LHAfm, **(E)** LHAfl subregions. As in Figure 4, panels are 60 µm axial projections of sub-volumes. From left to right: anterior to posterior (rostral→caudal). The labels indicate the middle z positions of the sub-volumes. Segmentation masks for cells belonging to the highlighted molecularly defined clusters are colored accordingly. Maximum intensity projections of cytosolic DAPI staining are shown in light gray. Image dimensions are before expansion. Scale arrows: 150 µm. **(F-J)** Average distance to nearest neighbor among molecularly defined cell types enriched in LHAd-db, LHAs-db, LHAdl, LHAfm and LHAfl, as shown in A-E. Rows represent molecularly defined cell types that were used to look for nearest neighbor and columns represents the corresponding molecularly defined cell types from which the nearest neighbor was identified (see Methods for detail). **(K)** Quantification of axonal input in LHA subregion as measured by mean fluorescence intensity (z-score normalized) based on data from the Allen Brain Atlas Connectivity database. **(L)** Representative images showing differential projections in the LHA subregions from the central amygdala (CEA), ventral tegmental area (VTA), medial amygdala (MEA), diagonal band nucleus (NDB) and medial mammillary nucleus (MM).

The subregions in the LHA that we identified by *Otp*, *Meis2*, *Slc17a6* and *Slc32a1* co-expression coarsely tracked with neuronal density in this region (Figure S5F). For example, a dense group of neurons running from the fornix to the ZI was noted previously in the rat brain and named the suprafornical LHA (Hahn et al., 2019; Hahn and Swanson, 2010, 2015). However, we discovered that consideration of molecular identity revealed a more intricate structural organization that was subdivided between the LHAfm, LHAs-db, and LHAd-db.

#### Neuronal cell types in LHA subregions

Next, we considered the relationship of molecularly defined cell types to these LHA subregions. Although molecularly defined cell types were highly intermingled, they were not randomly distributed in the LHA (Complete Spatial Randomness testing, p-value<0.05, **Table S1**). Most cell types (45/48, chi-square test, p<0.05) were spatially enriched in one or more of the LHA subregions (Figure 4E, Figure 5A-E, **Table S7** and Figure S5G), with high correlation between animals (Figure S5H). Additional differential spatial enrichment of molecularly defined cell types was observed within LHAfl and LHAs-db, allowing further division into 9 LHA subregions (dotted lines in Figure 4D and Figure S5I-J).

To make sure the parcellations proposed above were not unique to the four marker-genes used to generate them, we also looked at the molecularly defined cell type distribution in the LHA independently by measuring 1) spatial overlaps and 2) distances to averaged nearest neighbor (ANN) cell types (see Methods). The spatial overlaps between molecularly defined cell types were calculated as the number of overlapping voxels selected cell clusters occupy divided by the total number of voxels occupied by both clusters. Grouping the molecularly defined cell types based on their fractional overlap, we found that subsets of molecularly defined cell types clustered to regions that correspond to LHAs-db, LHAd-db, LHAfl and LHAfm (Figure S6A-B). Consistent with this, when we grouped cell types based on ANN distances, cell types enriched in the same subregions were clustered together (Figure S6C). Taken together, the 3D-molecular organization of LHA subregions can be determined with either a limited set of marker-genes for broad cell classes or a larger set of marker-genes for individual cell types.

#### LHAd-db

The LHAd-db contains a mixture of mainly excitatory and a few inhibitory cell types. Excitatory cell types Ex-13 (*Gpr101/Calb1^−^*), Ex-15 (*Th/Trh^−^*), Ex-19 (*Nrgn/Otp*) and Ex-22 (*Otp/Gpr101^−^*) are primarily localized to LHAd-db (Figure 5A). Ex-10 (*Gal*/*Slc17a6*) is also enriched in this subregion, but unlike the other cell types, Ex-10 forms a band around the LHAd-db (Figure 5A). In addition, a variety of inhibitory cell types are present in LHAd-db that are also distributed in the adjacent ZI and LHAs-db (Figure 4E). Some broadly distributed inhibitory cell types are also observed in LHAd-db, such as Inh-4 (*Gpr101/Gpr83*), Inh-10 (*Tac2/Gpr101*), and Inh-16 (*Calb2/Nrgn^−^*) (Figure 4E).

#### LHAs-db

Most LHAs-db cell types are inhibitory, with small population size clusters Inh-2 (*Sst/Th*), Inh-6 (*Tac2*/*Tac1*), Inh-12 (*Tac2*/*Nrgn*) and Inh-20 (*Meis2/Calb2^low^/Nrgn*) almost exclusively localize to this subregion (Figure 5B). Several LHAs-db clusters are also found in the ZI, such as Inh-9 (*Nts/Meis2*), Inh-13 (*Tac1/Tac2^−^*), Inh-17 (*Calb2/Nrgn*) and Inh-19 (*Meis2/Calb2^high^*) (Figures 4E and 5B). There is only scattered contribution from excitatory cell types, none of which were primarily located in LHAs-db. Examination of the spatial positioning of molecularly defined cell types within the LHAs-db revealed additional spatial segregation. Inh-2, 6, 9, 17 and 19 are enriched in the medial part of the LHAs-db and Inh-12 is primarily concentrated in the lateral part of the LHAs-db (Figure 4D and S5I), while Inh-13 and Inh-20 are more evenly distributed.

#### LHAdl

LHAdl is a relatively cell-sparse zone (Figure S5F), which is largely comprised of excitatory cell types similar to those in the adjacent EP: Ex-14 (*Gad1/Meis2*), Ex-18 (*Gad1/Tac1^−^*), Ex-20 (*Tac1/Gad1*), and Ex-23 (*Slc17a6/Slc32a1/Sst^−^*) (Figure 4E and 5C).

#### LHAfm

A mix of excitatory and inhibitory cell types are localized in the LHAfm, although there is a higher fraction of inhibitory neurons. Inhibitory cell types enriched in this subregion are Inh-1(*Sst/Otp*), Inh-8 (*Col25a1/Otp^−^*), Inh-18 (*Nts/Meis2^−^/Gpr101^−^*), and Inh-22 (*Calb1^high^*). Excitatory cell types are Ex-3 (*Trh/Tac1*), Ex-9 (*Tac1/Nrgn*), and Ex-17 (*Gpr101/Calb1*) (Figure 4E and 5D). Because the LHAfm is at the border of the dissection region for scRNA-Seq analyses, there appears to be a larger number of cells lacking specific marker-genes (i.e., Ex-25 and Inh-23). Some of the LHAfm cell types are also shared with the adjacent medial-ventral LHAfl, such as Ex-6 (*Tac1/Nrgn*) and 7 (*Otp/Calb2*) (Figure 5D).

#### LHAfl

LHAfl is primarily comprised of excitatory cell types with major *Trh*-expressing cells (Ex-4, Ex-8, Ex-11) and Ex-5 (*Sst*/*Slc17a6*), Ex-6 (*Tac1/Nrgn^−^*), Ex-7 (*Otp/Calb2*) (Figure 4E and 5E). Examination of the spatial positioning of molecularly defined cell types within LHAfl revealed additional segregation of this region into 3 subdomains (Figure 4D and S5J). Ex-4, Ex-6, Ex-7 are located medial-ventrally (LHAfl-mv) and Ex-11 dorsal-laterally (LHAfl-dl), with additional poorly classified excitatory neurons (Ex-25) in the ventral lateral subdomain (LHAfl-vl) (Figure S5J). In addition, there is a sparse distribution of inhibitory cell types, such as Inh-11 (*Gal*). Another two inhibitory cell types (Inh-8 and Inh-22) that were primarily localized in the LHAfm also had some representation in the LHAfl-mv.

Molecularly defined cell types enriched in the same subregions are typically spatially intermixed, based on average distance to nearest neighbor (ANN) analysis (Figure 5F-J). But segregation of some cell types within these subregions is also evident, which supports the additional subdivisions of LHAs-db and LHAfl noted above (Figures 4D and 5G, J).

#### Transcriptional relatedness of spatially clustered cell types

For molecularly defined cell types enriched in the same subregion, some corresponded to the same scRNA-Seq clusters. For example, in the LHAd-db, Ex-13 and Ex-19 both have high correlation with seq-e9 (*Otp*/*Gpr101*) from scRNA-Seq (Figure S4G), revealing additional heterogeneity within scRNA-Seq clusters. However, many molecularly defined cell types in the same subregion corresponded to transcriptionally distant scRNA-seq clusters. For instance, both Ex-4 and Ex-6 were enriched in LHAfl, but Ex-4 is a *Trh*^+^ subpopulation, whereas Ex-6 was characterized as a *Tac1*^+^ subpopulation (Figure 3E and S4G). Additionally, instances of intermingled excitatory and inhibitory cell types were common in all regions, such as Ex-3 and Inh-8 in the LHAfm and Ex-13, Ex-19 and Inh-4 in the LHAd-db. Taken together, this revealed the cell type distribution across 9 topographically organized LH subregions and showed groupings that are likely relevant to the diverse functions attributed to this complex brain region.

#### Marker-gene spatial distribution

Notably, despite the observation that some cell types showed restricted distribution in the LHA, most individual marker-genes (except *Hcrt*) were not restricted to a single subregion (Supplemental Information fig.6 and Figure S6D). Thus, co-expression relationships between multiple marker-genes are essential to reveal the underlying spatial organization of molecularly defined cell types. To determine how well combinatorial marker expression could predict spatial positions in the LHA, we trained a random-forest regression model with 24 marker-gene expression to predict the spatial positions (in x, y and z coordinate) of neurons. We found that combinatorial expression of 24 marker-genes explained 60 ± 2% of spatial variation. *Hcrt*, *Pmch* and *Trh* appeared as top features, but excluding any of these 3 genes only reduced prediction accuracy by ∼5%, showing that the model was not dominated by individual genes (Figure S6E). Exclusion of *Otp*, *Meis2*, *Slc17a6*, and *Slc32a1* did not substantially decrease the prediction accuracy either (*exclude Otp/Meis2*: 57 ± 2%, exclude S*lc17a6/Slc32a1*: 57 ± 1%, exclude *Otp/Meis2/Slc17a6/Slc32a1*: 54 ± 2%) (Figure S6F), indicating that the spatial variation we observed was not dominated by these four genes.

**Figure 6.**
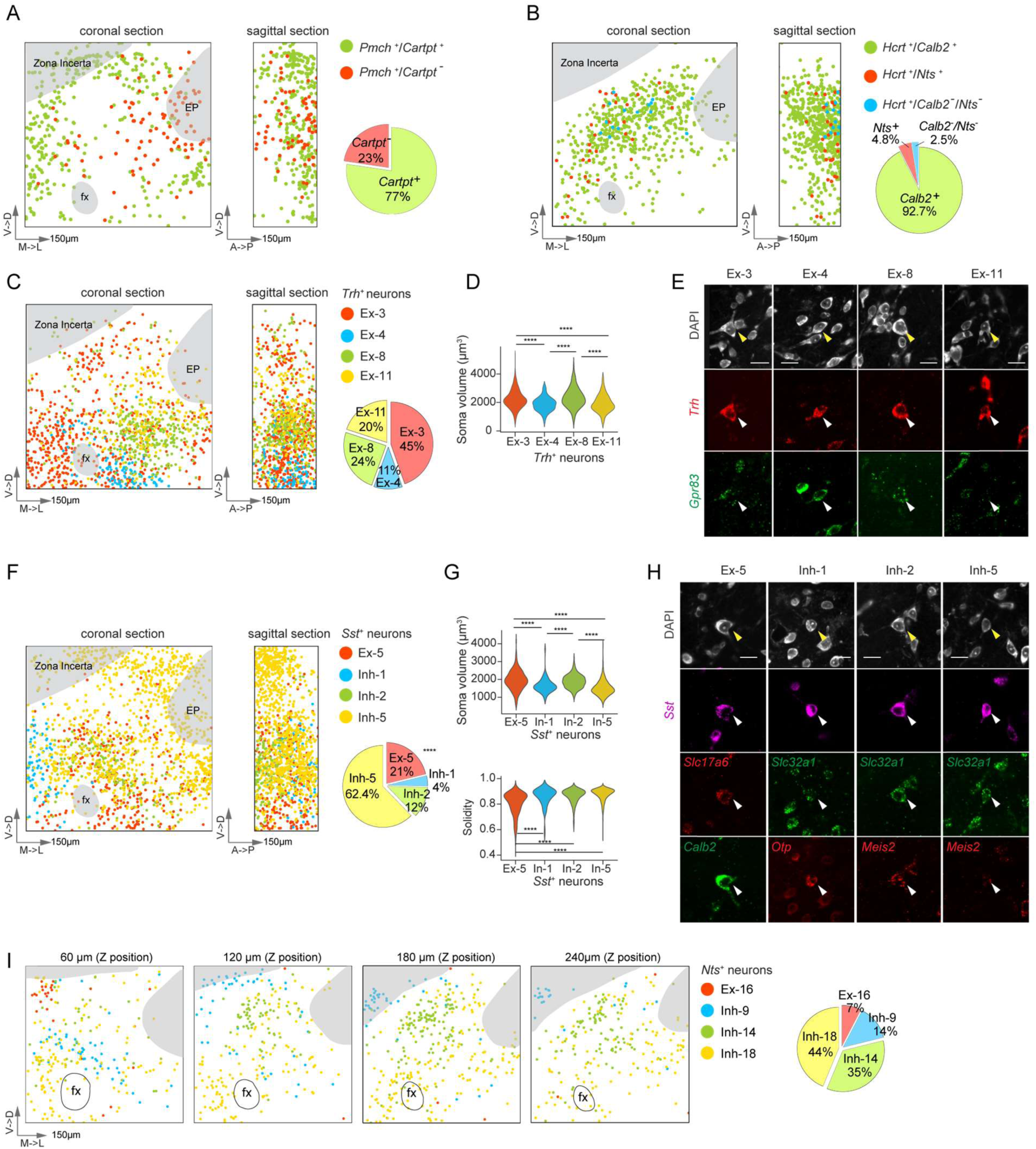
Molecular and spatial organization of major neuropeptide neurons in the LHA. **(A)** Based on *Cartpt* expression and location, *Pmch^+^* neurons can be subdivided into two populations. Max projected coronal section and sagittal section are shown with *Cartpt^+^* population in green and *Cartpt*^−^ population in red. Each dot represents the centroid position of a neuron. **(B)** *Hcrt^+^* neuron subpopulations are spatially intermixed. **(C)** *Trh*-expressing neurons can be subdivided into 4 molecularly defined clusters. Left: Spatial distribution of *Trh*-expressing neuronal clusters. Right: Percent of *Trh*-expressing neurons in each cluster. **(D)** The *Trh*^+^ clusters are also separable by soma volume. **(E)** Representative images showing the cell body morphology from *Trh*^+^ subtypes, as highlighted by arrowheads. **(F)** *Sst*-expressing neurons can be subdivided into 4 clusters. Left: Spatial distribution of *Sst*-expressing neuronal clusters. Right: Percent of *Sst*-expressing neurons belonging to each cluster. **(G)** Cell body volume (top) and shape (bottom) differences among the *Sst*-expressing clusters. **(H)** Representative images showing the soma morphology from *Sst^+^* subtypes, as highlighted by arrowheads. **(I)** *Nts-* expressing neurons can be subdivided into 4 clusters. Left: Spatial distribution of *Nts*-expressing neuronal clusters, representative coronal sections are shown to highlight the diverse anterior-posterior distribution of *Nts^+^* clusters. From left to right: anterior to posterior (rostral→caudal). Right: Percent of *Nts*-expressing neurons belonging to each cluster. Image dimension in physical units before expansion. Scale arrows: 150 µm. **** p < 0.0001. Statistics: **Table S1**.

### Axonal inputs to LHA subregions

The LHA is highly interconnected with other brain areas. We analyzed the localization of afferent axonal projections into LHA subregions by mapping this detailed LHA parcellation onto the expert-annotated Common Coordinate Framework (CCF) from the Allen Mouse Connectivity Atlas (Oh et al., 2014). We found that most axonal inputs were not broadly distributed across the LHA. The greatest selectivity was between the LHAdl and LHAfl regions, which were largely mutually exclusive in their input patterns. We found that some axonal inputs to the LHA mapped onto specific LHA subregions (Figure 5K). For example, CEA projections were enriched in the LHAd-db; VTA projections were enriched in LHAdl, MEA projections were enriched in the LHAfm, and MM, NDB projections were enriched in LHAfl (Figure 5L).

### Neuropeptide cell types in the LHA

The LHA is well known for diverse neuropeptide-expressing cell types, which display considerable heterogeneity in their somatic morphology (Shimono et al., 1985). scRNA-Seq suggested that many of the neuropeptide-expressing cell types can be further subdivided by co-expression of additional marker-genes. With EASI-FISH, we identified groupings that were also spatially distinct. The EASI-FISH analysis pipeline with *Starfinity* also recovered somatic size and shape measurements. In many cases, cell types expressing a common neuropeptide gene could be subdivided by additional marker-genes that were associated with morphological differences.

#### *Pmch^+^* neurons and *Hcrt^+^* neurons

*Pmch^+^* and *Hcrt^+^* neurons are two well-known populations in the LHA that are involved in feeding and sleep-wake behaviors. Within the tissue volume that we analyzed, 83% of *Pmch^+^* neurons were in the LHA and 17% in the ZI. *Pmch^+^* neurons could be subdivided into two populations based on *Cartpt* expression that encodes a neuropeptide that regulates energy homeostasis (*Cartpt*^+^: 77%, 388/501 neurons; *Cartpt*^−^: 22%,113/501), which was largely consistent with scRNA-Seq data (Mickelsen et al., 2019). Nearly all *Pmch^+^* neurons in the ZI were *Cartpt*^+^ (99%). Within the LHA, we observed distinct spatial distributions of the two *Pmch^+^* subpopulations, with the *Pmch/Cartpt*^−^ neurons enriched in the LHAdl, while the *Pmch/Cartpt*^+^ population segregated into a medial population and a ventral lateral population (Figure 6A). More than 92% of *Pmch^+^* neurons analyzed by EASI-FISH co-expressed *Gad1* and *Slc17a6*, consistent with scRNA-Seq data and previous report (van den Pol et al., 2004). Most PMCH neurons (77%) co-expressed the obesity-related GPCR, *Gpr83*, and there was a 2-fold greater frequency of *Gpr83* co-expressing cells in the *Cartpt^+^* population (*Pmch/Gpr83* co-expression: *Cartpt*^+^: 87%, *Cartpt*^−^: 43%).

*Hcrt^+^* neurons were spatially restricted in the LHA samples we examined and were enriched in a dorsal diagonal band that was caudal to the LHAd-db. Our manually picked scRNA-Seq dataset contained many *Hcrt^+^* neurons, revealing two main subdivisions based on the expression of *Calb2* and *Nts*. The majority of *Hcrt^+^* neurons (93%, 593/640) expressed *Calb2*, with a small population expressing *Nts* (5%, 31/640). The remainder of *Hcrt^+^* neurons were negative for both markers (2%, 16/640), which were enriched in a more caudal position. Although there were some spatial differences in the distribution of *Hcrt^+^* subtypes, they were largely intermingled (Figure 6B).

#### *Trh^+^* neurons

The thyrotropin-releasing hormone (*Trh*)-expressing neurons in the LHA have been implicated in promoting arousal behaviors (Horjales-Araujo et al., 2014). EASI-FISH identified four prominent *Trh*-expressing cell types (Ex-3, Ex-4, Ex-8 and Ex-11), which were spatially and molecularly distinct (Figure 6C). Consistent with scRNA-Seq data, 97.6% (1483/1519) of *Trh*-expressing neurons co-expressed *Otp* and were found in *Otp* zones (LHAfm and LHAfl). Compared to scRNA-Seq data, where two *Trh^+^* clusters were identified, EASI-FISH revealed additional molecular heterogeneity within *Trh*-expressing neurons that were spatially segregated. Ex-3 was spatially enriched in the LHAfm, had high expression of *Trh*, and co-expressed *Calb1* and *Tac1*. In contrast, Ex-4, Ex-8, and Ex-11 were positioned lateral to the fornix in the LHAfl. Ex-4 was enriched in the LHAfl-mv subdomain and co-expressed *Th* and *Calb2*. Ex-8 and Ex-11 were intermingled in a separate position in the LHAfl-dl subdomain and were discriminated by expression of *Gpr83* and *Synpr*, respectively (Figure 2E). In addition to spatial differences, the four *Trh* cell types also showed different cell volumes, with larger cell bodies in Ex-3 and Ex-8 relative to Ex-4 and Ex-11 (Figure 6D and E).

#### *Sst^+^* neurons

For *Sst*-expressing populations, we identified one excitatory (Ex-5) and three inhibitory (Inh-1, Inh-2 and Inh-5) cell types in the imaged LHA volume. These populations were spatially separated, with Ex-5 enriched in the LHAs-db and LHAfl, Inh-1 enriched in the LHAfm, and Inh-2 enriched in the LHAs-db. Inh-5 was diffusely dispersed across multiple LHA subregions (Figure 6F). Inh-1 had the highest *Sst* expression level (Figure 3F), and co-expressed *Gpr101* and *Nrgn*. The excitatory *Sst^+^* cluster Ex-5 and inhibitory cluster Inh-2 had larger cell bodies than Inh-1 and Inh-5 (Figure 6G and H). In addition, many neurons in the Ex-5 cluster were less convex compared to the inhibitory *Sst* neuron clusters (Figure 6G and H). We also observed co-expression of *Sst* in a subset of the *Trh*-expressing cluster Ex-4 and *Trh* in the *Sst*-expressing cluster Ex-5. Ex-4 and Ex-5 differed in their *Th* expression, with *Trh*-expressing cells in the Ex-5 negative for *Th*.

#### *Nts*^+^ neurons

*Nts^+^* neurons in the LHA have been examined repeatedly using functional perturbations during behavior, as well as with other methods (Kempadoo et al., 2013; Patterson et al., 2015). *Nts* was expressed in several transcriptionally distinct excitatory (Ex-16) and inhibitory (Inh-9, Inh-14, Inh-18) cell types in the LHA. Ex-16, the single *Slc17a6*-expressing *Nts^+^* population, was enriched in the dorsomedial region and was more anterior than the inhibitory populations (Figure 6I). Inh-9 was spatially enriched in the ZI and LHAs-db and was characterized by *Meis2* co-expression. Inh-14 was enriched in a dorsal diagonal band that spatially overlapped with *Hcrt^+^* neurons (33% overlap) and co-expressed *Gpr101* and *Galanin.* Inh-18 was more spatially dispersed in the LHA.

#### *Slc17a6*/*Slc32a1* co-expressing populations

Our image volume also included the glutamate and GABA co-releasing populations (Ex-12) in the entopeduncular nucleus (EP), which have been reported previously (Wallace et al., 2017). 62% (742/1188) of *Slc17a6*/*Slc32a1* EP neurons in our image volumes co-expressed *Sst* (Figure S7A). Interestingly, we also identified a cluster of this dual neurotransmitter cell type in the anterior part of the LHAd-db (Figure S7B and C).

### Somatic morphology in the LHA

Automatic 3D segmentation with *Starfinity* did not impose convexity constraints and thus allowed us to characterize the cell body volume and somatic morphological diversity in the LHA in an unbiased manner (Figure 7A and B). *Pmch^+^* and *Hcrt^+^* neurons were the primary large neuronal populations in this region. These neurons were ∼ 2.5-fold larger in cell body volume than the average LHA neuron volume (*Pmch^+^*: 3412±76.1 µm^3^, *Hcrt^+^*: 3690±40.6 µm^3^, average LHA excluding *Pmch^+^* and *Hcrt^+^* neurons: 1533±3.5 µm^3^) (Figure 7C). Excluding the *Pmch^+^* and *Hcrt^+^* cell types, we found that the *Slc17a6*^+^ neurons (1624±6 µm^3^) on average are close to 250 µm^3^ larger in volume than *Slc32a1*^+^ neurons (1389±3 µm^3^) (p<0.0001). Neuronal somatic volume was positively correlated with total RNA content, as indicated by cytosolic DAPI staining (r=0.93, p=0) and expression of the neuronal cytoskeleton-related gene, *Map1b* (r=0.82, p=0) (Figure S7D and E).

**Figure 7.**
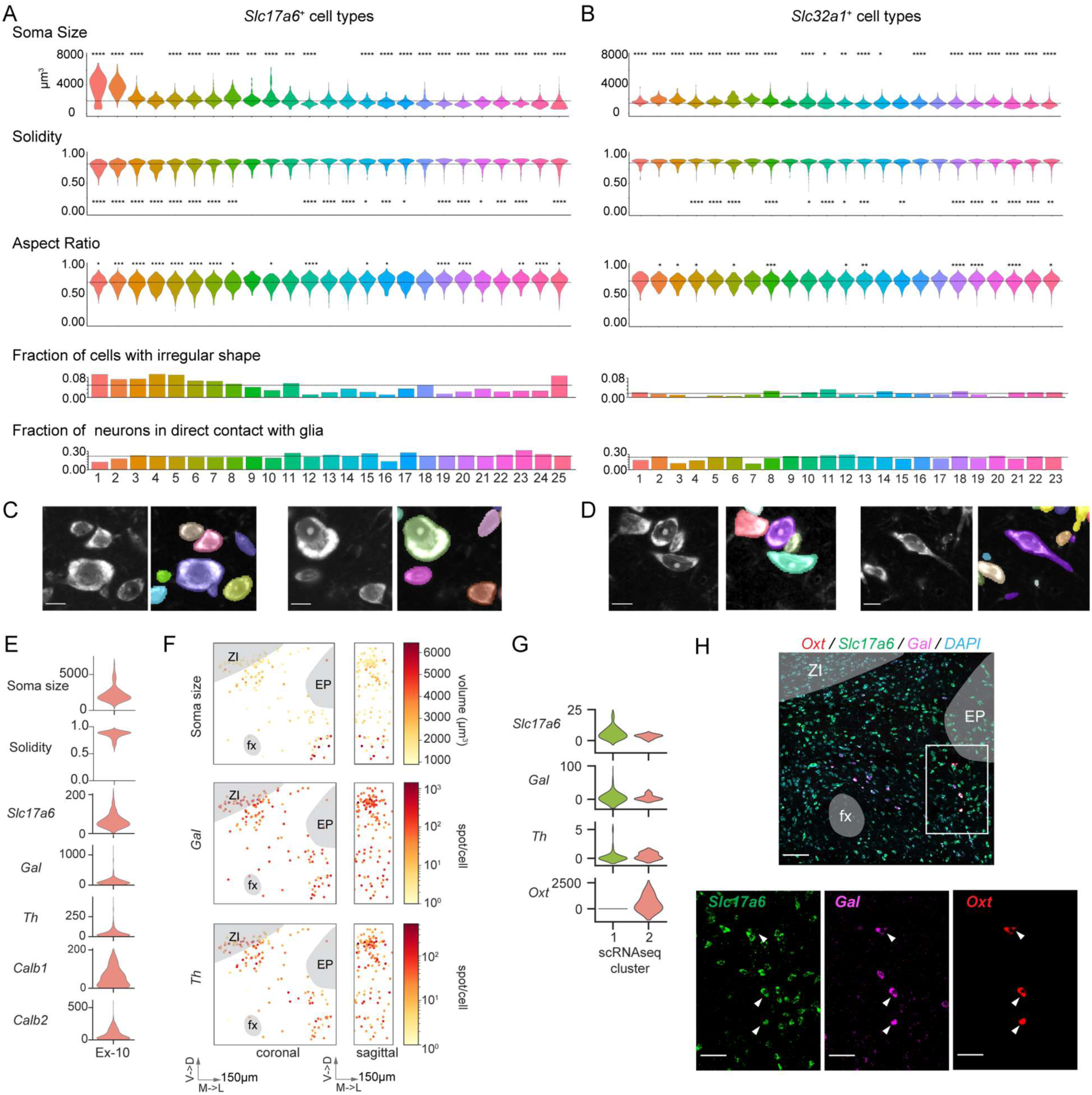
Morphological diversity in the LHA. **(A-B)** Somatic volume and morphological characteristics in **(A)** excitatory and **(B)** inhibitory clusters. Dotted lines in each plot indicate average measurements in the excitatory or inhibitory clusters. Unpaired two sample t-test (parametric) was used to compare soma size within each cluster to the population average. Unpaired two sample Wilcoxon test (non-parametric) was shown here to compare soma solidity and aspect ratio of molecularly defined clusters to the population average. * p<0.05, ** p<0.01, *** p<0.001, **** p<0.0001. Statistics: Table S1. **(C)** Representative images showing soma size diversity in the LHA. **(D)** Representative images showing soma shape diversity in the LHA. Scale bar in C and D: 10 µm. **(E)** Soma volume, shape and marker-gene expression in Ex-10 cluster. **(F)** Spatial distribution of soma size (top panel), *Galanin (Gal*, middle panel) and *Th* (bottom panel) expression in the Ex-10 cluster. A subpopulation of Ex-10 cluster is spatially enriched in the LHAfl subregion with large somata. **(G)** The scRNA-Seq cluster that corresponds to Ex-10 was further subdivided into two subclusters and differentially expressed gene, *Oxt* was identified. Gene expression (UMIs) in corresponding scRNA-Seq subclusters are shown here. **(H)** *Oxt* marks the Ex-10 subpopulation. Top: Representative single plane image showing the spatial distribution of Ex-10 subpopulation marked by *Oxt* expression. Scale bars: 100µm. Bottom: zoom-in of the white box in the top image showing co-expression of *Slc17a6* and *Galanin (Gal)* in the *Oxt* subpopulation. Scale bar: 25 µm.

We also evaluated 3D somatic shape based on cellular aspect ratio and solidity measurements (Figure S7F). To identify cells with the most extreme shapes (Figure 7D), we selected for cells with low aspect ratio (elongated, <0.7) and solidity (less convex, <0.7). This revealed a higher fraction of irregular somatic shapes in *Slc17a6*^+^ neurons (6%) compared to the *Slc32a1*^+^ neurons (2%). Neurons with most irregular shapes were enriched in the LHAfl region (Figure S7G).

### Iterative refinement of cell type marker-genes

Further spatial and morphological variations were observed in these identified molecularly defined cell types, which raised the question of whether they can be subdivided. For example, we found that the distribution of cell sizes in cell type Ex-10 (*Slc17a6*/*Gal*) had a long tail of large neurons (Figure 7E). In addition, this cell type was spatially enriched in two subregions (LHAd-db and LHAfl-vl) (Figure 7F), with the large neurons primarily in the LHAfl-vl subregion. This raised the possibility that this is a distinct but rare cell type.

To test this hypothesis, we subdivided the most similar scRNA-Seq cluster and identified Oxytocin (*Oxt*) as the top differentially expressed gene between the subdivided scRNA-Seq clusters (Figure 7G). To examine whether *Oxt* can be used to separate this population from the mediodorsal *Slc17a6*^+^/*Gal*^+^ subpopulation, we probed for *Slc17a6*, *Gal* and *Oxt* with EASI-FISH in a new sample to independently validate the existence of this population. Indeed, *Oxt* expression was detected in a small group of neurons in the ventral lateral part of the LHA. Consistent with our predictions, these neurons had large cell bodies (3089±140 µm^3^) and co-expressed *Slc17a6* and *Gal* (100%, 13/13) (Figure 7H). Taken together, this showed how spatial and morphological measurements can facilitate the discovery of rare cell types with unusually large cell bodies, using a limited number of marker-genes.

## Discussion

We report a resource that includes new methods and a turnkey computational analysis pipeline for multi-round FISH datasets in thick sections of brain tissue. EASI-FISH enables quantitative *in situ* measurements of gene expression with cellular resolution using commercial laboratory equipment in a format that includes single RNA puncta counts, detailed spatial information, and morphological characteristics of the underlying cells. We demonstrated EASI-FISH in the portion of the LHA that is associated with eating, drinking, arousal, and sleep. We characterized the molecular, spatial, and morphological diversity of LHA neurons and discovering an unanticipated degree of anatomical organization.

EASI-FISH bridges a gap between commonplace single-round 3-plex FISH experiments and highly specialized methods for obtaining spatial gene expression information from 100s-1000s of genes. To prioritize ease-of-use, we chose to use non-barcoded sequential probing, where transcripts are amplified and distinguished using different fluorophores. Each round of FISH can be analyzed independently, which allows flexibility and tolerance in experimental design. Although the number of marker-genes scales linearly with the number of rounds, non-barcoded sequential probing removes constraints on gene selection based on expression levels and requirements for voxel-precision alignment from round-to-round (Moffitt et al., 2018; Shah et al., 2017). Spot-to-spot alignment is likely possible using this method, which would facilitate barcoding methods and greatly increase throughput for the number of genes that could be probed, but this would also increase imaging complexity and analysis requirements.

The *Starfinity* cell segmentation method improves 3D morphological reconstruction of cells, which is a key aspect of multiplexed FISH analysis. The EASI-FISH analysis pipeline is highly distributed and flexible. Individual computational components can be exchanged for others, such as alternative 3D segmentation methods (e.g. Cellpose (Stringer et al., 2021)). Although the analysis reported here was initially performed on an LSF compute cluster, the pipeline is containerized so that it can also be executed on a standard high-performance workstation or public cloud platforms. The convenient tissue processing, image registration, and image analysis tools offered by this resource should enable widespread laboratory adoption of high-multiplex FISH methods.

EASI-FISH was developed for 300-µm-thick tissue sections. This thickness is suitable for aligning samples from different subjects. We found that using thick tissue sections was critical for mapping the novel structural subdivisions that we discovered in the LHA across multiple animals. In addition, it allowed identification of LHA parcellation that showed the greatest inter-subject consensus. This is important because we found structural variability in the LHA spatio-molecular domains, so it was necessary to establish common features. Tissue thickness around 300 µm is also consistent with typical dimensions for other neuroscience data collection modalities that could be potentially combined with EASI-FISH, for example brain slice recordings or *in vivo* two-photon calcium imaging (Xu et al., 2020). Application to thicker tissue sections is possible but data sizes become considerably larger, especially in light of tissue-expansion, creating challenges for data storage and manipulation.

We chose to demonstrate EASI-FISH in the LHA because it has been intensively studied for decades but there has been limited progress towards neural circuit principles in this brain region (Burdakov and Karnani, 2020; Carus-Cadavieco et al., 2017; Kosse and Burdakov, 2018; O’Connor et al., 2015). This is, in part, related to the rudimentary understanding of LHA structural organization as well as the fact that single marker-genes are insufficient to represent discrete cell types. Applying EASI-FISH to the LHA, we found a remarkable level of subregional parcellation into which distinct cell types were selectively localized. The EASI-FISH pipeline generated high-quality quantitative spatial gene expression data, which enabled a machine learning approach to identify LHA subregions based on combinations of four marker-genes. We examined the high-level organization of the LHA using *Otp* and *Meis2* expression, because there was an established role for these genes in specifying large areas of the hypothalamus during development (Ferran et al., 2015; Romanov et al., 2020). Within these domains, we found distinct patterns of *Vgat* and *Vglut2* expression. Consistent with the parcellation based on combinations of these four genes, we observed that many individual molecularly defined cell types were highly enriched in just one of these specific spatial subregions. Past efforts at parcellation of the rat LHA have not elucidated most of the regions that we examined (Geeraedts et al., 1990; Hahn, 2010; Hahn and Swanson, 2010). For example, as with prior work in the rat brain, we found evidence for a suprafornical cell dense zone (Hahn and Swanson, 2010) in the mouse. However, consideration of molecularly defined cell type information subdivided this suprafornical cell density into distinct, molecularly defined laminar structures that extended laterally, distorted by fiber bundles going through this region. More generally, molecularly defined cell types are increasingly appreciated as a fundamental unit of brain organization. Thus, brain parcellation should incorporate organizational principles that are reliant on cell type distributions, which requires detailed spatial analysis of cellular multi-gene co-expression relationships.

The hypothalamus is well known for magnocellular neurons, including *Pmch^+^* and *Hcrt^+^* neurons in the LHA (Croizier et al., 2013; de Lecea et al., 1998; Elias et al., 1998; Li and de Lecea, 2020; Qu et al., 1996; Sakurai et al., 1998; Skofitsch et al., 1985). The high-quality segmentation of the EASI-FISH pipeline allowed us to examine the relationship of marker-gene expression to cell size and somatic morphology in this region. With morphological analysis of LHA cell types, we demonstrated that Ex-10 (*Th/Gal*) could be subdivided to reveal an *Oxt*-expressing subpopulation that defined a spatially segregated set of large magnocellular neurons. This is another advantage of the simple multi-round EASI-FISH method where a few dozen marker-genes can be mapped and efficiently analyzed, leading to generation of new hypotheses about transcriptionally defined cell types. These hypotheses can be examined by re-analysis of the scRNA-Seq dataset, and because of high RNA stability after gel-anchoring, additional marker-genes can be subsequently probed on the brain region under investigation. More generally, the output of the EASI-FISH analysis pipeline provides a comprehensive summary of molecular and morphological cellular heterogeneity in the lateral hypothalamus. This approach will be similarly useful in other brain regions, especially poorly subdivided subcortical areas in hypothalamus and hindbrain.

A clear challenge in neuroscience is to establish the functional importance of the different cell types in complex brain areas, for example by immediate-early-gene analysis (Kim et al., 2019; Moffitt et al., 2018) or by genetically targeting reporters and perturbation tools (Fenno et al., 2020). Our analysis of the LHA highlights the extent of spatial diversity in cell types and axonal inputs that should be taken into account for investigating LHA function. Thus, detailed examination of LHA function will be dependent on convenient and robust multi-gene co-expression analysis for all aspects of the LHA. In addition, the boundaries of these structures change along the anterior-posterior axis, which could be mapped in the future with assistance from a more anatomically distributed scRNA-Seq LHA dataset and by extending the EASI-FISH method throughout the LHA. Based on this, EASI-FISH fills an important gap in neural circuit research by facilitating a seamless extension of cell type analysis generated in dissociated neurons to any experimental modality that requires information about *in situ* anatomical localization. These attributes of EASI-FISH make it well-suited to transform the currently complex spatial analysis of ≥30-plex gene expression into a routine procedure that can be readily integrated with other high information data streams. This will lower the barrier for the field to make rapid progress towards extending the molecular revolution in neuroscience to functional and systems analysis, which is essential for understanding the interplay between neural coding and molecular properties in behavior and disease.

## Data and Code Availability

We are committed to open science with the scientific community. Details on software (github.com/multiFISH/EASI-FISH) and pipeline (https://github.com/JaneliaSciComp/multifish) generated for EASI-FISH data processing can be found on Github. Example dataset and Starfinity training data and model used in this manuscript can be found on figshare (https://doi.org/10.25378/janelia.c.5276708). All data generated with EASI-FISH can be accessed from figshare at https://figshare.com/s/72cec70e8a057bc749f0. Custom codes used in the LHA study are provided at github.com/multiFISH/LHA_analysis.

## Acknowledgments

This work was conducted as part of the multiFISH Project Team at Janelia Research Campus and supported by the Howard Hughes Medical Institute. D. Alcor and M. DeSantis provided microscopy support. A. Hu, M. Copeland, S. C. Michael, and K. M. McGowan performed mouse perfusion and brain sectioning. The Janelia Vivarium staff provided animal care. L. Wang performed cell sorting and sequencing data collection. S. Preibisch, H. Otsuna, and D. Ackerman consulted on software. T. Lionnet at NYU School of Medicine provided Airlocalize. P. Hanslovsky, I. Pisarev, and J. Bogovic helped with image processing and visualization. L. Lavis and S. Banala suggested fluorescent dyes. S. Long consulted on smFISH methods. We consulted J. Hahn at USC about LHA anatomy. A. Osowski and M. Jefferies provided administrative support and G. Ihrke staff support.

## Author contribution

S.M.S., P.W.T., and W.K. conceptualized and supervised the project; P.W.T., Y.W., and M.E. developed the method with input from S.M.S. and T.W.; G.F. developed the registration software with input from T.W., Y.W. and S.X.; M.W., U.S., S.S., and E.W.M. developed the segmentation software; Y.W. annotated training data and evaluated the segmentation prediction; Y.W. and G.F. wrote scripts for distributed spot detection; K.R. and C.G. integrated the analysis into Nextflow pipeline; F.E.H. and A.L. performed the LHA scRNA-Seq experiment and identified marker-genes; M.E. collected the LHA EASI-FISH datasets; Y.W. performed all data analysis with input from S.M.S. and P.W.T.; Y.W. and S.M.S. wrote the manuscript with input from all co-authors.

---

**Figure S1.**
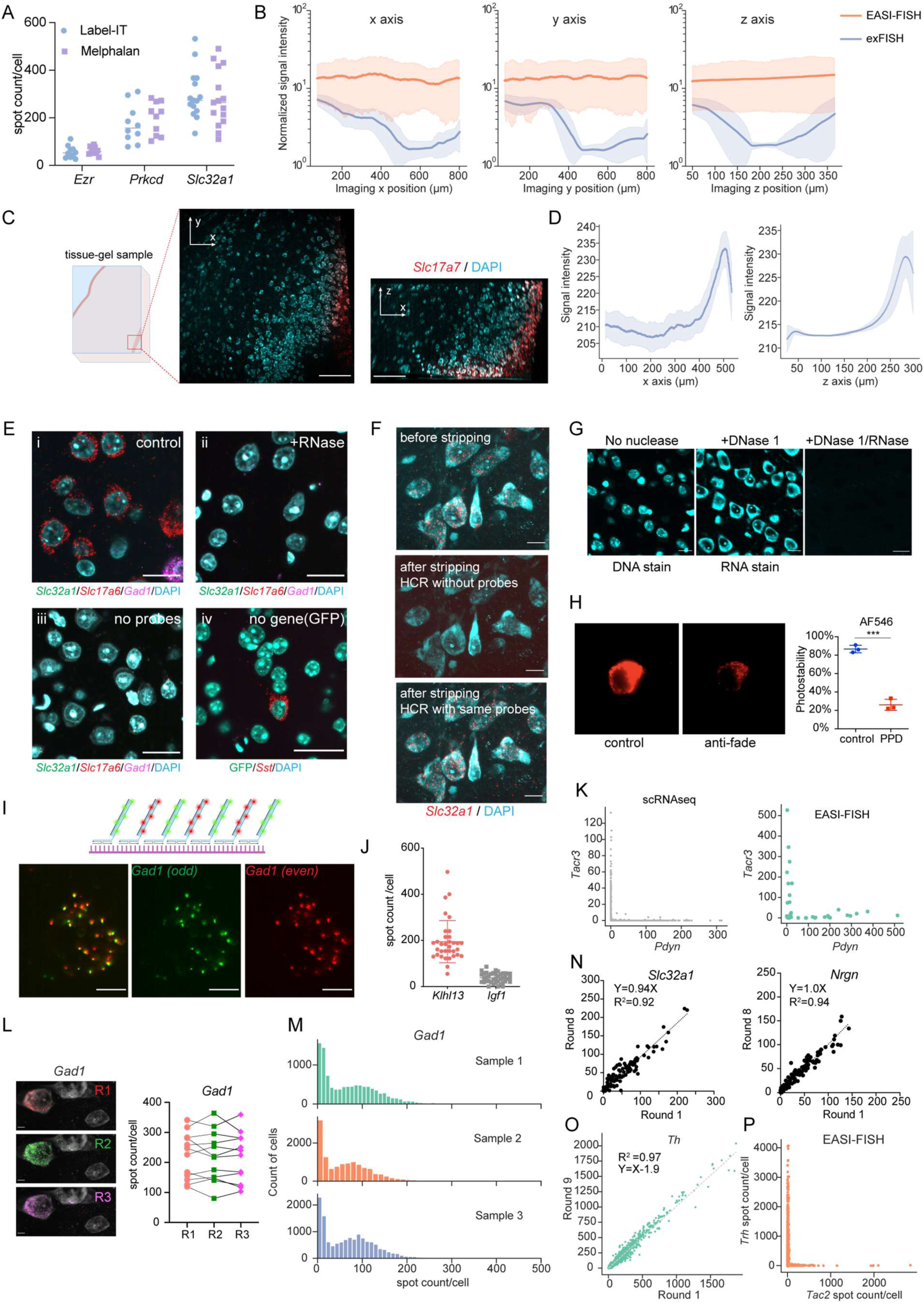
optimization of EASI-FISH protocol, related to Figure 1. **(A)** *Ezr*, *Prkcd* and *Slc32a1* spot counts detected with different RNA anchoring methods (Label-IT or Melphalan). **(B)** EASI-FISH was optimized to eliminate light scattering in large tissue specimens (0.8 mm × 0.8 mm × 0.3 mm), as demonstrated by a continuous distribution of signal intensity across the x, y and z axes compared to original ex-FISH protocol (Chen et al., 2016). For exFISH, signal intensities were still compromised at the 0.8 mm position in x and y because that is not the edge of the tissue (tissue volume used here was bigger than 0.8mm × 0.8mm in x and y dimension). Shading denotes standard deviation (SD) around mean. **(C)** Representative images showing RNAscope detection of *Slc17a7* in 300µm EASI-FISH cortical sample. A small field of view on the right edge of the tissue-gel sample is shown here. Left: schematic showing where the representative image volume (middle and right) was taken, middle: single optical plane from the image volume, right: side-view of the image volume. Scale bar: 100µm. **(D)** Signal intensity quantification of the image volume shown in C. Shading denotes standard deviation (SD) around mean. **(E)** EASI-FISH is optimized for high specificity, as indicated by minimal spot detection in the absence of RNA (ii), low non-specific binding/HCR initialization of hairpins in the absence of probes (iii), and low spot detection in the absence of the target gene, GFP (iv, *Sst* detection as positive control) as compared to control (i). Scale bar: 25 µm. **(F)** HCR products are completely eliminated with DNase 1 (middle panel) and can be re-probed (bottom panel) without signal loss, shown as z-projection. Scale bar: 10 µm. **(G)** DAPI staining of DNA (left) and RNA (middle) in EASI-FISH samples. Note that cells with low RNA content can have very weak DAPI RNA staining. Right panel: no DAPI staining in tissue in the absence of oligonucleotides (DNase and RNase treatment). Scale bar: 25 µm. **(H)** Representative image (left) and quantification (right) showing rapid photobleaching of Alexa-fluor 546 (AF-546) in the presence of anti-fade. **(I)** EASI-FISH detection efficiency assessed by co-localization of interleaved HCR probes targeting the same gene, *Gad1*. **(J)** EASI-FISH false negative detection as indicated by co-detection of low expressors *Klhl13* and *Igf1* in PMCH neurons from the LHA. **(K)** EASI-FISH false positive detection in mutually exclusive genes. Left: UMIs of mutually exclusive genes based on scRNA-Seq, Right: Spot counts in these genes with EASI-FISH. **(L)** Reliable detection of the same gene (*Gad1*) across 3 rounds of FISH in cortex. Representative images on the left and quantifications on right. Scale bar: 5 µm. **(M)** Reliable spot detection of the same gene (*Gad1*) in biological replicates. **(N)** Spot count of *Slc32a1* and *Nrgn* in round 1 and round 8. **(O)** RNA stability and **(P)** false positive detection in the LHA samples used for EASI-FISH demonstration. *** p<0.001. Statistics: **Table S1**.

**Figure S2.**
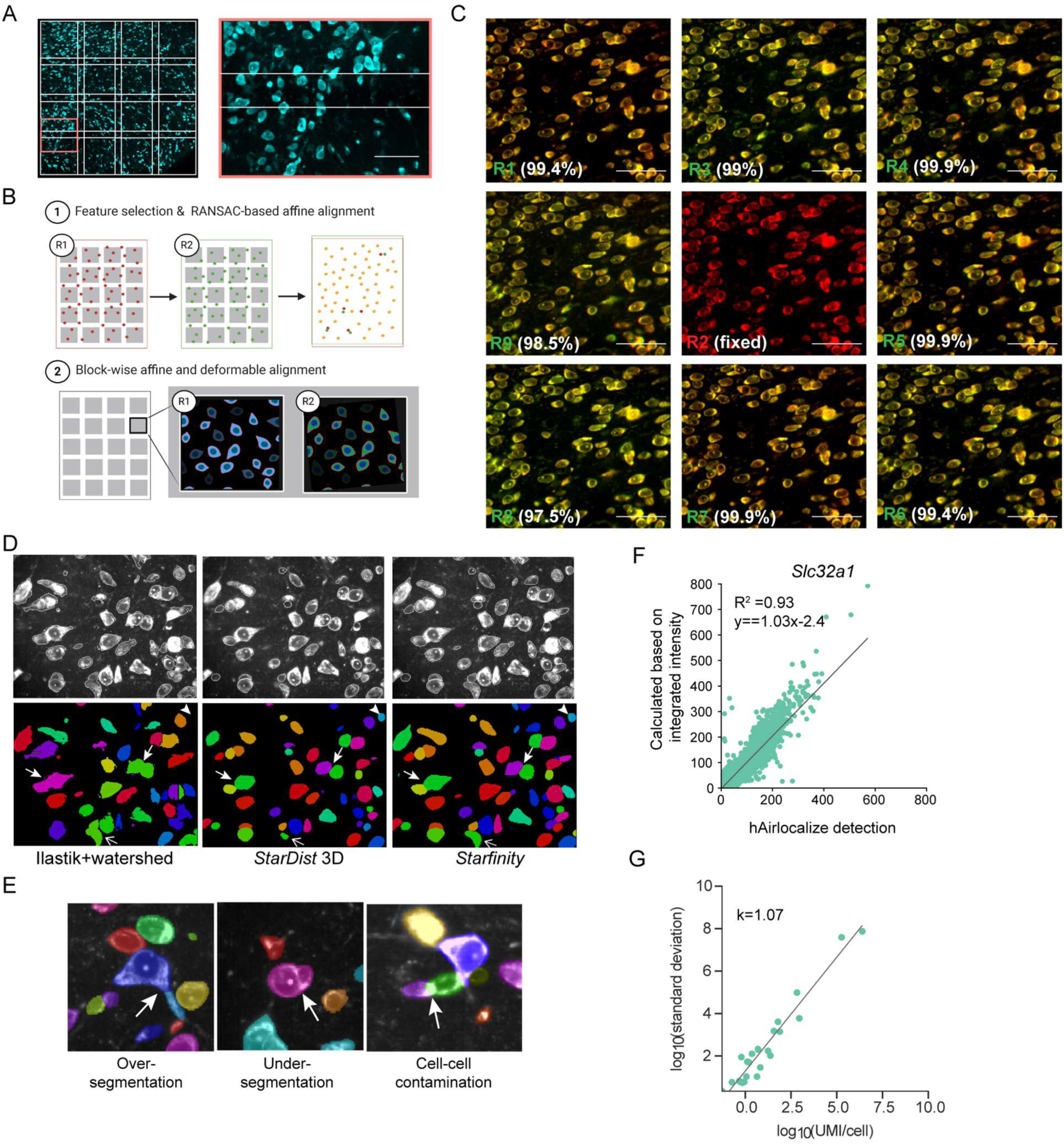
EASI-FISH data processing pipeline, related to Figure 2. **(A)** Representative multi-tile images after stitching. Left: Representative single plane image after tile-tile stitching. White grids indicate the dimensions of each tile with overlaps. Right: zoom-in of region highlighted in red box (left) to demonstrate overlapping regions after stitching. Scale bar: 50 µm. **(B)** Schematic illustration (cartoon images) of the round-to-round registration pipeline. **(C)** Representative images showing registration of 9 rounds of EASI-FISH images based on RNA-staining (DAPI). Round 2 was used as the fixed round (shown in red). Structural similarity to fixed round is shown. Scale bar: 50 µm. **(D)** Segmentation comparisons among Ilastik in combination with watershed (left), *StarDist* (middle) and *Starfinity* (right). Representative segmentation errors are highlighted by arrows. Thick arrows indicate under-segmentation errors (cell-cell merges); Arrowheads indicate under-detection errors; Thin arrows indicate *StarDist-*specific errors due to star-convexity constraints. **(E)** Representative segmentation errors in *Starfinity*, over-segmentation (one cell was split into multiple labels), under-segmentation (multiple cells assigned the same label), contamination (segmentation boundaries not properly drawn), highlighted with white arrows. **(F)** Spot count comparison between Airlocalize and integrated intensity-based estimation in low-expressor-cells (*Slc32a1*). **(G)** Gene expression level variation, shown as log_10_ (standard deviation), as a function of average gene expression log_10_(average UMI/cell) in scRNA-Seq population. Statistics: **Table S1**.

**Figure S3.**
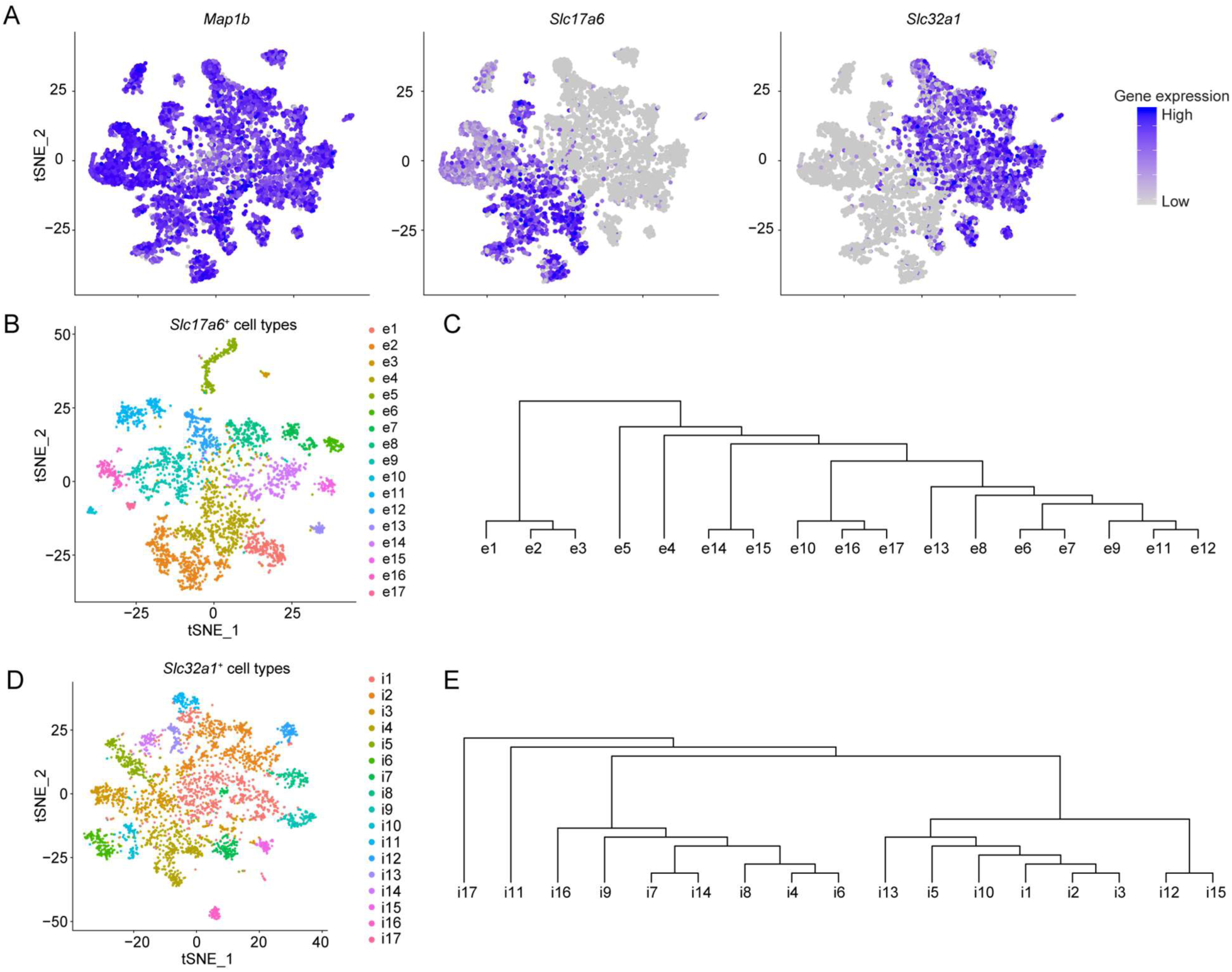
scRNA-Seq analysis of LHA molecularly defined cell types. **(A)** tSNE plot of scRNA-Seq data showing *Map1b, Slc17a6*, and *Slc32a1* expression in the LHA neurons. **(B-C)** 17 molecularly defined cell types identified from the *Slc17a6^+^* population. **(B)** tSNE plot for *Slc17a6^+^* clusters in the LHA, with cells color-coded by cluster. **(C)** Hierarchical analysis of the *Slc17a6^+^* clusters. **(D-E)** 17 molecularly defined cell types are identified from the *Slc32a1^+^* population. **(D)** tSNE plot for *Slc32a1^+^* clusters in the LHA, with cells color-coded by cluster. **(E)** Hierarchical analysis of the *Slc32a1^+^* clusters.

**Figure S4.**
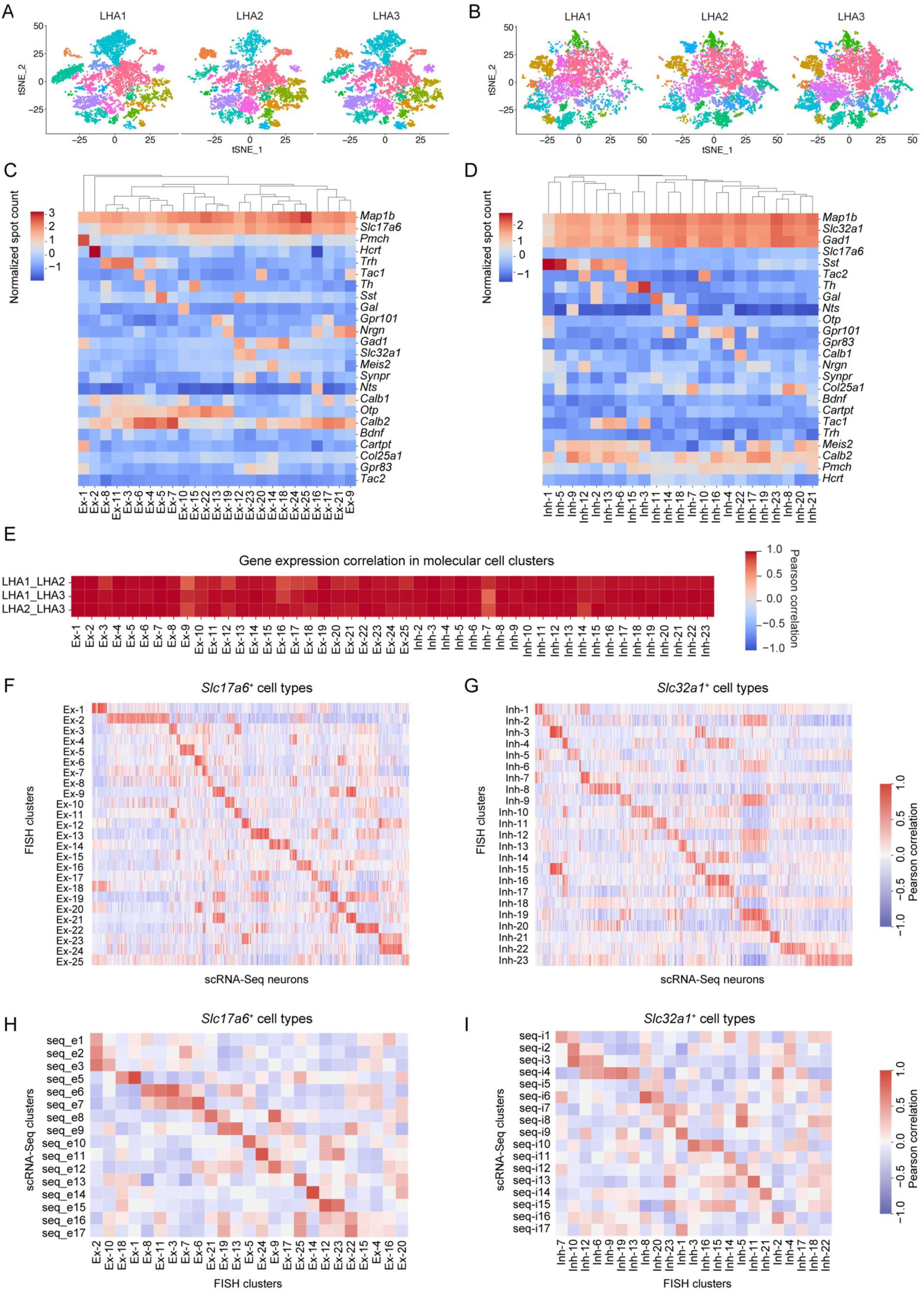
EASI-FISH for profiling LHA molecular markers, related to Figure 3. **(A-B)** tSNE plot of showing sample-to-sample variations in *Slc17a6^+^* and *Slc32a1^+^* clusters (from left to right: LHA1, LHA2 and LHA3), see Figure 3C-D for cell type correspondence. **(C-D)** Heatmap and hierarchical analysis of marker-gene expression for **(C)** *Slc17a6^+^* and **(D)** *Slc32a1^+^* molecularly defined cell types. **(E)** Correlation analysis of marker-gene expression between samples for the molecularly defined cell types. **(F-G)** Correlation analysis between EASI-FISH clusters and scRNA-Seq neurons. **(H-I)** Correlation analysis between EASI-FISH clusters and scRNA-Seq clusters based on average marker-gene expressions. The pairwise Pearson correlation coefficients were computed using the z-score normalized spot count of EASI-FISH and the normalized UMIs from scRNA-Seq data.

**Figure S5.**
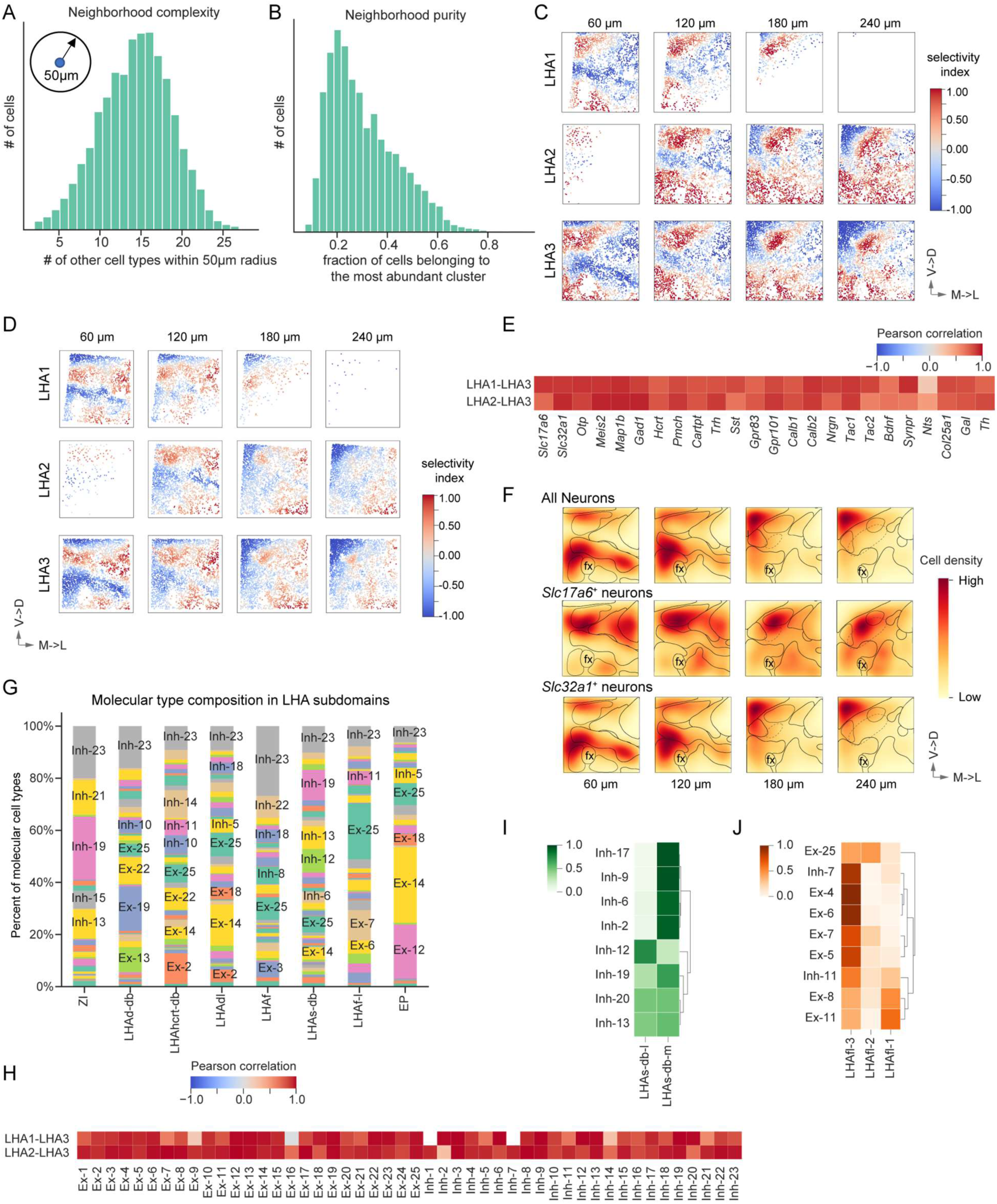
Spatio-molecular parcellation of the LHA, related to Figure 4 and Figure 5. **(A)** Neighborhood complexity analysis of the number of neuronal types within a 50 µm radius of each neuron. **(B)** Neighborhood purity analysis for the fraction of neurons that are part of the most abundant cluster within the neighborhood (50 µm radius). **(C)** Relative *Otp*/*Meis2* neighborhood enrichment analysis across three samples. LHA1 and LHA2 were aligned to LHA3 with rigid registration. **(D)** Relative *Slc17a6*/*Slc32a1* neighborhood enrichment analysis across three samples. **(E)** Correlation analysis of marker-gene spatial distributions across samples. Image volumes were first aligned and then binned to 10 × 10 × 4 bins. The number of marker-gene positive cells in each bin was calculated and used for the correlation analysis. **(F)** Spatial neuronal density (top), *Slc17a6^+^* neuronal density (middle) and *Slc32a1^+^* neuronal density (bottom) in the LHA, computed using kernel density estimation (kde). **(G)** Molecularly defined cell type compositions in LHA subregions. **(H)** Correlation analysis of molecularly defined cluster spatial distribution across samples. Fraction of molecularly defined neuron clusters in each subregion was used for the correlation analysis. **(I)** Relative molecularly defined cluster enrichment in the medial and lateral part of the LHAs-db, as indicated in Fig 4D. **(J)** Selective molecularly defined cluster enrichment in the LHAfl subdomains, as indicated in Fig 4D.

**Figure S6.**
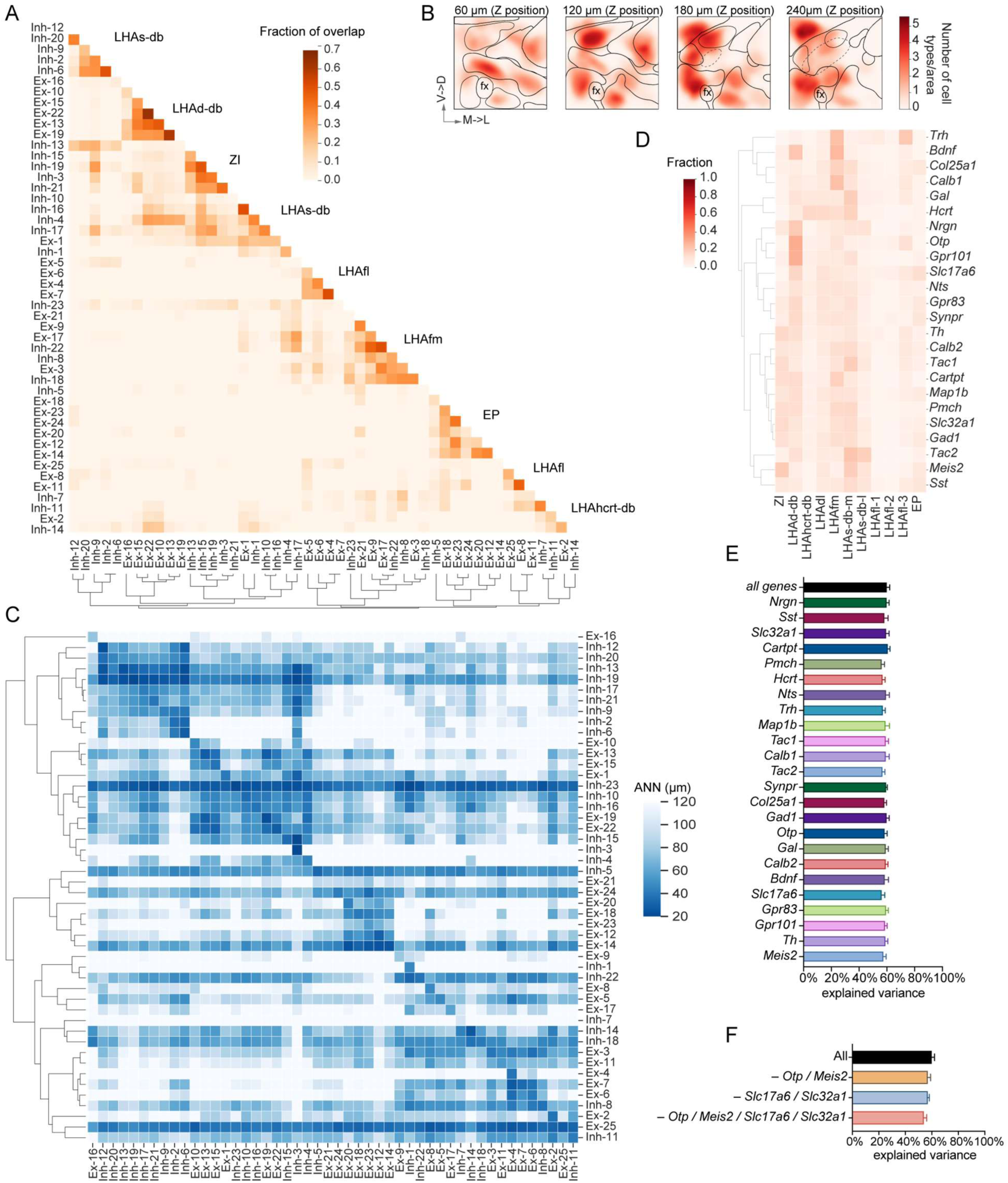
Molecularly defined cell types are enriched in LHA subregions, related to Figure 4 and Figure 5. **(A)** Molecularly defined cell types grouped by their fractional overlap. **(B)** Molecularly defined-cluster-occupancies in the LHA. Colormap indicates the number of clusters occupying each location. **(C)** Average nearest neighbor analysis between molecularly defined cell types. **(D)** Spatial distribution of marker-gene expression in parcellated LHA subregions. **(E)** Random forest regression models were used to predict the spatial position of neurons based on all 24 marker-genes. The prediction accuracy with all 24 marker-genes were calculated and compared to that with 23 marker-genes, with the selected marker-gene removed. **(F)** Prediction accuracy of neuronal spatial position in the presence or absence of *Otp*, *Meis2*, *Slc17a6*, *Slc32a1*.

**Figure S7.**
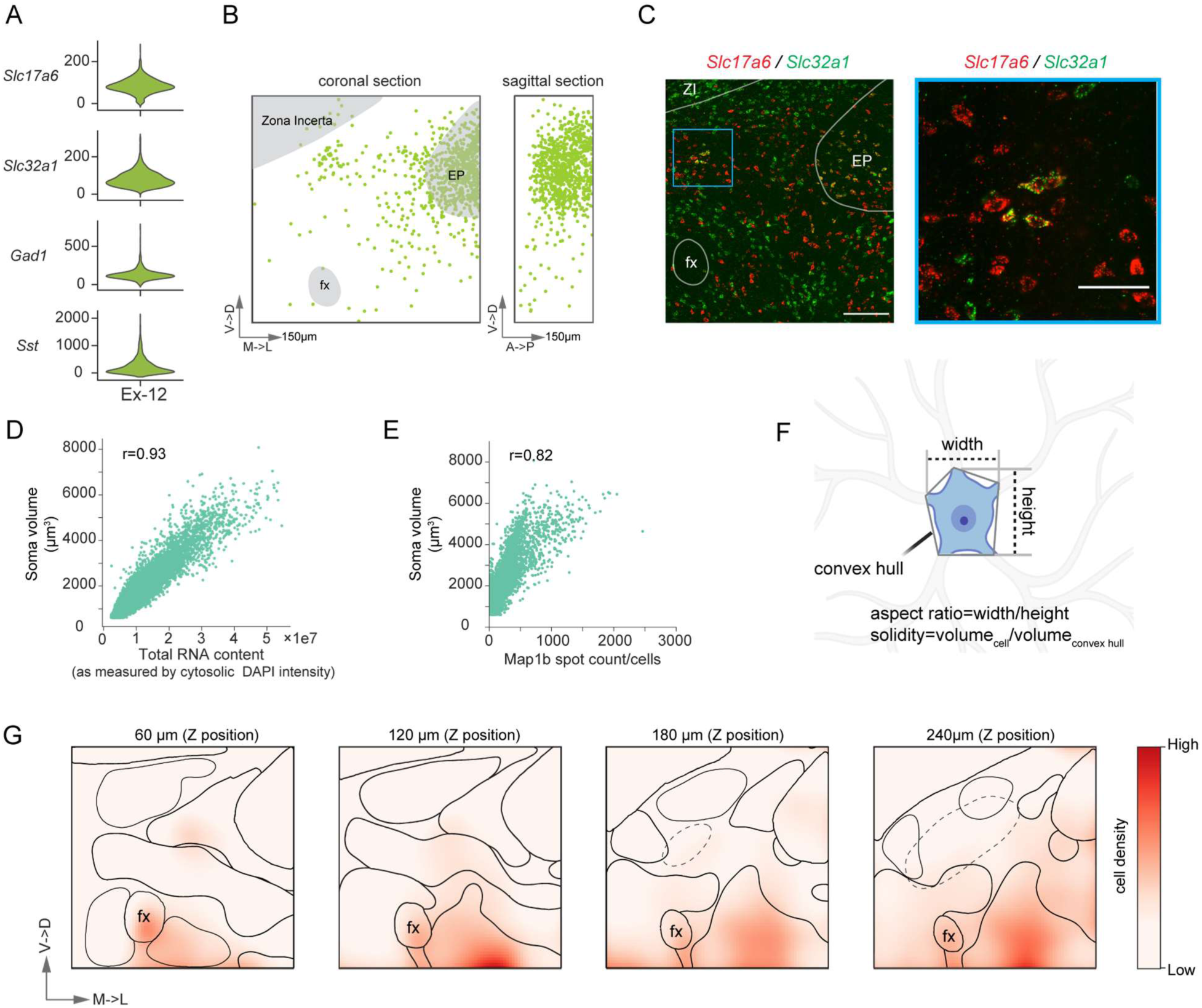
Morphological measurements of molecularly defined clusters, related to Figure 7. **(A)** Expression of selected genes in *Slc17a6* and *Slc32a1* co-expressing neurons. **(B)** Spatial distribution of *Slc17a6* and *Slc32a1* co-expressing cluster, Ex-12. Majority of these neurons are in the entopeduncular nucleus (EP), with a small cluster in the LHAd-db. **(C)** Representative image highlighting the *Slc17a6* and *Slc32a1* co-expressing neurons in the EP and LHA. Left: Representative single optical plane image showing *Slc17a6* and *Slc32a1* expression in the LHA. Scale bar: 100µm. Right: zoom-in of region highlighted in blue box (left) to show the *Slc17a6*/*Slc32a1* neurons in the LHA, Scale bar: 50µm. **(D)** Correlation between total RNA content and soma size. **(E)** Correlation between pan-neuronal marker, *Map1b* expression and soma size. **(F)** Schematic showing 3D morphological measurement (solidity and aspect ratios) performed in Figure 7A. **(G)** Density analysis of neurons with low convexity. Image slice of every 60 µm shown as representative images. From left to right: anterior to posterior (rostral→caudal). Scale arrow: 150 µm. Statistics: **Table S1**.

## Materials and methods

All methods for animal care and use were conducted according to National Institutes of Health guidelines for animal research and approved by the Institutional Animal Care and Use Committee (IACUC) at Janelia Research Campus.

### EASI-FISH protocol

#### 1) Reagents and chemicals

##### MelphaX

Melphalan (Cayman Chemicals) was dissolved to 2.5 mg/ml in anhydrous DMSO (Sigma). Acryloyl-X, SE (Thermo Fisher, 20770) was dissolved in anhydrous DMSO (10 mg/ml). 4 parts of Melphalan stock was combined with 1 part of Acryloyl-X, SE stock and reacted overnight with shaking at room temperature to make Melphalan-X (2 mg/ml). Aliquots were stored at −20 ℃ in a desiccated environment and used at 1 mg/ml by diluting in MOPS buffer (20 mM, pH 7.7).

##### Sodium Acrylate (4M)

The purity of commercial sodium acrylate was variable, so we made sodium acrylate by reacting acrylic acid (Sigma) with NaOH. Briefly, in a fume hood, acrylic acid (5.5 ml) was mixed with nuclease free water (4.5 ml). 10 M NaOH (7.2 ml) (Fisher) was added gradually to prevent excessive heating. Then 1M NaOH (Fisher) was added dropwise until the pH reached 7.6-7.8. Water was added to reach a final volume of 20 ml.

##### Stock-X

4M Sodium Acrylate (4.6 ml), 50% Acrylamide (w/v in water, 1 ml) (Sigma, A9099), 2% N, N’-Methylenebisacrylamide (1.5 ml) (Sigma, M7279), 5M NaCl (8 ml), 10 × PBS (2 ml), Nuclease Free Water (1.8 ml). Aliquots were stored at −20 ℃.

##### ExM gel solution

Before gelation, Stock-X was mixed with 0.5% 4-Hydroxy-TEMPO (Sigma, 176141), 10% TEMED (Sigma, T22500) and 10% APS (Sigma, A3678) at a ratio of 94:2:2:2.

##### Proteinase K digestion buffer

50 mM Tris-HCl (pH 8), 50mM NaCl, 1 mM EDTA, 0.5% Triton X-100 and 0.3% SDS.

##### DNase 1 Buffer

10 mM Tris-HCl (pH 8), 2.5 mM MgCl_2_, 0.5 mM CaCl_2_.

##### Poly-L-Lysine coating solution

Photo Flo 200 (3.2 µl, EMS 74257) was added to Poly-L-Lysine (1.6 ml. Pelco, 18026) to make the Poly-L-Lysine coating solution.

##### HCR hairpin conjugation with custom fluorophores

We custom conjugated the photostable fluorophore JF-669, NHS (Tocris, 6420) to amine-modified hairpin oligos from Molecular Instruments. Briefly, 100 mM amine-modified hairpin oligos (h1 and h2) (5 µl each) were air-dried with SpeedVac for 30 min. Dried oligos were dissolved in 0.1 M sodium bicarbonate (3 µl) (pH 8-9). JF-669, NHS (0.1 mg) was dissolved in DMSO (2 µl) and reacted to amine-modified oligos overnight in the dark at room temperature. Excess JF-669, NHS was removed with QIAquick Nucleotide removal kit (Qiagen, 28304) and the conjugated JF-669 hairpins were reconstituted with nuclease free water to a final concentration of 60 ng/ml (3 mM).

#### 2) Tissue fixation and preparation

C57Bl/6 male mice (8 weeks old) were used for all FISH experiments described in this study. Animals were anesthetized with isoflurane and perfused with RNase-free PBS (15ml) followed by ice-cold 4% paraformaldehyde (PFA) (50 ml). Brain tissue was dissected and fixed in 4% PFA overnight before sectioning on a vibratome. Brain coronal slices (300 µm) were sectioned and stored in 70% ethanol at 4℃ for up to 6 months. For the lateral hypothalamus experiment, LHA region (∼2.5 × 4 mm) around Bregma −1.355 to - 1.155 was cut out using anatomical landmarks as boundaries, including the mammillothalamic tract (mtt), zona incerta (ZI), fornix and optic tract. For ease of orientation and optimal imaging, the tissue was cut as a rectangle. An RNase-free paintbrush was used for tissue handling.

#### 3) RNA anchoring, gelation and Proteinase K digestion

The tissue slice was rehydrated in PBS at room temperature (RT) (2 × 15 min) and incubated in MOPS buffer (20 mM, pH 7.7, 30 min). Tissue was incubated overnight (37 ℃) in MOPS buffer (50 µl) with 1 mg/ml MelphaX and 0.1mg/ml Acryloyl-X, SE. The next day, tissue was rinsed in PBS (2 × 5 min) and placed in a 9 mm wide × 0.5 mm deep gasket (Invitrogen) on a glass slide that was previously coated with Poly-L-Lysine (1 µl) and allowed to dry. Gel solution was freshly made (see recipe above) and kept on ice. Tissue was equilibrated with gel solution (40 µl, 3 × 10 min) at 4 ℃. A coverslip was used to seal the gasket and gel was allowed to form at 37 ℃ for 2 hours. The coverslip and gasket were then removed to recover the tissue-gel. The tissue-gel was trimmed to a rectangle shape, with a corner cut to help with orientation. Tissue-gel samples were then transferred into a 2 ml Eppendorf tube and digested overnight (37 ℃) in Proteinase K buffer (750 µl) with 7.5 µl of 800U/ml Proteinase K (NEB, P8107S). After digestion samples were trimmed again and washed in PBS (4 × 15 min).

#### 4) DNase digestion

Tissue-gel samples were equilibrated in DNase buffer (750 ml) for 30 min and then incubated with RNase-free DNase1 (2.7 Kunitz units/µl, 50 µl) (Qiagen) in DNase buffer (450 µl) at 37℃ for 2 h. After DNase digestion, samples were washed in PBS (4 × 15 min) to remove DNase1. DNase 1 treatment (2 × 2 h) was performed prior to FISH to completely digest nuclear DNA.

#### 5) *In-situ* Hybridization and HCR

For hybridization, tissue-gel samples were first equilibrated in hybridization buffer (500 µl) for 30 min at 37 ℃. Samples were then hybridized with probe sets (10 nM each probe) in hybridization buffer (300 µl) overnight at 37 ℃. The next morning, samples were washed in probe wash buffer (2 × 15 min, then 3 × 30 min), followed by PBS (6 × 30 min) at 37 ℃. Samples were left in PBS overnight at room temperature for hybridization chain reaction (HCR) the next day or stored in 4 ℃. For HCR, samples were first equilibrated in amplification buffer (500 ml) for 30 min. Hairpins (3 mM) (conjugated to AF-488, AF-546, JF-669) were snap-cooled in a PCR thermocycler (95 ℃ for 90 seconds then cooled at room temperature for 30 min). To initiate HCR, hairpins h1 and h2 were mixed and diluted to 30 nM in 300 µl fresh amplification buffer. Samples were then left in the dark for HCR reaction at room temperature for 3 h. After HCR, samples were first washed in 5 × SSCT (5 × SSC + 0.1% Tween) (2 × 30 min), then washed in 0.5 × SSCT (0.5 × SSC + 0.1% Tween) (2 × 30 min) and stored in 0.5 × SSC at 4℃ before imaging. All HCR 3.0 probe and hairpin oligos were purchased from Molecular Instruments. *Gad1* HCR 3.0 probes used for assessing the detection efficiency of EASI-FISH were designed based on (Choi et al., 2018) and purchased from IDT.

#### 6) Image acquisition, sample handling and multiplexing

All samples were imaged on a Zeiss Lightsheet Z.1 microscope. A 20× water-immersion objective (RI=1.33) was used for imaging with 1× zoom. Single-side illumination was used to reduce light exposure and imaging time. Images were collected at 0.23 × 0.23 × 0.42 µm voxel resolution (post-expansion) with two tracks: the 488 nm and 669 nm channels were collected together followed by dual-track collection of 546 nm and 405 nm channels. For large volume imaging, each image tile was 1920 × 1920 pixels (438.5 µm × 438.5 µm, post-expansion) in size with around 1500 z-slices, with overlap between tiles set to 8%. For imaging of the lateral hypothalamus samples, zona incerta, the fornix and optic tract were used to guide selection of field of view (FOV). 4 × 4 tiles were taken from the LHA tissue-gel sample (usually 2-3mm in x and y dimension).

Before imaging, samples were stained in PBS with 5 µg/ml DAPI (2 × 30 min). At this concentration and in the absence of DNA, DAPI stains the RNA in the cytoplasm. Samples were mounted to a Poly-L-Lysine coated 8mm glass coverslip (Harvard Apparatus) that was glued to a custom-made plastic holder and imaged in PBS (Expansion factor: 2×). After image acquisition the probes and HCR products were removed using DNase1 (see above) with samples attached to the holder. To remove samples from the holder for re-hybridization, they were incubated at room temperature in 10% dextran sulphate (500 ml) (Fisher Scientific) in PBS for 30 min. A paintbrush can be used to assist with gel removal. Note that all units discussed in this paragraph regarding pixel size and image size were in post-expansion units.

We found that HCR spots were susceptible to light-induced fragmentation, producing mobile spots of reduced brightness (wigglers). Wigglers were difficult to detect without time-resolved imaging and led to false positive spots outside of cell bodies. Carefully controlling the light dose (reduced laser power and exposure time) alleviated light-induced fragmentation of HCR spots. Anti-fade compounds, such as PPD and DABCO also reduced spot fragmentation (**Table S2**). However, note that antifade compounds dramatically increased the bleaching rate of one of the commercially available HCR hairpin fluorophores (AF-546) (Figure S1H).

### EASI-FISH data processing

Large volumetric EASI-FISH datasets were collected on a Zeiss Lightsheet Z.1 microscope and saved as single multidimensional (multi-tile, multi-channel, z-stack) CZI files. For stitching, we also export a metadata file (MVL format) that includes tile configurations, which were later converted into a JSON format.

#### 1) Image Stitching

We used a previously developed flat-field correction and stitching package (Gao et al., 2019) that enabled rapid processing of 3D image tiles with Apache Spark based high performance computing environments (https://github.com/saalfeldlab/stitching-spark). The flat-field correction was based on corrected intensity distributions using regularized energy minimization (CIDRE) (Smith et al., 2015), where the flat-fields were calculated for each tile and channel independently and applied to each tile stack prior to stitching. The tiles were then stitched through translation to maximize cross-correlations in overlaps (Preibisch et al., 2009) from the DAPI channel. The same transformation was applied to the other three image channels. Stitched image volumes were then exported and saved into N5 format. This flat-field correction and stitching pipeline enabled rapid and automated data processing. All flat-field correction, stitching, and data export were executed on HHMI Janelia’s LSF computing cluster.

#### 2) Image Registration

After stitching, the DAPI channels were used again for registration of multi-round image volumes. To enable rapid and robust data processing, we developed a registration package that combined random sample consensus (RANSAC) (Fischler and Bolles, 1981) based feature matching with non-symmetric, diffeomorphic image registration, called Bigstream (https://github.com/GFleishman/bigstream). First, a Difference of Gaussian (DoG) filter was applied to 8 × 8 × 4 down sampled (1.84 × 1.84 × 1.68 µm) image pairs and local maxima above a selected threshold were selected as features, matched with RANSAC and affine transformed. After applying this global affine transformation to 4 × 4 × 2 down sampled (0.92 × 0.92 × 0.84 µm, post-expansion) image volumes, the transformed image volumes were split into 256 × 256 × 256 pixel-chunks with 12.5% spatial overlaps along each boundary for further processing. Another round of feature selection and affine transformation was performed on each image chunk at 0.92 × 0.92 × 0.84 µm (post-expansion) scale, followed by deformable registration. For better integration with the rest of the image processing pipeline, we created an implementation in Python of the non-symmetric diffeomorphic registration algorithm from the Greedy software package (Yushkevich, 2016). This enables in-memory data sharing of objects produced by different steps of the pipeline and thus avoids data saving in intermediate steps. It also guarantees compatibility of object formats such as transforms. (Yushkevich, 2016) All registration steps on image chunks can be executed in parallel. The global affine, piecewise affine, and piecewise deformable transforms were composed to a single displacement vector field stored in N5 format. Both forward transform and inverse transforms were computed. The forward transform maps on-grid-positions from the fixed coordinate system to corresponding (potentially off-grid) locations in the moving coordinate system and is required for resampling moving image data to match fixed image data. The inverse transform maps on-grid-positions in the moving coordinate system to (potentially off-grid) locations in the fixed coordinate system and is required for moving explicit positional data (such as detected FISH spot coordinates) from the moving coordinate system to the fixed coordinate system. For assessing the registration accuracy, mean structural similarity index (SSIM) (Wang et al., 2004) between the moving and fixed image was computed.

#### 3) FISH Spot detection

The spot detection approach was based on Airlocalize (Lionnet et al., 2011) and optimized for large image volumes (see below). Briefly, for Airlocalize, the images were first passed through a Difference of Gaussian (DoG) filter and global background subtracted. Next, a pre-detection step was performed that identifies local maxima above threshold within a 5-pixel radius, which was used to estimate the rough spot position. Then a local background correction was performed based on background estimation around each spot and a 3D Gaussian fit was used to estimate the spot location as well as the spot fluorescence intensity. Airlocalize was originally written in MATLAB and works best for epifluorescence and confocal image data (∼1 GB in size).

For large image data associated with the EASI-FISH data collection pipeline, we developed a high-throughput version to allow for distributed data access and processing. In order to integrate it with other parts of the EASI-FISH pipeline, we compiled Airlocalize (with minor modifications) into a Python package for distribution. Large image volumes were split into overlapping chunks (∼1GB in size, 10% overlap in z and 5% overlap in x and y). Spot detection on each chunk was performed simultaneously in parallel. The detected spots from individual image chunks were combined, with repetitive detections removed. Detected spot positions and intensities were exported as a single CSV file. All spot detection procedure was performed on full resolution images. Linear spectral unmixing was applied to images acquired in the red channel (AF546) to correct for bleed-through from the co-acquired DAPI channel before spot detection.

We called this Airlocalize-based high-throughput spot detection approach hAirlocalize, and it is available at https://github.com/multiFISH/EASI-FISH. After spot detection from each EASI-FISH round, the inverse transformation matrix acquired from the registration step (see above) was applied to the spot point cloud to transform spots to fixed image coordinate. Spot detection and warping were confirmed by visual inspections.

#### 4) 3D soma Segmentation

To achieve automatic and accurate detection and segmentation of cells in 3D image volumes, we developed a deep learning-based segmentation method based on the previously published 3D *StarDist* approach (Weigert et al., 2020). In a first step, *StarDist* predicts for each pixel its cell center probability and its radial distances to the nearest cell borders. In a second step, it selects the center point of each cell and uses the radial distances at this point to create a star-convex polyhedra that approximates the cell outlines. While this approach provides high detection accuracy in situations with many crowded and dense objects, the resulting segmentation masks are imprecise for non-convex cell shapes which could lead to unwanted misallocation of FISH spots. To address this problem, we retained the center probability and distance prediction step from *StarDist*, but afterwards aggregated pixel affinity maps from the densely predicted distances, an approach that we called *Starfinity*. The final segmentation masks were obtained by applying a watershed segmentation on the affinity maps while using the thresholded center probability as seeds. We trained such a *Starfinity* model with annotated cells from 3 different brain areas (LHA, CEA and cortex) using an iterative approach. Manual annotation was performed using Paintera (https://github.com/saalfeldlab/paintera). First, image stacks with 248 intact cells were manually annotated at full resolution (0.23 × 0.23 × 0.42 µm) to train a *Starfinity* model. Then from the prediction, we selected for images where cells were mistakenly segmented and corrected the errors in these images manually. We then feed the corrected annotations (827 cells) to train the model. For both training and predictions, images were down-sampled (4 × 4 × 2-fold) to increase the receptive field of the model, make the pixel resolution approximately isotropic, and reduce computational demands. The training and prediction were based on the cytosolic DAPI channel. The trained model gave a segmentation accuracy of 93%, with 4% over-segmentation errors, 1% under-segmentation errors and another 2% contaminated by neighboring cells.

After *Starfinity* segmentation, a semi-automated approach was used to further correct over-segmentation errors. Over-segmented ROI pairs were automatically flagged and merged based on the following criteria: 1) very high gene expression correlation between ROIs (Pearson Correlation Coefficient greater than 0.998, this cutoff was determined by maximum Youden’s index calculated on manually inspected data); 2) the centroid positions of the selected ROI pair were less than 23 µm (pre-expansion) apart and the two ROIs were touching/connected; 3) At least one of the ROIs had to be greater than 600 µm^3^ in size, to avoid merging of neuronal fragments with non-neurons or fragments from surrounding neuronal processes, which were usually small. For ROI pairs both greater than 1500 µm^3^ in size (average soma size in this region) and ROIs that had more than one matches, the corresponding ROI pairs were ranked based on their correlation coefficient and inspected manually (less than 15% out of all flagged ROI pairs). This method was estimated to eliminate 62% over-segmentations errors, bringing down the total over-segmentation error to less than 2%.

#### 5) End-to-end analysis pipeline

To enable easy adoption of EASI-FISH, we built a self-contained, highly distributed, and platform agnostic computational pipeline for analyzing EASI-FISH data. The pipeline was built using Nextflow, with all software containerized, which makes it portable and reproducible. The pipeline can be executed out-of-the box on multiple platforms, including stand-alone workstations and the IBM Load Sharing Facility (LSF) computing cluster. It could also be adapted to execute on other batch schedulers (such as SLURM) and on cloud platforms (e.g. Amazon Web Services, Google Cloud).

The pipeline is also highly flexible. We provide an end-to-end pipeline for EASI-FISH users to analyze their imaging data and directly translate over 10 terabytes of image data into spot counts per cell. The EASI-FISH steps (stitching, registration, segmentation, spot detection, etc.) in the pipeline are modularized, allowing flexibility for users to select only the necessary modules for their specific application. Furthermore, each module is containerized using Docker, and can therefore be easily substituted with a different implementation or algorithm.

The pipeline is openly available at https://github.com/JaneliaSciComp/multifish and includes extensive documentation and automatic download of example data sets for push-button replication. Monitoring of task execution and resource utilization is available through the use of the Nextflow Tower web UI.

For analysis of an example dataset (35GB, 4 channels, 600 z-slices for each EASI-FISH round), we provide the processing time below for each step on a stand-alone workstation (128GB RAM, 40 cores). For data that is larger than 100GB, we recommend usage of cluster or cloud for improved data parallelization and speed.

|  | image scale | Stand-alone workstation<br>(8h total) | LSF cluster<br>(3h total) |
| --- | --- | --- | --- |
| Stitching | full resolution | 2.5h | 1.5h |
| Registration | $4 \times 4 \times 2$ or<br>$8 \times 8 \times 4$ ds | 0.5h | 1.5h |
| Spot detection | full resolution | 7.5h<br>(2.5h/channel) | 1.5h |
| Segmentation | $4 \times 4 \times 2$ ds | 0.2h | 0.2h |
| Post-processing | -- | 0.05h | 0.05h |

Although this breakdown shows the time for each step, it is important to note that, once the stitching step is finished, the subsequent steps (registration, segmentation, and spot detection) can be performed simultaneously in parallel. The pipeline can be used to analyze multi-round EASI-FISH experiment all at once. Users can also choose to interleave the data collection and analysis with minimal modification. This way, the time lag between data acquisition and analysis will be negligible and users can start running through the pipeline once two rounds of EASI-FISH data have been acquired.

#### 6) Visualization

The stitched and registered volumetric image data was visualized with Fiji plugin N5 Viewer (https://github.com/saalfeldlab/n5-viewer) based on BigDataViewer (Pietzsch et al., 2015) for interactive browsing of multichannel multiscale N5 datasets. More flexible visualization options, including overlaying with segmentation mask, inspection of detected spots, ROIs and spots queries were performed using Napari (https://github.com/napari/napari) with custom scripting in python.

#### 7) Post-processing

With the segmentation masks from the DAPI channel and spot extracted from FISH channels, extracted spots from each FISH channel were then assigned to individual ROI to obtain the spot count for each FISH channel (gene). We also performed the following steps for improved data quality.

##### a. Filter ROIs

Cells on the edge of the field of view that were only partially captured in 3D or failed to be captured in one or more EASI-FISH image rounds were removed from the analysis. Additionally, we observed autofluorescence in the red channel on the surface of the specimen in later rounds (>7) and excluded part of the specimen (∼30-50µm pre-expansion, from the top surface along the z-axis) from the downstream analysis. After this procedure, 77% of detected ROIs were used for the following analysis (**Table S6**).

##### b. Lipofuscin spots

We observed autofluorescence puncta-like signals in the tissue-gel samples that are likely lipofuscins. Lipofuscins are lysosomal storage bodies, and their presence could lead to false positive spot detection. We took advantage of the fact that lipofuscins had a broad excitation/emission spectrum and can therefore be identified by signal colocalization in more than one channels. In the current application, the 488 nm and 669 nm channels were acquired at the same time and were used to identify lipofuscin spots by signal colocalization. We identified and spots whose centroid positions were within a radius of 3 pixels (345 nm, pre-expansion) between the 488 nm and 647 nm channels and subtracted these spots for all FISH channels. For genes with high expression levels (spot count exceeding 200 per cell, detected in round 4 to round 10), we chose to use the median lipofuscin spot counts for that cell across all rounds to avoid subtraction of real spots due to high spot density in both channels.

##### c. Dense spots

For highly expressed genes, the integrated fluorescence intensity was used instead to estimate spot counts. First, we analyzed the distribution of spot fluorescence intensity from the hAirlocalize output for each gene. The spot intensity followed a right-skewed distribution. Outliers with high intensity values were likely detection of multiple spots in cells with high expression. The mode of the distribution was used as an estimate for single spot intensity for any given gene. Spot counts in any given cell was calculated by dividing the sum of fluorescence intensity in that cell with the estimated single spot intensity. We compared the estimated spot counts with hAirlocalize spot counts for genes that showed low or medium expression levels (less than 200 spots/cell) in the majority of cells and that the two measurements were highly correlated and comparable in the low to medium spot density range. Figure S2F showed *Slc32a1* as an example, the least-square linear regression fitting to the data indicates a slope of 1.03 and R^2^ of 0.93. Because intensity-estimated spot counts can be sensitive to background fluorescence in cells with low gene expression, we chose to use hAirlocalize measurement in these cells. And for cells with spot density higher than 0.01spot/voxel (corresponding to spot separation less than 3.3µm apart), we applied the intensity-estimated spot counts.

##### d. Neuronal morphological analysis

Taking advantage of the accurate 3D segmentation, we measured the morphological properties of neuronal cell body (soma) with segmentation mask using scikit-image implementation of *regionprops*. The spatial position of each neuron was defined as the centroid coordinate of the segmentation mask. Solidity was a measure of the overall concavity and was defined as the neuronal cell body volume divided by its convex hull. The aspect ratio was defined as the length ratio between the major axis and the minor axis of the neuron. Other measurements that were not being discussed here, but have been performed include major axis length, minor axis length, neuronal density around each neuron, orientation, extent and whether a neuron is in direct contact with a non-neuron.

All measurements reported here were in pre-expansion unit in µm, unless otherwise noted.

### Single-cell RNA sequencing

The single-cell RNA sequencing was focused on the suprafornical LHA and its surrounding areas. For single cell dissociation and collection, we used transgenic animal *Agrp-IRES-Cr*e × *Ai9* crosses, where the tdTomato-labeled AGRP neurons extend a prominent set of axonal projections to the suprafornical region of the LHA (Betley et al., 2013) and provide a useful signal for visually guided dissection of the targeting brain region. The manual sorting procedure to isolate non-fluorescent cell bodies from micro-dissected brain slices was similar to previously described (Hempel et al., 2007). Briefly, we sectioned 300 μm coronal slices from male *Agrp-IRES-Cr*e × *Ai9* mice (age 6-8 weeks) and used the tdTomoato fluorescence signal from AGRP neuron axon bundle terminal in the LHA to identify the boundaries of the LHA and then manually dissected with spring scissors. The dissected tissue sections were then subject to protease digestion, after which cells were dissociated. Dissociated neurons from 7 animals were pooled and intact neurons were manually selected into individual wells based on size. Sorted single cell was lysed with 3 µl lysis buffer (0.2% Triton X-100 (Sigma) and 0.1 U/µl RNase inhibitor (Lucigen)) and cDNA libraries were prepared using the Smart-SCRB chemistry as described previously (Cembrowski et al., 2018; Xu et al., 2020). Barcoded cDNA libraries were then pooled and sequenced on a NextSeq 550 high-output flowcell with 27 bp in read 1 to obtain the barcode and UMI, and 125 bp in read 2 for cDNA. PhiX control library (Illumina) was spiked in at a final concentration of 15% to improve color balance in read 1. Libraries were sequenced to an average depth of 135,025 ± 38401 (mean ± S.D.) reads per cell.

Sequencing alignment was performed similar to (Gur et al., 2020). Sequencing adapters were trimmed from the sequencing reads with Cutadapt v2.10 (Martin, 2011) prior to alignment with STAR v2.7.5c (Dobin et al., 2013) to the M. musculus GRCm38.90. genome assembly from Ensembl (ensembl.org). Gene counts were generated using the STARsolo algorithm (https://github.com/alexdobin/STAR/blob/master/docs/STARsolo.md). Gene counts for the subset of barcodes used in each library were extracted using custom R scripts.

### scRNA-Seq analysis and marker-gene selection for EASI-FISH

Analysis of the single-cell RNA sequencing data, including filtering, variable gene selection, dimensionality reduction and clustering was performed with Seurat (v2.3.4) (Butler et al., 2018; Satija et al., 2015) in R (v3.4.3). First, cell doublets/multiplets and low-quality cells were filtered based on the total number of detected genes (1,500-7,500), relative abundance of mitochondrial transcripts (percent.mito < 0.055) and number of unique molecular identifiers (nUMI) per cell (< 2 × 10^5^) respectively. Genes expressed in less than 3 cells were also removed. The resulting dataset consisted of 1507 cells and 17535 genes. The filtered dataset was then log-normalized and scaled, while regressing out the effects of latent variables including nUMI, and percent.mito. Next, we performed principal component analysis (PCA) and used the first 31 principal components for downstream analysis. For clustering, we used the graph-based clustering approach implemented in Seurat, with the original Louvain algorithm and 10 iterations. Non-neuronal cells were identified and removed from the dataset before further analysis.

The remaining 1,425 cells were processed similar to what was described above and 4 neuronal cell types were identified, whose preliminary identities were assigned based on unique expression of enriched genes: Group 1 (70%, 1043 cells) consisted primarily of cells with high levels of the neuropeptide *Hcrt*, Group 2 (15%, 227 cells) was a heterogenous population best characterized by common expression of *Sparcl1*, Group 3 (10%, 151 cells) contained a high percentage of cell strongly expressing *Nts*, and Group 4 (4.5%, 67) was defined by very high levels of *Pmch*. Assessment of differential gene expression between neuronal cell types was performed using the FindAllMarkers function in Seurat (Wilcoxon rank sum test, logfc.threshold = 0.55, min.pct = 0.25), with p-values adjusted based on the Bonferroni correction. The full list of enriched genes for each major neuronal subclass is provided in **Table S4**.

To identify marker-genes for EASI-FISH, we started with the list of differentially expressed genes as outlined in **Table S4**. We applied a series of selection criteria designed to allow classification of a maximum number of unique cell types using the fewest number of genes possible. As such, in addition to limiting our search to genes with an adjusted p-value cutoff of at least 0.05 and an average log-fold change of 0.55 or over, we also specifically selected markers with as close to binary “on/off” expression patterns in the cell type of interest as possible, based on high percentage of marker positive cells in the target population compared to low percentage of marker positive cells outside the target population (displayed as pt.1 and pt.2 in **Table S4**, respectively). While not explicitly used as a limiting factor for the selection of marker-genes in this experiment, we found that a value of 0.4 for the ratio of minimum difference in the fraction of detection between the two groups to be an informative rubric for aiding in selection of marker-genes with close to binary characteristics. We also looked for genes that could further split the HCRT and PMCH neurons as they are well known neuropeptide secreting neurons with important functions. Using these parameters, alongside manual inspection of the Allen brain ISH atlas (Lein et al., 2007) for cross-validation, we settled on the following genes to represent the neuronal cell types of greatest value and highest confidence given the number of assessed cells in the scRNAseq dataset: *Hcrt*, *Bdnf*, *Calb2*, *Nts*, *Gpr83*, *Pmch*, *Cartpt*, *Tac2*. Additionally, *Slc17a6* and *Slc32a1* were included to specify excitatory and inhibitory neurons respectively and *Map1b* was used as a pan-LHA neuronal marker. Due to the over-representation of specific neuronal cell types (e.g HCRT neurons) and under-representation of some neuronal cell types (e.g inhibitory neurons) in this dataset (likely due to bias during hand sorting), we chose to supplement the marker list with those identified in a recently published dataset (Mickelsen et al., 2019). Additional genes were selected for inclusion in an effort to represent a broader diversity of cell types in the LHA. The collected final list of marker-genes are listed in **Table S5**. We acknowledge that this is not the only combination of genes that could feasibly serve to represent these molecularly defined cell types.

### Integration of scRNA-Seq datasets

In order to obtain a broader diversity of molecularly defined cell types in the LHA, we integrated our scRNA-Seq data with published LHA scRNA-Seq datasets (Mickelsen et al., 2019; Rossi et al., 2019). Processed gene count expression matrices were directly downloaded from Gene Expression Omnibus (GEO). For the Mickelsen et al. dataset, female and male mice were combined. For the Rossi et al. dataset, only control groups were included and combined. Integration of multiple scRNA-Seq datasets was performed using Seurat (v3.2.0) (Stuart et al., 2019). First of all, similar to described above, data were filtered based on the number of features and total RNA counts to eliminate doublets and cells with low quality. Genes that were expressed in less than 5 cells in the dataset were removed from the analysis. Non-neurons were then removed from the three datasets before data integration based on highly variable features. 4,418 cells from the Mickelsen et al. dataset and 2,087 cells from Rossi et al. dataset were included for integration with our dataset. Clustering analysis was performed similar to described above and the Silhouette score was used to determine the optimal resolution for cluster numbers. To identify the optimal k parameter (neighborhood size) for clustering, we ran a bootstrap analysis by randomly selecting 80% of cells 100 times and performing the analysis described above. The Jaccard similarity coefficient was used for evaluation. The bootstrap analysis was performed in R using the *scclusteval* package (Tang et al., 2020) with modifications.

### Assessment of EASI-FISH method

For assessment of detection efficiency, *Gad1* was labelled with interleaved probe sets conjugated to different fluorophores (10 probes in each set). Assuming probe independence from the two probe sets, the detection efficiency with 10 probes/set can be measured as the square root of colocalization efficiency. We found that 65 ± 2 % spots colocalized when we probe Gad1 with two independent probe sets, corresponding to 81 ± 1.4% detection efficiency.

To test detection sensitivity of the EASI-FISH method, we focused on a single neuronal cell type from the scRNA-Seq dataset, the PMCH neurons. We identified genes that are selectively expressed in this population at various levels (**Table S4**). *Klhl13* (UMI_mean_=48) and *Igf1* (UMI_mean_=15) were selected specifically due to their unique high levels of expression in most of PMCH neurons (high pct.1) and relative absence in other cell types (low pct.2), as well as their absolute low levels of gene expression as indicated by UMI counts.

### FISH cluster analysis

First, neurons were identified using a two-components Gaussian Mixture Model (GMM) based on spot count of LHA pan-neuronal marker, *Map1b* with probability greater than 0.7. 36,423 out of 66,488 cells were identified as neurons in this way. Clustering analysis was performed on z-score normalized spot counts of 24 marker-genes. Unlike the scRNA-Seq analysis, no logarithmic transformation was applied to minimize the weight of false positive spot detection. Similar to described above, principal component analysis (PCA) was performed, and graph based SNN clustering analysis implemented in Seurat was used for an initial clustering using all marker-genes. Like the scRNA-Seq analysis, this separated neurons into *Slc17a6* and *Slc32a1* populations. Subsequent clustering was performed on *Slc17a6* and *Slc32a1* populations separately. For *Slc17a6* population, *Map1b* and *Slc17a6* were excluded from the PCA and clustering analysis as they were not considered variables. For *Slc32a1* population, non-variable genes (widely expressed: *Map1b, Slc32a1*; not expressed: *Bdnf*, *Cartpt*, *Trh*, *Pmch*, *Hcrt*) were excluded from PCA and clustering analysis. The Silhouette score, SC3 stability index and Jaccard similarity coefficient as calculated by bootstrapping 80% of the data 100 times (same as above) were used to choose clustering parameters: the nearest neighbourhood size k and resolution to maximize the number of stable clusters (k parameter chosen for *Slc17a6* cells: 20, k parameter chosen for *Slc32a1* cells: 10). Cell types with averaged total spot counts from all genes below 20 percentiles of all neurons were aggregated and assigned as the poorly classified cluster due to lack of marker-genes. Using this cutoff, 80% of neurons from the *Slc17a6* population (Ex-1 to Ex-24) and 74.5% of neurons from the *Slc32a1* population (Inh-1 to Inh-22) were classified. tSNE was used for visualization with perplexity of 50 for *Slc17a6*^+^ population and 40 for *Slc32a1*^+^ population.

### LHA boundaries and neighboring brain regions

The zona incerta (ZI) is characterized by high density inhibitory neurons with small and regular cell bodies (Kawana and Watanabe, 1981; Kolmac and Mitrofanis, 1999). Therefore, we drew boundaries between the ZI and the LHA based on neuronal density, morphology, as well as *Slc17a6* and *Slc32a1* expression. The entopeduncular nucleus (EP) is enriched for glutamate/GABA co-releasing somatostatin neurons (Wallace et al., 2017), which were used to identify boundaries for EP. The fornix was identified based on its location, circular profile, and lack of cell bodies.

### Spatial mixing analysis of molecularly defined cell types

The neighborhood complexity and purity analysis were similar to what has been described previously (Moffitt et al., 2018). Briefly, the neighborhood complexity was defined as the number of distinct other molecularly defined cell types within the neighborhood of any given neuron. The neighborhood purity was defined as the fraction of neurons within the neighborhood that were part from the most abundant molecularly defined cell type. A 50 µm radius surrounding any given neuron was used as the neighborhood for this analysis.

### Regional enrichment of selected marker-genes

To compute the selective regional enrichment for *Otp*/*Meis2*, we first counted the number of *Otp^+^* and *Meis2^+^* neurons within a 50 µm radius neighborhood of any given neuron. The selectivity index for *Otp*/*Meis2* is calculated as (*Otp^+^_num_ – Meis2^+^_num_) /* (*Otp^+^_num_ + Meis2^+^_num_).* For *Slc17a6*/*Slc32a1*, the same procedure was performed based on the number of *Slc17a6^+^* and *Slc32a1^+^* neurons in the neighborhood, except for cells that co-expressed *Slc17a6* and *Slc32a1*, which were excluded from the analysis. This procedure was advantageous compared to an image smoothing filter as it preserved the neuronal density information, which was useful for segmentation later.

### Automatic segmentation of the LHA

The initial segmentation of the LHA was based on expression of two pairs of genes *Otp*/*Meis2* and *Slc17a6*/*Slc32a1*. We first classified neurons into 4 classes based on their regional enrichment for these four genes (Class 1: *Slc17a6*^+^/*Otp*^+^; Class 2: *Slc32a1*^+^/*Otp*^+^; Class 3: *Slc17a6*^+^/*Meis2*^+^; Class 4: *Slc32a1*^+^/*Meis2*^+^) and then used Gaussian Mixture Models (n components=50) implemented in scikit-learn to generate the 3D segmentation (1µm isotropic voxel resolution). This segmentation was performed separately on each LHA volume. Then, the segmentation masks from LHA1 and LHA2 were aligned to LHA3 using the rigid registration implemented in Greedy (Yushkevich, 2016). The fiducial landmarks (ZI, EP and fornix) and spatial distribution of marker-genes from three animals were used to cross validate the registration. The simultaneous truth and performance level estimation (STAPLE) (Warfield et al., 2004) implemented in SimpleITK was used to generate a unified atlas with a probability threshold of more than 0.9. Disconnected regions that were from the same molecularly defined class (1-4) were further split and assigned unique ROIs. Small ROIs (< 0.5 × 10^6^ µm^3^) were removed and the segmentation boundaries were smoothened with a Gaussian filter (sigma=30). Detailed segmentation procedures can be found at https://github.com/multiFISH/LHA_analysis.

### Average distance to nearest neighbor (ANN) analysis

The nearest neighbor of each neuron was queried using kdtree (scipy) and the average distance to the nearest neighbor in each molecularly defined cell type was defined as the mean distance of neurons in a given cell type to its nearest neighbor. Note that neuron number in each molecularly defined cell type is different, therefore it is not a one-to-one relationship (e.g. multiple neurons can have the same nearest neighbor), and the ANN between cell type A and cell type B can be different depending on which cell type was queried.

### Spatial distribution of molecularly defined cell types

First, to determine the spatial distribution of molecularly defined cell types (whether they were clustered, dispersed or uniformly distributed), we compared the cell types with a CSR (complete spatial randomness) process and performed a Monte Carlo test of CSR (Cressie; Waller). We simulated the CSR process by randomly sampling cells in the data 1,000 times to generate a distribution of the averaged distance to nearest neighbor under CSR (ANN_CSR_). The number of random sampled cells was matched to that in each molecularly defined cell type. The ANN from each molecularly defined cell types (ANN_Mol_) was calculated and compared to the CSR distribution to calculate the p-value. We found that all (48/48) of the molecularly defined cell types were spatially clustered (ANN_Mol_<ANN_CSR_, p<0.05) compared to a CSR process (**Table S1**).

Second, to determine whether the molecularly defined cell types were enriched within proposed subregions, we used an approach similar to the Quadrat statistic (Cressie; Waller), instead of quadrat, the proposed anatomical parcellations were used for this analysis. One hypothesis was that the unequal distributions of molecularly defined cell types within proposed LHA subregions was due to differences in cell/point densities in these subregions. To test this, we simulated the distribution by shuffling neurons’ molecular identity 1000 times to compute the distribution of the χ ^2^ statistics for each cell type. The χ ^2^ statistic from the observed molecularly defined cell types was compared to the distribution of expected χ ^2^ statistics under the above hypothesis to calculate the p values (**Table S1**).

Last, to determine which subregion the given molecularly defined cluster was enriched in, we performed the permutation test, where we shuffled the position of neurons from each molecularly defined cell type 1,000 times and calculated the distribution of regional enrichment for any given molecularly defined cell type. The observed fraction of neurons enriched in a given subregion from each molecularly defined cell type was compared to the expected distribution from the random process to calculate the p values (**Table S7**).

### Fractional overlap analysis of molecularly defined cell types

To determine the molecularly defined cluster enrichment, we first calculated the local neuronal density of each molecularly defined cell type using kernel density estimation (kde) (scipy). The segmentation mask for each molecularly defined cell type was generated by thresholding the resulting density at the 95th percentiles. The binarized segmentation masks can be combined to generate a pixelated image with ‘intensity values’ at each voxel representing the number of overlapping cell types. This overlaid image was used to further subdivide the LHAfl and LHAs-db subregions. The segmentation masks generated for each molecularly defined neuronal type was used to calculate the fractional overlap between pairs of molecularly defined clusters. The overlap fraction was defined as the number of pixels occupied by both cell types divided by the total number of pixels occupied by either cell type.

### Prediction of spatial position with gene expression

To determine whether neuronal spatial position could be predicted based on combinatorial expression of marker-genes, we trained a multi-output Random Forest regressor to predict the spatial positions of neurons. First of all, gene expression data were normalized (z-score) to a normal distribution with similar scale ranges for improved model performance. We trained the random forest regressor using the z-score normalized expression of 24 marker-genes as input and the x, y, z positions (pre-expansion, in µm) as output. The prediction performance was evaluated using 10-fold cross-validation and random permutation cross-validation (Shuffle & Split). Both coefficient of determination (R^2^) and the explained variance regression score were calculated, and they were identical when rounded to 2 decimal places. For absolute measurement, the predicted position using this model can be, on average, ± 105 µm off from the true position. To evaluate the importance of individual genes and their combinations in predicting neuronal position, models were trained and tested by removing one or combinations of features (genes) and same as described above, cross validation was used to evaluate the predictive power of the model based on R^2^ score. To evaluate the statistical significance, we shuffled the relationship between the gene expression and neuronal position 1000 times and compared the prediction accuracy of our model to the distribution of prediction accuracy from shuffled data to calculate the p value.

### Cross-correlation analysis between biological replicates

To determine whether findings presented here were reproducible across animals, we performed cross-correlation analysis on 1) molecularly defined cell type gene expression (Fig S4F), 2) marker-gene spatial expression (Fig S5E), 3) molecularly defined cell type spatial distribution (Fig S5H) between biological replicates. For molecularly defined cell type gene expression, averaged marker-gene expressions in any given molecularly defined cell type were used to calculate the correlation coefficient. For the spatial distribution of marker-genes and molecularly defined cell types, correlation analyses were performed on tissue volumes after rigid alignment. For gene expression spatial distribution, gene expression per cell were binarized and cells with spot count greater than 30 were defined as positive for any given gene. The image volumes from different animals were discretized into 100µm × 100µm × 100µm bins and number of positive cells for any given gene was used to generate the correlation coefficient. For molecularly defined cell type distribution, instead of number of positive cells for any given gene, number of cells from any given molecularly defined cluster in each bin was used for cross-correlation. All correlation analyses were performed by comparing LHA1 and LHA2 to LHA3 and p value < 0.05 were reported here.

### Connectivity analysis

To determine whether neuronal inputs to the LHA follow the regional parcellation, we looked at dataset collected from the Allen Mouse Brain Connectivity database. We utilized the spatial search function from the Allen Brain Atlas API and selected for experiments meeting the following criteria: 1) rAAV-EGFP tracing in wildtype animals for broad neuronal cell type tracing; 2) density of projection signal (ratio of thresholded fluorescence pixel over all pixels in the structure) greater than 0.1 in the LHA (6350, 5850, 6850 in the reference space was chosen as the LHA target location). 148 experiments met these criteria. Affine alignment was applied to transform the LHA parcellation map to the reference atlas based on fiducial landmarks (ZI, EP and fornix). Mean fluorescence intensities in each LHA subregions were extracted from the 148 experiments. For multiple experiments from the same injection site, cross-correlation analysis was performed to remove outliner experiments, which is mostly likely due to variations in injection site. Additionally, experiments with injection site in the LHA and its neighboring brain regions (AHN, VMH) were removed as there could be strong signal spread from the injection site to the LHA. This allowed us to examine a total of 64 different brain regions that projects to the selected LHA target location and their relative projection intensities in proposed LHA subregions.

### Statistics

Statistical analyses were performed in Python or Prism GraphPad and described in method, figure legends and **Table S1**.

### Figures and scales

All scales (scale bar and scale arrows) in the text, figures and figure legends were converted to pre-expansion unit in µm.

