## Supplemental Information_Figures&Tables for "Expansion-Assisted Iterative-FISH defines lateral hypothalamus spatio-molecular organization"

**movie 1** Anti-fade eliminates laser induced mobile spots (provided as a separate file). Representative time-lapse videos (1 frame/second) showing laser induced mobile spots in EASI-FISH samples with or without anti-fade. Left: EASI-FISH sample imaged in PBS. Right: EASI-FISH sample imaged in antifade, p-phenylenediamine (PPD).

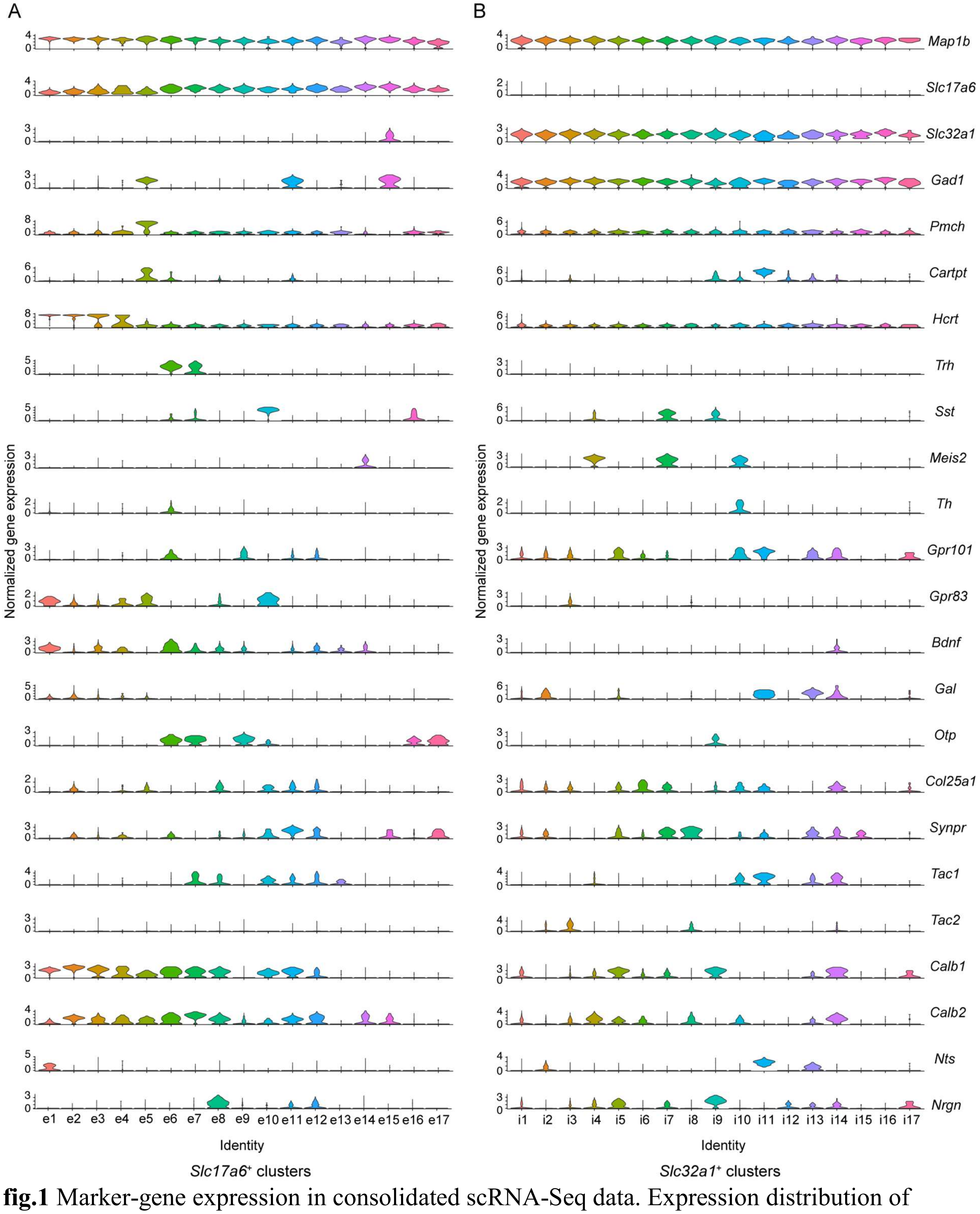


**fig.1** Marker-gene expression in consolidated scRNA-Seq data. Expression distribution of marker-genes in identified *Slc17a6*$^{+}$ (A) and *Slc32a1*$^{+}$(B) scRNA-Seq clusters. Normalized gene expression is calculated as the log-transformed and Z-score normalized UMI counts.

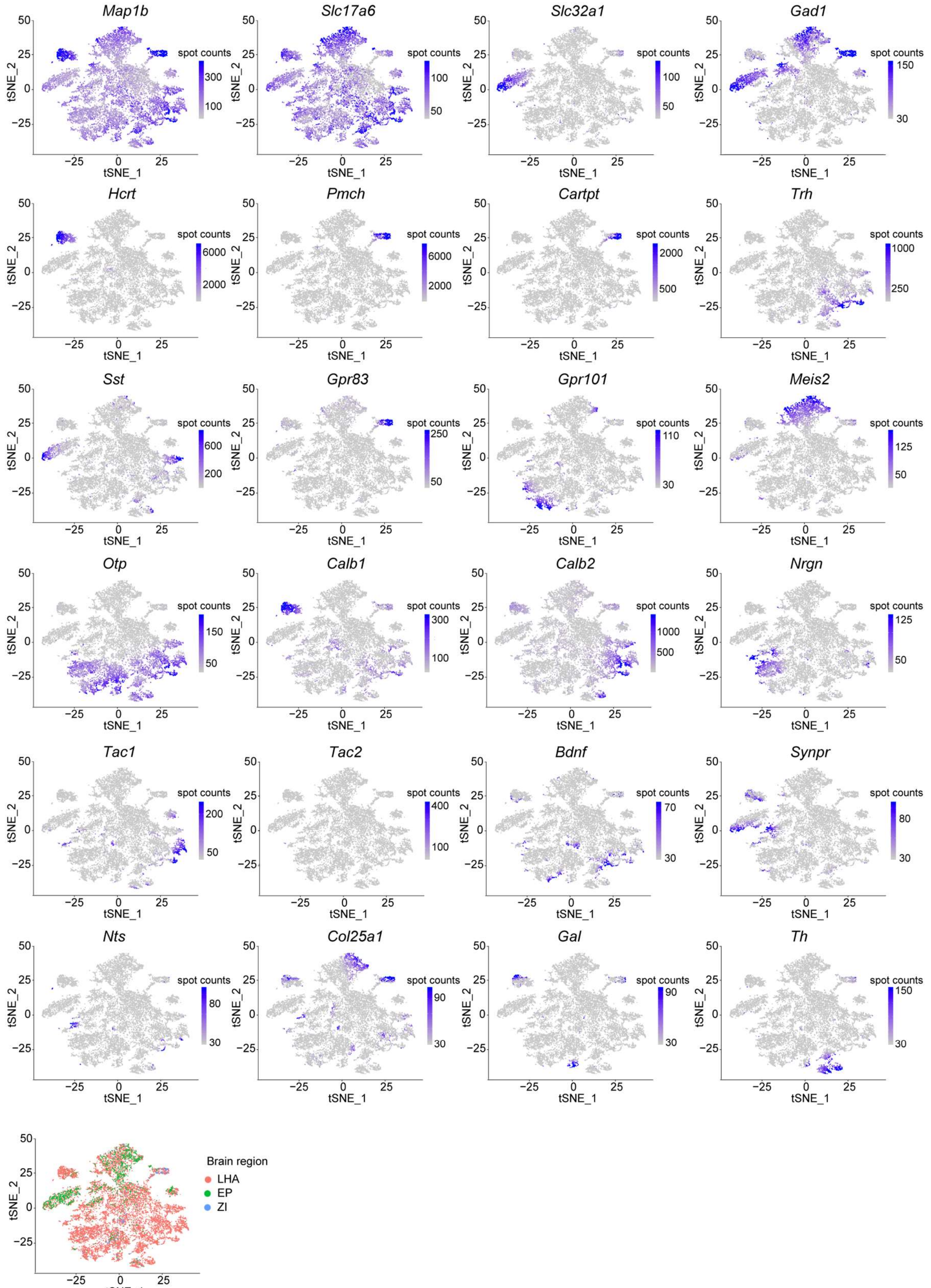


**fig.2** Marker-gene expression (spot/transcript counts) in *Slc17a6*$^{+}$ neurons as measured by EASI-FISH.

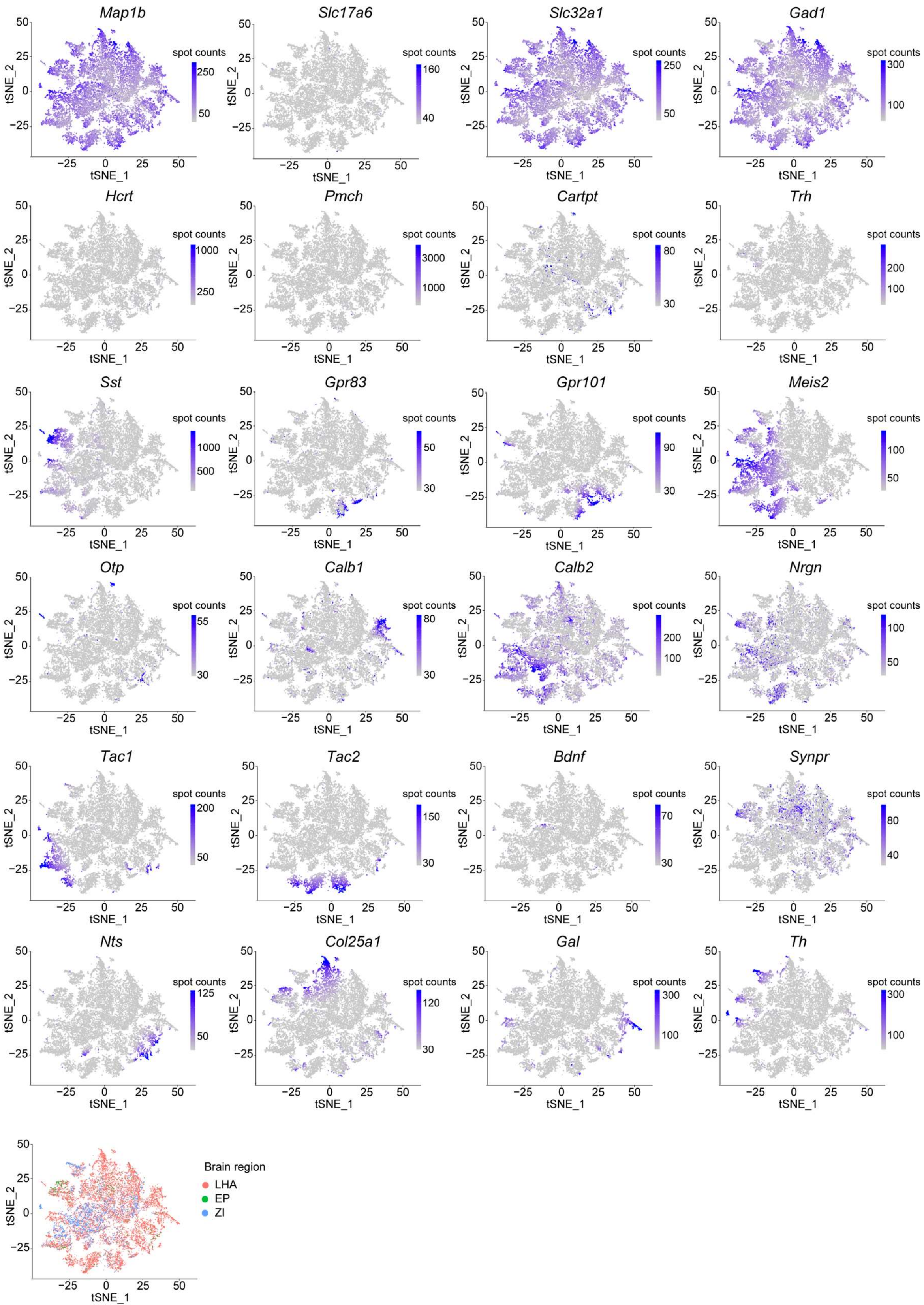


**fig.3** Marker-gene expression (spot/transcript counts) in *Slc32a1*$^{+}$ neurons as measured by EASI-FISH.

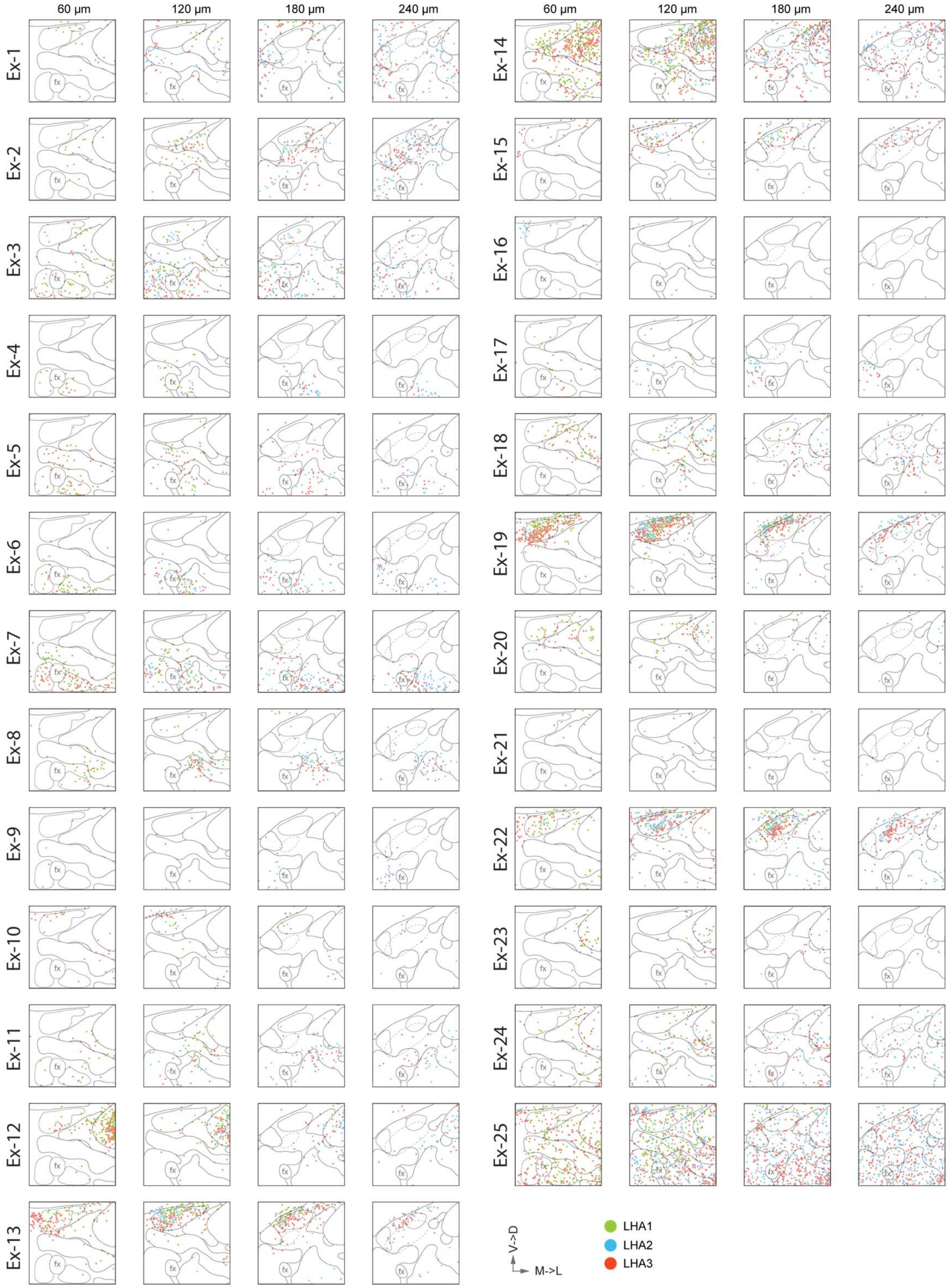


**fig.4** Spatial distribution of *Slc17a6*$^{+}$ clusters. Each dot indicates the centroid position of a neuron. Colors indicate neurons from different animals. Panels are 60 µm axial projections of sub-volumes. From left to right: anterior to posterior (rostral→caudal). Scale arrows: 150 µm. M→L: medial to lateral, V→D: ventral to dorsal.

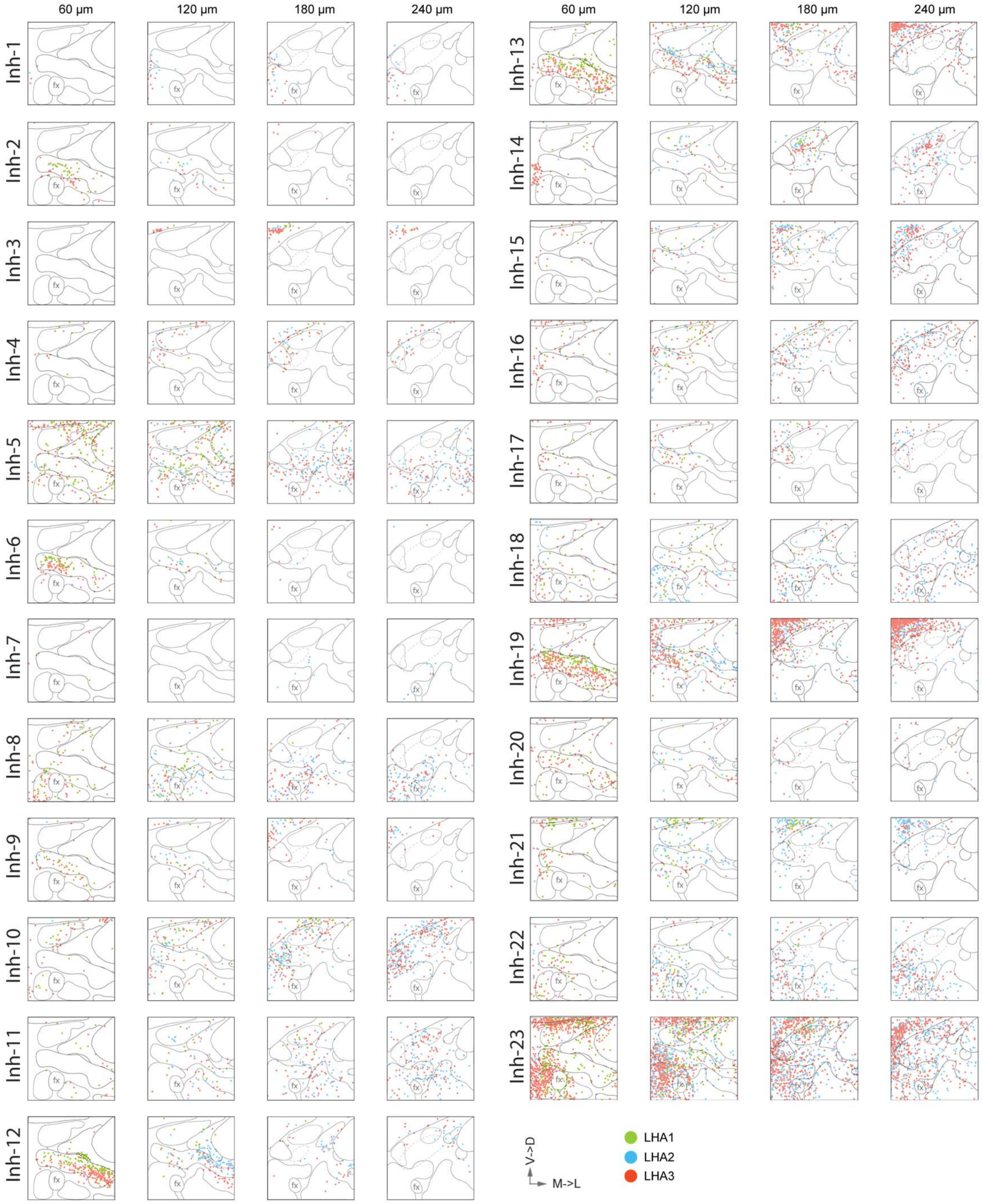


**fig.5** Spatial distribution of *Slc32a1*$^{+}$ clusters. Each dot indicates the centroid position of a neuron. Colors indicate neurons from different animals. Panels are 60 µm axial projections of sub-volumes. From left to right: anterior to posterior (rostral→caudal). Scale arrows: 150 µm. M→L: medial to lateral, V→D: ventral to dorsal.

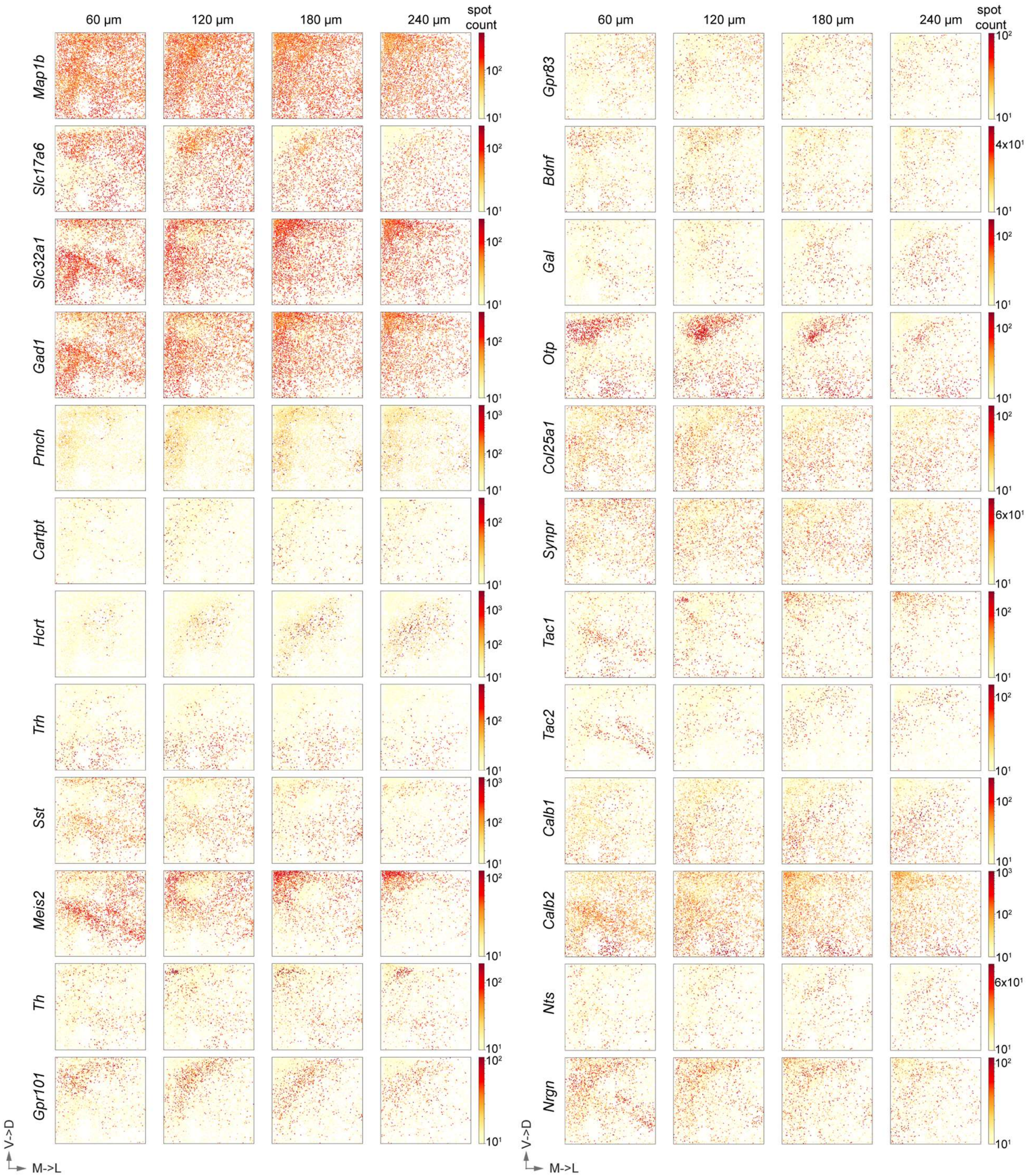


**fig.6** Spatial distribution of marker-gene expression in the LHA. Each dot indicates the centroid position of a neuron. Colors indicate the gene expression levels as measured by spot counts per cell. Panels are 60 µm axial projections of sub-volumes. From left to right: anterior to posterior (rostral→caudal). Scale arrows: 150 µm. M→L: medial to lateral, V→D: ventral to dorsal.

**Table S1.** Summary of statistical analyses

<table>
<tr><th>Figure</th><th>Sample size (n)</th><th>Statistical Test</th><th>Values</th></tr>
<tr><td>Fig. 1E</td><td>Prkcd (Label-IT: 5728, Melphalan: 7059)<br>Slc32a1 (Label-IT: 17771, Melphalan: 12252)</td><td>Unpaired student's t-test</td><td>Average integrated spot intensity:<br>Prkcd (Label-IT: 19313; Melphalan: 40550)<br>Slc32a1 (Label-IT: 40565; Melphalan: 55031)<br>Prkcd: p < 0.0001 (two-tailed)<br>Slc32a1: p < 0.0001(two-tailed)</td></tr>
<tr><td>Fig. 1G</td><td>Shown in Fig. 1G</td><td>Unpaired student's t-test</td><td>SNR (Label-IT: 13.97; Melphalan:18.18)<br>p < 0.0001 (two-tailed)</td></tr>
<tr><td>Fig. 1M</td><td>Shown in Fig. 1M</td><td>least-squares regression</td><td>As indicated in Fig. 1M</td></tr>
<tr><td>Fig. 1N</td><td>Shown in Fig. 1N</td><td>Pearson correlation</td><td>As indicated in Fig. 1N</td></tr>
<tr><td>related to Fig. 4E</td><td>Ex-1: 465<br>Ex-2: 640<br>Ex-3: 676<br>Ex-4: 160<br>Ex-5: 407<br>Ex-6: 432<br>Ex-7: 850<br>Ex-8: 372<br>Ex-9: 118<br>Ex-10: 174<br>Ex-11: 279<br>Ex-12: 1059<br>Ex-13: 821<br>Ex-14: 2433<br>Ex-15: 306<br>Ex-16: 156<br>Ex-17: 182<br>Ex-18: 538<br>Ex-19: 1291<br>Ex-20: 288<br>Ex-21: 138<br>Ex-22: 1003<br>Ex-23: 126<br>Ex-24: 693<br>Ex-25: 2787<br>Inh-1: 81<br>Inh-2: 243<br>Inh-3: 88<br>Inh-4: 1238<br>Inh-5: 332<br>Inh-6: 69<br>Inh-7: 184<br>Inh-8: 844<br>Inh-9: 282<br>Inh-10: 964<br>Inh-11: 648</td><td>Complete Spatial Randomness test with Monte Carlo method<br>(see Method)</td><td>
<table>
<tr><th></th><th>p-value</th><th>$ANN_{Mol}$ -$ANN_{CSR}$</th></tr>
<tr><td>Ex-1</td><td>0</td><td>-12.008419</td></tr>
<tr><td>Ex-2</td><td>0</td><td>-9.315098</td></tr>
<tr><td>Ex-3</td><td>0</td><td>-4.555526</td></tr>
<tr><td>Ex-4</td><td>0</td><td>-29.722482</td></tr>
<tr><td>Ex-5</td><td>0</td><td>-9.056687</td></tr>
<tr><td>Ex-6</td><td>0</td><td>-11.916794</td></tr>
<tr><td>Ex-7</td><td>0</td><td>-8.014789</td></tr>
<tr><td>Ex-8</td><td>0</td><td>-10.062707</td></tr>
<tr><td>Ex-9</td><td>0</td><td>-11.625993</td></tr>
<tr><td>Ex-10</td><td>0</td><td>-3.12223</td></tr>
<tr><td>Ex-11</td><td>0</td><td>-5.905959</td></tr>
<tr><td>Ex-12</td><td>0</td><td>-10.38412</td></tr>
<tr><td>Ex-13</td><td>0</td><td>-9.148289</td></tr>
<tr><td>Ex-14</td><td>0</td><td>-4.624816</td></tr>
<tr><td>Ex-15</td><td>0</td><td>-13.51638</td></tr>
<tr><td>Ex-16</td><td>0</td><td>-17.465758</td></tr>
<tr><td>Ex-17</td><td>0</td><td>-9.69543</td></tr>
<tr><td>Ex-18</td><td>0</td><td>-7.549514</td></tr>
<tr><td>Ex-19</td><td>0</td><td>-10.723788</td></tr>
<tr><td>Ex-20</td><td>0</td><td>-14.344153</td></tr>
<tr><td>Ex-21</td><td>0</td><td>-3.635274</td></tr>
<tr><td>Ex-22</td><td>0</td><td>-8.749299</td></tr>
<tr><td>Ex-23</td><td>0</td><td>-19.049987</td></tr>
<tr><td>Ex-24</td><td>0</td><td>-2.261705</td></tr>
<tr><td>Ex-25</td><td>0</td><td>-0.850436</td></tr>
<tr><td>Inh-1</td><td>0</td><td>-31.127834</td></tr>
<tr><td>Inh-2</td><td>0</td><td>-20.546199</td></tr>
<tr><td>Inh-3</td><td>0</td><td>-45.60516</td></tr>
<tr><td>Inh-4</td><td>0</td><td>-17.039251</td></tr>
<tr><td>Inh-5</td><td>0</td><td>-4.176389</td></tr>
<tr><td>Inh-6</td><td>0</td><td>-20.811941</td></tr>
<tr><td>Inh-7</td><td>0</td><td>-20.32339</td></tr>
<tr><td>Inh-8</td><td>0</td><td>-5.392525</td></tr>
<tr><td>Inh-9</td><td>0</td><td>-13.006909</td></tr>
</table>
</td></tr>
</table>

<table>
<tr>
<td></td>
<td>Inh-12: 875<br>Inh-13: 1307<br>Inh-14: 721<br>Inh-15: 589<br>Inh-16: 645<br>Inh-17: 284<br>Inh-18: 910<br>Inh-19: 2182<br>Inh-20: 425<br>Inh-21: 1158<br>Inh-22: 926<br>Inh-23: 5034</td>
<td></td>
<td><table>
<tr><td>Inh-10</td><td>0</td><td>-6.607319</td></tr>
<tr><td>Inh-11</td><td>0</td><td>-5.476935</td></tr>
<tr><td>Inh-12</td><td>0</td><td>-13.600546</td></tr>
<tr><td>Inh-13</td><td>0</td><td>-8.664817</td></tr>
<tr><td>Inh-14</td><td>0</td><td>-11.434014</td></tr>
<tr><td>Inh-15</td><td>0</td><td>-13.229296</td></tr>
<tr><td>Inh-16</td><td>0</td><td>-7.184522</td></tr>
<tr><td>Inh-17</td><td>0</td><td>-15.423329</td></tr>
<tr><td>Inh-18</td><td>0</td><td>-3.86001</td></tr>
<tr><td>Inh-19</td><td>0</td><td>-6.832312</td></tr>
<tr><td>Inh-20</td><td>0</td><td>-7.189582</td></tr>
<tr><td>Inh-21</td><td>0</td><td>-9.744092</td></tr>
<tr><td>Inh-22</td><td>0</td><td>-4.753114</td></tr>
<tr><td>Inh-23</td><td>0</td><td>-2.60416</td></tr>
</table></td>
</tr>
<tr>
<td>related to Fig. 4E</td>
<td>same as above</td>
<td>Chi-square statistics (see Method)</td>
<td>Ex-1: p=0.01; *<br>Ex-2: p=0; ****<br>Ex-3: p=0; ****<br>Ex-4: p=0; ****<br>Ex-5: p=0; ****<br>Ex-6: p=0; ****<br>Ex-7: p=0; ****<br>Ex-8: p=0; ****<br>Ex-9: p=0; ****<br>Ex-10: p=0; ****<br>Ex-11: p=0; ****<br>Ex-12: p=0; ****<br>Ex-13: p=0; ****<br>Ex-14: p=0; ****<br>Ex-15: p=0; ****<br>Ex-16: p=0.058; ns<br>Ex-17: p=0; ****<br>Ex-18: p=0; ****<br>Ex-19: p=0; ****<br>Ex-20: p=0; ****<br>Ex-21: p=0.459; ns<br>Ex-22: p=0; ****<br>Ex-23: p=0; ****<br>Ex-24: p=0.001; **<br>Ex-25: p=0; ****<br>Inh-1: p=0; ****<br>Inh-2: p=0; ****<br>Inh-3: p=0; ****<br>Inh-4: p=0; ****<br>Inh-5: p=0.759; ns<br>Inh-6: p=0; ****<br>Inh-7: p=0; ****<br>Inh-8: p=0; ****</td>
</tr>
</table>

| | | | |
|---|---|---|---|
| | | | Inh-9: p=0; ****<br>Inh-10: p=0; ****<br>Inh-11: p=0; ****<br>Inh-12: p=0; ****<br>Inh-13: p=0; ****<br>Inh-14: p=0; ****<br>Inh-15: p=0; ****<br>Inh-16: p=0; ****<br>Inh-17: p=0; ****<br>Inh-18: p=0; ****<br>Inh-19: p=0; ****<br>Inh-20: p=0; ****<br>Inh-21: p=0; ****<br>Inh-22: p=0; ****<br>Inh-23: p=0; **** |
| related to **Fig. 4E** | same as above | Permutation test (see Method) | As indicated in Table S7 |
| **Fig. 6D** | Ex-3: 676<br>Ex-4: 160<br>Ex-8: 372<br>Ex-11: 279 | Unpaired Welch's t-test | Ex-3 vs Ex-4: $p = 2.10 \times 10^{-15}$<br>Ex-3 vs Ex-11: $p = 3.21 \times 10^{-11}$<br>Ex-8 vs Ex-4: $p < 2.2 \times 10^{-16}$<br>Ex-8 vs Ex-11: $p < 2.2 \times 10^{-16}$ |
| **Fig. 6G** | Ex-5: 407<br>Inh-1: 81<br>Inh-2: 243<br>Inh-5: 332 | Soma volume: Unpaired Welch's t-test;<br>Solidity:Wilcoxon signed rank test | Soma volume:<br>Ex-5 vs Inh-1: $p = 2.83 \times 10^{-9}$<br>Ex-5 vs Inh-5: $p < 2.2 \times 10^{-16}$<br>Inh-2 vs Inh-1: $p = 3.446 \times 10^{-7}$<br>Inh-2 vs Inh-5: $p < 2.2 \times 10^{-16}$<br>Solidity (morphology):<br>Ex-5 vs Inh-1: $p = 8.12 \times 10^{-11}$<br>Ex-5 vs Inh-2: $p < 2.2 \times 10^{-16}$<br>Ex-5 vs Inh-5: $p < 2.2 \times 10^{-16}$ |
| **Fig. 7A** (Soma volume) | Ex-1: 465<br>Ex-2: 640<br>Ex-3: 676<br>Ex-4: 160<br>Ex-5: 407<br>Ex-6: 432<br>Ex-7: 850<br>Ex-8: 372<br>Ex-9: 118<br>Ex-10: 174<br>Ex-11: 279<br>Ex-12: 1059<br>Ex-13: 821<br>Ex-14: 2433<br>Ex-15: 306<br>Ex-16: 156<br>Ex-17: 182<br>Ex-18: 538<br>Ex-19: 1291 | Welch's one-way ANOVA (parametric without assumption of equal variance) followed by unpaired Welch's t-test compared to the population mean with Benjamini, Hochberg, and Yekutieli false discovery rate adjustment | Average *Slc17a6*$^+$ cell body volume: 1771.258μm$^3$, ANOVA, p=0, ****<br>Average soma volume in each molecular cluster:<br>Ex-1: 3642.133 μm$^3$, $p=2.00 \times 10^{-92}$, ****<br>Ex-2: 3690.510 μm$^3$, $p=3.10 \times 10^{-212}$, ****<br>Ex-3: 2191.752 μm$^3$, $p=3.30 \times 10^{-53}$, ****<br>Ex-4: 1802.396 μm$^3$, p=0.44, ns<br>Ex-5: 1951.366 μm$^3$, $p=1.30 \times 10^{-10}$, ****<br>Ex-6: 1967.923 μm$^3$, $p=1.30 \times 10^{-14}$, ****<br>Ex-7: 1952.845 μm$^3$, $p=5.40 \times 10^{-16}$, ****<br>Ex-8: 2337.702 μm$^3$, $p=3.60 \times 10^{-42}$, ****<br>Ex-9: 1975.957 μm$^3$, $p=9.20 \times 10^{-4}$, ***<br>Ex-10: 2262.927 μm$^3$, $p=7.90 \times 10^{-9}$, ****<br>Ex-11: 1894.662 μm$^3$, $p=1.10 \times 10^{-3}$, ***<br>Ex-12: 1410.385 μm$^3$, $p=2.70 \times 10^{-159}$, ****<br>Ex-13: 1741.036 μm$^3$, $p=1.00 \times 10^{-1}$, ns<br>Ex-14: 1753.767 μm$^3$, $p=1.80 \times 10^{-1}$, ns<br>Ex-15: 1586.896 μm$^3$, $p=1.10 \times 10^{-12}$, **** |

| | | | |
|---|---|---|---|
| | Ex-20: 288<br>Ex-21: 138<br>Ex-22: 1003<br>Ex-23: 126<br>Ex-24: 693<br>Ex-25: 2787 | | Ex-16: 1533.906 $\mu m^3$, p=3.00 × $10^{-10}$, ****<br>Ex-17: 1475.152 $\mu m^3$, p=5.70 × $10^{-20}$, ****<br>Ex-18: 1553.394 $\mu m^3$, p=2.80 × $10^{-16}$, ****<br>Ex-19: 1423.846 $\mu m^3$, p=2.00 × $10^{-183}$, ****<br>Ex-20: 1382.159 $\mu m^3$, p=7.90 × $10^{-53}$, ****<br>Ex-21: 1539.889 $\mu m^3$, p=3.50 × $10^{-11}$, ****<br>Ex-22: 1531.342 $\mu m^3$, p=7.40 × $10^{-58}$, ****<br>Ex-23: 1427.399 $\mu m^3$, p=8.10 × $10^{-28}$, ****<br>Ex-24: 1422.796 $\mu m^3$, p=2.10 × $10^{-67}$, ****<br>Ex-25: 1332.447 $\mu m^3$, p=8.50 × $10^{-229}$, **** |
| **Fig. 7A**<br>(Solidity) | same as above | Kruskal-Wallis rank sum test (non-parametric without assumption of equal variance), Wilcoxon rank sum test compared to the population mean with Benjamini, Hochberg, and Yekutieli false discovery rate adjustment, Two-sample Kolmogorov–Smirnov test compared to *Slc17a6*$^+$ population distribution | Kruskal test: p=0, ****<br>p-values:<br>Ex-1: Wilcox: 2.90 × $10^{-56}$, KS: < 2.2 × $10^{-16}$<br>Ex-2: Wilcox: 5.40 × $10^{-46}$, KS: < 2.2 × $10^{-16}$<br>Ex-3: Wilcox: 7.70 × $10^{-25}$, KS: < 2.2 × $10^{-16}$<br>Ex-4: Wilcox: 2.10 × $10^{-14}$, KS: 4.34 × $10^{-11}$<br>Ex-5: Wilcox: 9.60 × $10^{-23}$, KS: < 2.2 × $10^{-16}$<br>Ex-6: Wilcox: 1.40 × $10^{-14}$, KS: 5.84 × $10^{-14}$<br>Ex-7: Wilcox: 3.10 × $10^{-20}$, KS: < 2.2 × $10^{-16}$<br>Ex-8: Wilcox: 8.00 × $10^{-4}$, KS: 1.20 × $10^{-3}$<br>Ex-9: Wilcox: 1.50 × $10^{-1}$, KS: 1.50 × $10^{-1}$<br>Ex-10: Wilcox: 1.80 × $10^{-1}$, KS: 8.10 × $10^{-2}$<br>Ex-11: Wilcox: 1.40 × $10^{-1}$, KS: 3.62 × $10^{-2}$<br>Ex-12: Wilcox: 1.90 × $10^{-47}$, KS: < 2.2 × $10^{-16}$<br>Ex-13: Wilcox: 4.80 × $10^{-21}$, KS: 4.80 × $10^{-15}$<br>Ex-14: Wilcox: 5.60 × $10^{-20}$, KS: < 2.2 × $10^{-16}$<br>Ex-15: Wilcox: 3.50 × $10^{-2}$, KS: 3.30 × $10^{-3}$<br>Ex-16: Wilcox: 1.10 × $10^{-3}$, KS: 3.06 × $10^{-2}$<br>Ex-17: Wilcox: 1.90 × $10^{-2}$, KS: 5.13 × $10^{-2}$<br>Ex-18: Wilcox: 5.60 × $10^{-1}$, KS: 2.78 × $10^{-1}$<br>Ex-19: Wilcox: 2.40 × $10^{-52}$, KS: < 2.2 × $10^{-16}$<br>Ex-20: Wilcox: 2.10 × $10^{-10}$, KS: 3.23 × $10^{-9}$<br>Ex-21: Wilcox: 4.00 × $10^{-2}$, KS: 1.46 × $10^{-1}$<br>Ex-22: Wilcox: 3.40 × $10^{-4}$, KS: 1.27 × $10^{-6}$<br>Ex-23: Wilcox: 1.50 × $10^{-6}$, KS: 1.40 × $10^{-4}$<br>Ex-24: Wilcox: 3.10 × $10^{-1}$, KS: 1.67 × $10^{-2}$<br>Ex-25: Wilcox: 5.30 × $10^{-16}$, KS: 7.96 × $10^{-12}$ |
| **Fig. 7A**<br>(Aspect Ratio) | same as above | Kruskal-Wallis rank sum test (non-parametric without assumption of equal variance), Wilcoxon rank sum test compared to the population mean with Benjamini, Hochberg, and Yekutieli false discovery rate adjustment, | Kruskal test: p= 2.8 × $10^{-94}$, ****<br>p-values:<br>Ex-1: Wilcox: 2.50 × $10^{-2}$, KS: 8.69 × $10^{-3}$<br>Ex-2: Wilcox: 1.50 × $10^{-3}$, KS: 1.14 × $10^{-2}$<br>Ex-3: Wilcox: 8.40 × $10^{-11}$, KS: 3.78 × $10^{-11}$<br>Ex-4: Wilcox: 3.30 × $10^{-6}$, KS: 7.24 × $10^{-5}$<br>Ex-5: Wilcox: 2.50 × $10^{-18}$, KS: 3.22 × $10^{-15}$<br>Ex-6: Wilcox: 1.20 × $10^{-6}$, KS: 8.28 × $10^{-6}$<br>Ex-7: Wilcox: 1.40 × $10^{-18}$, KS: 1.02 × $10^{-14}$<br>Ex-8: Wilcox: 5.60 × $10^{-2}$, KS: 1.81× $10^{-2}$<br>Ex-9: Wilcox: 5.70 × $10^{-2}$, KS: 1.66× $10^{-1}$<br>Ex-10: Wilcox: 5.20 × $10^{-2}$, KS: 5.91 × $10^{-2}$<br>Ex-11: Wilcox: 5.90 × $10^{-1}$, KS: 6.46 × $10^{-1}$<br>Ex-12: Wilcox: 2.10 × $10^{-6}$, KS: 5.24 × $10^{-5}$ |

| | | | |
|---|---|---|---|
| | | Two-sample Kolmogorov–Smirnov test compared to *Slc17a6*$^+$ population distribution | Ex-13: Wilcox: $2.10 \times 10^{-1}$, KS: $9.66 \times 10^{-3}$<br>Ex-14: Wilcox: $1.50 \times 10^{-1}$, KS: $2.59 \times 10^{-1}$<br>Ex-15: Wilcox: $5.70 \times 10^{-2}$, KS: $1.07 \times 10^{-1}$<br>Ex-16: Wilcox: $6.20 \times 10^{-2}$, KS: $8.60 \times 10^{-2}$<br>Ex-17: Wilcox: $3.30 \times 10^{-1}$, KS: $2.86 \times 10^{-1}$<br>Ex-18: Wilcox: $1.70 \times 10^{-1}$, KS: $4.72 \times 10^{-1}$<br>Ex-19: Wilcox: $2.00 \times 10^{-20}$, KS: $4.44 \times 10^{-16}$<br>Ex-20: Wilcox: $7.70 \times 10^{-5}$, KS: $3.90 \times 10^{-4}$<br>Ex-21: Wilcox: $2.10 \times 10^{-1}$, KS: $3.33 \times 10^{-1}$<br>Ex-22: Wilcox: $5.90 \times 10^{-1}$, KS: $2.02 \times 10^{-2}$<br>Ex-23: Wilcox: $3.10 \times 10^{-3}$, KS: $1.24 \times 10^{-2}$<br>Ex-24: Wilcox: $1.00 \times 10^{-8}$, KS: $7.66 \times 10^{-9}$<br>Ex-25: Wilcox: $2.40 \times 10^{-2}$, KS: $4.24 \times 10^{-3}$ |
| **Fig. 7B** (Soma volume) | Inh-1: 81<br>Inh-2: 243<br>Inh-3: 88<br>Inh-4: 1238<br>Inh-5: 332<br>Inh-6: 69<br>Inh-7: 184<br>Inh-8: 844<br>Inh-9: 282<br>Inh-10: 964<br>Inh-11: 648<br>Inh-12: 875<br>Inh-13: 1307<br>Inh-14: 721<br>Inh-15: 589<br>Inh-16: 645<br>Inh-17: 284<br>Inh-18: 910<br>Inh-19: 2182<br>Inh-20: 425<br>Inh-21: 1158<br>Inh-22: 926<br>Inh-23: 5034 | Welch's one-way ANOVA (parametric without assumption of equal variance) followed by *post hoc* multiple comparison to the population mean with Benjamini, Hochberg, and Yekutieli false discovery rate adjustment | Average *Slc32a1*$^+$soma volume: 1403.199μm$^3$, ANOVA, p=0, ****<br>Average soma volume in each molecular cluster:<br>Inh-1: 1641.356 μm$^3$, p=$2.40 \times 10^{-7}$, ****<br>Inh-2: 1896.634 μm$^3$, area$6.90 \times 10^{-56}$, ****<br>Inh-3: 1933.433 μm$^3$, $1.60 \times 10^{-18}$, ****<br>Inh-4: 1504.277 μm$^3$, $1.60 \times 10^{-17}$, ****<br>Inh-5: 1611.615 μm$^3$, $1.80 \times 10^{-30}$, ****<br>Inh-6: 1862.899 μm$^3$, $3.00 \times 10^{-9}$, ****<br>Inh-7: 1841.264 μm$^3$, $5.00 \times 10^{-35}$, ****<br>Inh-8: 1783.614 μm$^3$, $1.00 \times 10^{-87}$, ****<br>Inh-9: 1398.871 μm$^3$, $8.20 \times 10^{-1}$, ns<br>Inh-10: 1554.772 μm$^3$, $2.20 \times 10^{-18}$, ****<br>Inh-11: 1450.228 μm$^3$, $1.80 \times 10^{-2}$, *<br>Inh-12: 1435.796 μm$^3$, $4.30 \times 10^{-3}$, **<br>Inh-13: 1270.102 μm$^3$, $3.50 \times 10^{-39}$, ****<br>Inh-14: 1443.659 μm$^3$, $1.60 \times 10^{-2}$, *<br>Inh-15: 1409.52 μm$^3$, $7.40 \times 10^{-1}$, ns<br>Inh-16: 1480.219 μm$^3$, $9.40 \times 10^{-7}$, ****<br>Inh-17: 1437.867 μm$^3$, $1.40 \times 10^{-1}$, ns<br>Inh-18: 1479.489 μm$^3$, $1.10 \times 10^{-7}$, ****<br>Inh-19: 1320.613 μm$^3$, $2.50 \times 10^{-26}$, ****<br>Inh-20: 1501.225 μm$^3$, $9.10 \times 10^{-9}$, ****<br>Inh-21: 1313.432 μm$^3$, $1.40 \times 10^{-11}$, ****<br>Inh-22: 1294.191 μm$^3$, $3.30 \times 10^{-19}$, ****<br>Inh-23: 1271.708 μm$^3$, $3.00 \times 10^{-99}$, **** |
| Fig. 7B (Solidity) | same as above | Kruskal-Wallis rank sum test (non-parametric without assumption of equal variance), Wilcoxon rank sum test compared to the population mean with Benjamini, Hochberg, and Yekutieli false | Kruskal test: p=$9 \times 10^{-96}$, ****<br>p-values:<br>Inh-1: Wilcox: $9.40 \times 10^{-1}$; KS: $8.06 \times 10^{-1}$<br>Inh-2: Wilcox: $9.00 \times 10^{-1}$; KS: $9.95\times 10^{-1}$<br>Inh-3: Wilcox: $7.60 \times 10^{-1}$; KS: $4.85\times 10^{-1}$<br>Inh-4: Wilcox: $3.40 \times 10^{-17}$; KS: $1.19\times 10^{-14}$<br>Inh-5: Wilcox: $2.30 \times 10^{-7}$; KS: $2.14\times 10^{-6}$<br>Inh-6: Wilcox: $4.30 \times 10^{-5}$; KS: $1.07\times 10^{-4}$<br>Inh-7: Wilcox: $1.70 \times 10^{-1}$; KS: $1.64\times 10^{-1}$<br>Inh-8: Wilcox: $4.00 \times 10^{-9}$; KS: $3.22\times 10^{-7}$<br>Inh-9: Wilcox: $2.10 \times 10^{-1}$; KS: $2.04\times 10^{-1}$ |

| | | | |
|---|---|---|---|
| | | discovery rate adjustment, Two-sample Kolmogorov–Smirnov test compared to Slc17a6+ population distribution | Inh-10: Wilcox: 4.90× $10^{-2}$; KS: 8.24× $10^{-2}$<br>Inh-11: Wilcox: 2.60× $10^{-18}$; KS: 1.55× $10^{-14}$<br>Inh-12: Wilcox: 4.90× $10^{-2}$; KS: 6.64× $10^{-2}$<br>Inh-13: Wilcox: 5.40× $10^{-4}$; KS: 5.83× $10^{-3}$<br>Inh-14: Wilcox: 5.20× $10^{-1}$; KS: 4.15× $10^{-1}$<br>Inh-15: Wilcox: 4.40× $10^{-3}$; KS: 6.05× $10^{-5}$<br>Inh-16: Wilcox: 5.90× $10^{-1}$; KS: 7.18× $10^{-1}$<br>Inh-17: Wilcox: 4.90× $10^{-1}$; KS: 1.52× $10^{-2}$<br>Inh-18: Wilcox: 1.20× $10^{-10}$; KS: 1.26× $10^{-10}$<br>Inh-19: Wilcox: 7.60× $10^{-10}$; KS: 9.52× $10^{-9}$<br>Inh-20: Wilcox: 1.50× $10^{-2}$; KS: 1.42× $10^{-2}$<br>Inh-21: Wilcox: 1.70× $10^{-19}$; KS: < 2.2× $10^{-16}$<br>Inh-22: Wilcox: 3.60× $10^{-6}$; KS: 6.47× $10^{-7}$<br>Inh-23: Wilcox: 5.20× $10^{-3}$; KS: 1.43× $10^{-3}$ |
| **Fig. 7B** (Aspect Ratio) | same as above | Kruskal-Wallis rank sum test (non-parametric without assumption of equal variance), Wilcoxon rank sum test compared to the population mean with Benjamini, Hochberg, and Yekutieli false discovery rate adjustment, Two-sample Kolmogorov–Smirnov test compared to Slc17a6+ population distribution | Kruskal test: p=4.80 × $10^{-36}$, ****<br>p-values:<br>Inh-1: Wilcox: 8.50 × $10^{-1}$; KS: 7.52 × $10^{-1}$<br>Inh-2: Wilcox: 6.10 × $10^{-2}$; KS: 2.53 × $10^{-2}$<br>Inh-3: Wilcox: 7.60 × $10^{-2}$; KS: 1.99 × $10^{-1}$<br>Inh-4: Wilcox: 7.50 × $10^{-2}$; KS: 6.08 × $10^{-2}$<br>Inh-5: Wilcox: 2.30 × $10^{-1}$; KS: 2.35 × $10^{-1}$<br>Inh-6: Wilcox: 5.20 × $10^{-2}$; KS: 3.49 × $10^{-2}$<br>Inh-7: Wilcox: 2.60 × $10^{-1}$; KS: 9.63 × $10^{-2}$<br>Inh-8: Wilcox: 2.40 × $10^{-3}$; KS: 2.62 × $10^{-3}$<br>Inh-9: Wilcox: 3.80 × $10^{-1}$; KS: 1.97 × $10^{-1}$<br>Inh-10: Wilcox: 9.90 × $10^{-1}$; KS: 8.50 × $10^{-1}$<br>Inh-11: Wilcox: 2.30 × $10^{-1}$; KS: 1.45 × $10^{-1}$<br>Inh-12: Wilcox: 7.50 × $10^{-2}$; KS: 9.25 × $10^{-2}$<br>Inh-13: Wilcox: 3.80 × $10^{-2}$; KS: 7.76 × $10^{-3}$<br>Inh-14: Wilcox: 9.90 × $10^{-1}$; KS: 5.12 × $10^{-1}$<br>Inh-15: Wilcox: 2.30 × $10^{-1}$; KS: 2.22 × $10^{-1}$<br>Inh-16: Wilcox: 7.60 × $10^{-1}$; KS: 8.16 × $10^{-1}$<br>Inh-17: Wilcox: 1.80 × $10^{-1}$; KS: 1.11 × $10^{-1}$<br>Inh-18: Wilcox: 9.60 × $10^{-9}$; KS: 3.44 × $10^{-7}$<br>Inh-19: Wilcox: 6.10 × $10^{-12}$; KS: 7.95 × $10^{-10}$<br>Inh-20: Wilcox: 2.30 × $10^{-1}$; KS: 3.55 × $10^{-1}$<br>Inh-21: Wilcox: 1.70 × $10^{-13}$; KS: 1.67 × $10^{-11}$<br>Inh-22: Wilcox: 8.50 × $10^{-1}$; KS: 8.10 × $10^{-1}$<br>Inh-23: Wilcox: 6.10 × $10^{-2}$; KS: 4.20 × $10^{-2}$ |
| Fig. S1A | Shown in Fig.S1A | Unpaired student's t-test | *Ezr:* p=0.59, ns<br>*Prkcd:* p=0.48, ns<br>*Slc32a1:* p=0.65, ns |
| Fig. S1F | Number of samples shown in Fig.S1F | Unpaired student's t-test | p=0.0001 |
| Fig. S1L, S1M | Number of samples shown | least-square linear regression | as indicated in Fig. S1L and S1M |

| | | | |
|---|---|---|---|
| Fig. S2F | Number of samples shown in Fig.S2F | least-square linear regression | As indicated in Fig. S2F |
| Fig. S2G | Number of genes shown in Fig.S2G | power-law fit | As indicated in Fig. S2G<br>$p=9.48 \times 10^{-15}$ |
| Fig. S4F-G | Shown in Fig.S4F-G | Pairwise Pearson correlation | As indicated in Fig. S4F-G |
| Fig. S5E | Shown in Fig.S5E | Pairwise Pearson correlation | As indicated in Fig. S5E |
| Fig. S5G | Shown in Fig.S5G | Pairwise Pearson correlation | As indicated in Fig. S5G |
| Fig. S7D | Shown in Fig.S7D | Pairwise Pearson correlation | p=0 |
| Fig. S7E | Shown in Fig.S7E | Pairwise Pearson correlation | p=0 |

**Table S2**. Anti-fade reduces non-stationary HCR spots induced by laser exposure

| | | reduce wigglers | induce AF546 photo-bleaching |
|---|---|---|---|
| Oxygen scavengers | Bubble argon gas | No | No |
| | Glucose oxidase/Catalase | No | Yes |
| | Oxyrase | No | Yes |
| Anti-fade reagents | PPD | Yes | Yes |
| | DABCO | Yes | Yes |
| | n-propyl-gallate | Yes | Yes |
| | Trolox | Yes | Yes |
| | COT | No | N/A |
| | Ascorbic acid | N/A | Yes |

**Table S3**. Segmentation methods evaluation (n=297 neurons)

| | Ilastik+watershed | Stardist 3D | Starfinity |
|---|---|---|---|
| under-segmentation (cell-cell merge) | 9.40% | 1% | 1% |
| under detection | 15.80% | 0.30% | 0.30% |
| over-segmentation | 7% | 2% | 2% |
| contamination errors | 6.70% | 3% | 2% |

**Table S4**. Differentially expressed genes in molecularly defined clusters identified from scRNA-Seq data (provided as a separate .xls file).

Gene names are listed in the first column, and molecular clusters in second column. p_val_adj: Adjusted p-value is based on the Bonferroni correction; avg_logFC: log fold-chage of the average expression between cells in the cluster of interest and all other cells; pct.1: The percentage of cells where the feature is detected in the cluster of interest; pct.2: The percentage of cells where the feature is detected in all other cells.

**Table S5**. Marker-genes used for EASI-FISH experiment in the LHA

| | Genes | HCR hairpin | Fluorophores |
|---|---|---|---|
| Round 1 | *Meis2* | B1 | AF-488 |
| | *Th* | B2 | JF-699 |
| | *Gpr101* | B5 | AF-546 |
| Round 2 | *Slc17a6* | B1 | JF-699 |
| | *Gpr83* | B2 | AF-488 |
| | *Bdnf* | B5 | AF-546 |
| Round 3 | *Otp* | B1 | AF-546 |
| | *Calb2* | B2 | AF-488 |
| | *Gal* | B5 | JF-699 |
| Round 4 | *Col25a1* | B1 | JF-699 |
| | *Synpr* | B2 | AF-546 |
| | *Gad1* | B3 | AF-488 |
| Round 5 | *Tac1* | B1 | AF-546 |
| | *Tac2* | B2 | AF-488 |
| | *Calb1* | B5 | JF-699 |
| Round 6 | *Trh* | B1 | JF-699 |
| | *Map1b* | B2 | AF-488 |
| | *Nts* | B5 | AF-546 |
| Round 7 | *Pmch* | B1 | JF-699 |
| | *Cartpt* | B2 | AF-546 |
| | *Hcrt* | B5 | AF-488 |
| Round 8 | *Slc32a1* | B1 | AF-488 |
| | *Nrgn* | B2 | AF-546 |
| | *Sst* | B5 | JF-699 |
| Round 9 | *Meis2* | B1 | AF-488 |
| | *Th* | B2 | JF-699 |
| | *Gpr101* | B5 | AF-546 |

**Table S6**. Cellular of EASI-FISH experiments in the LHA from three animals

| | LHA1 | LHA2 | LHA3 | total |
|---|---|---|---|---|
| # of cells in the tissue volume | 27,199 | 28,714 | 30,069 | 85,982 |
| # of cells included in the analysis | 19,694 | 21,912 | 24,882 | 66,488 |
| # of neurons | 10,442 | 12,430 | 13,551 | 36,423 |
| # of *Slc17a6*$^+$ neurons | 4,954 | 7,129 | 7,946 | 16,394 |
| # of *Slc32a1*$^+$ neurons | 5,488 | 5,301 | 5,605 | 20,029 |

**Table S7**. Molecular cell type enrichment in LHA subregions
(Permutation test, p-values (<0.05 are shown) corresponding to **Figure 4E**)

| | EP | LHAfl | LHAs-db | LHAfm | LHAdl | LHAhcrt-db | LHAd-db | ZI |
|---|---|---|---|---|---|---|---|---|
| Ex-12 | 0 | | 0 | | | | | |
| Ex-14 | 0 | | 0 | | 0 | | | |
| Ex-23 | 0 | | 0.03 | | 0.005 | | | |
| Ex-18 | 0 | | 0 | | 0 | | | |
| Ex-20 | 0 | | 0 | | 0 | | | |
| Ex-24 | | 0 | 0 | | | | | |
| Inh-5 | | | 0 | 0.038 | | | | |
| Ex-6 | | 0 | 0.01 | 0 | | | | |
| Ex-4 | | 0 | | 0.027 | | | | |
| Ex-7 | | 0 | 0 | 0 | | | | |
| Ex-11 | | 0 | 0 | 0.001 | | | | |
| Ex-25 | | 0 | 0 | 0 | | | | |
| Inh-11 | | 0 | 0 | | | | | |
| Ex-8 | | 0 | 0 | | | | | |
| Ex-5 | | 0 | 0 | 0.043 | | | | |
| Inh-7 | | 0 | 0 | | | | | |
| Inh-18 | | | 0 | 0 | | | | |
| Ex-17 | | | 0 | 0 | | | | |
| Ex-9 | | | 0 | 0 | | | | |
| Inh-1 | | | 0 | 0 | | | | |
| Ex-3 | | 0 | 0 | 0 | | | | |
| Inh-8 | | 0 | 0 | 0 | | | | |
| Inh-22 | | 0 | 0 | 0 | | | | |
| Ex-1 | | | 0 | 0 | | | | |
| Inh-23 | | | 0 | 0 | | | 0.001 | |
| Ex-21 | | | 0 | 0.031 | | | | |
| Ex-16 | | | 0 | | | | 0 | |
| Ex-15 | | | 0 | | | | 0 | |
| Ex-10 | | | 0 | | | | 0 | |
| Ex-22 | | | 0 | | | | 0 | |
| Ex-13 | | | 0 | | | | 0 | |
| Ex-19 | | | 0 | | | | 0 | |
| Inh-10 | | | 0 | 0 | | | 0 | |
| Inh-4 | | | 0 | | | | 0 | |
| Inh-16 | | | 0 | | | | 0 | |
| Inh-3 | | | | | | | | 0 |
| Inh-15 | | | 0 | | | | 0 | 0 |
| Inh-21 | | | 0 | | | | | 0 |
| Ex-2 | | | 0 | | 0.026 | 0 | | |
| Inh-14 | | | 0 | | | 0 | | |
| Inh-20 | | | 0 | | | | | |
| Inh-12 | | | 0 | | | | | |
| Inh-2 | | | 0 | | | | | |
| Inh-6 | | | 0 | | | | | |
| Inh-17 | | | 0 | | | | | |
| Inh-19 | | | 0 | | | | | 0 |
| Inh-9 | | | 0 | | | | | |
| Inh-13 | | | 0 | | | | | 0 |
