## Supplemental Information_movie for "Expansion-Assisted Iterative-FISH defines lateral hypothalamus spatio-molecular organization"

### Slide 1
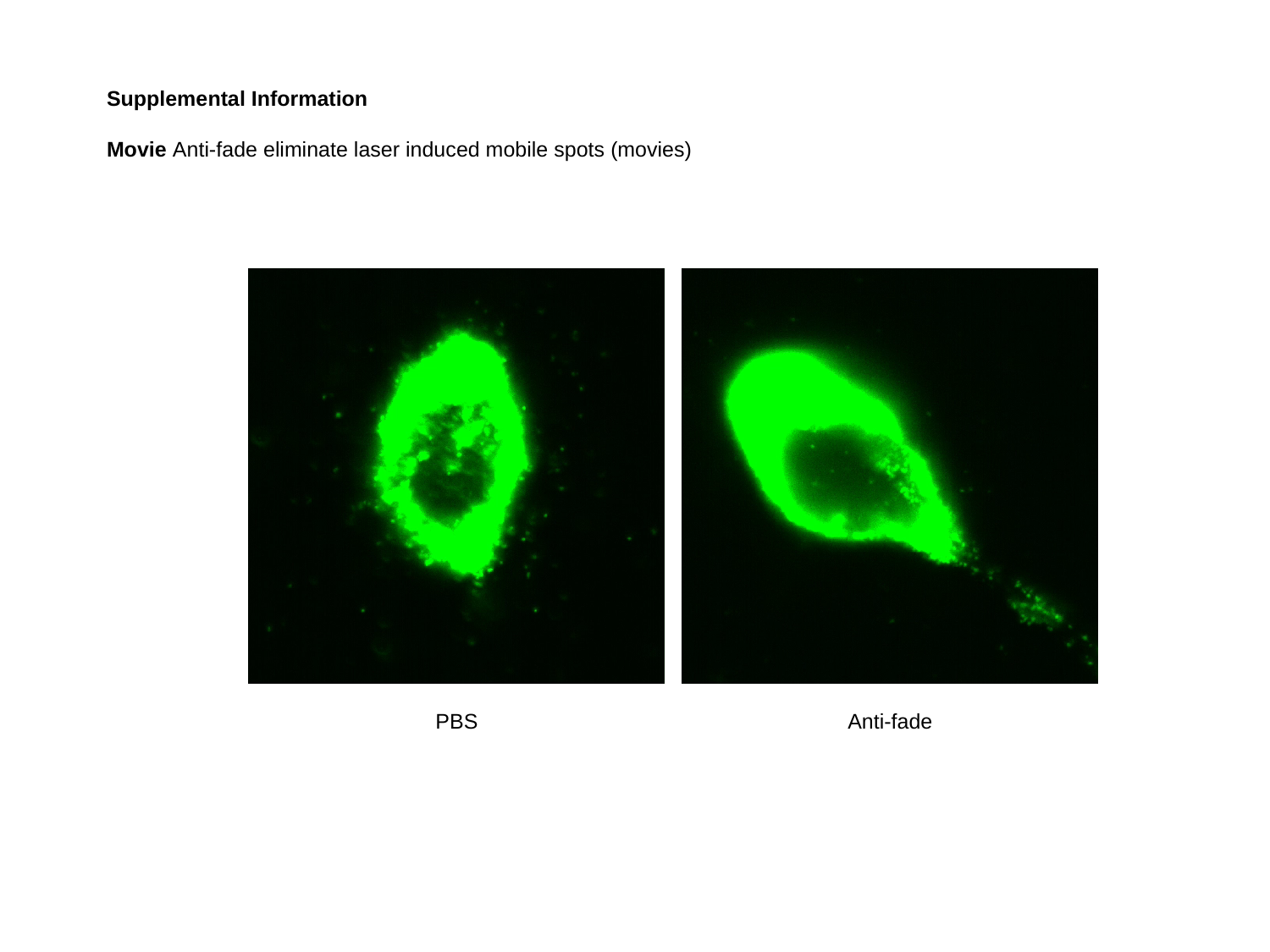

Supplemental Information
Movie Anti-fade eliminate laser induced mobile spots (movies)
PBS
Anti-fade
